# Accumulation of per- and polyfluoroalkyl substances (PFAS) in a terrestrial food web

**DOI:** 10.1101/2023.12.12.571392

**Authors:** Frauke Ecke, Alexandra Skrobonja, Jonas Malmsten, Lutz Ahrens

## Abstract

Per– and polyfluoroalkyl substances (PFAS) are synthetic organofluorine chemical compounds that are broadly used in amongst others aqueous firefighting foam, cosmetics, textiles, carpets, coatings, plastics, and ski wax. Their chemical properties make them persistent organic pollutants that are potential bioaccumulative and toxic. Most studies on PFAS have been performed in groundwater, surface water and aquatic biota. Our knowledge on the terrestrial fate of PFAS is therefore limited.

We sampled soil, berries, mushrooms, and wildlife on the island Frösön, central Sweden, in 2021 and 2022, to study the fate of 22 PFAS in a terrestrial food web. Groundwater, surface water and fish on Frösön have previously shown high PFAS concentrations. Soil, berries, and mushrooms were also concurrently sampled in a reference area in northern Sweden.

Overall, concentrations of the sum of PFAS were low in berries and mushrooms. In moose (*Alces alces*), roedeer (*Capreolus capreolus*), and bank vole (*Myodes glareolus*), concentrations were highest in liver. The maximum levels for PFOS in offal as set by the European Commission (50 ng/g ww) were exceeded in the liver of two of 10 roedeer. Bank voles (*n* = 12 pools) that were sampled in the vicinity of a firefighting training site showed partly extreme concentrations with none of the samples having liver concentrations <474 ng/g ww (maximum 11,600 ng/g ww). Bioaccumulation factors (BAFs) for multiple PFAS in bank voles were higher for the studied mushrooms and soil compared to studied berries and were >100 for 27 out of 265 calculated BAFs. BAFs in the ungulate species were generally lower than those for bank voles but were >1 for several PFAS from the mushroom.

The exact origin of PFAS in bank voles could not be identified in our study, but the BAFs are supported by the feeding and behaviour ecology of bank voles, i.e., there is likely bioaccumulation and biomagnification from soil and mushrooms to bank voles. The measured PFAS concentrations and BAFs, especially those in bank voles are worrying from an ecosystem and One Health perspective considering that voles are staple food for multiple predators.

## Introduction

Per– and polyfluoroalkyl substances (PFAS) include today more than 4700 synthetic organofluorine chemical compounds with hydrophilic, lipophilic and dirt repellent properties (Chelcea et al., 2020). Since their introduction in the 1950s, they have been broadly used in amongst others aqueous firefighting foam, cosmetics, textiles, carpets, coatings, plastics, and ski wax. Their chemical properties make them persistent organic pollutants that are potential bioaccumulative and toxic (Ahrens and Bundschuh, 2014). They are toxic since many PFAS compromise the immune system, are hormone-disrupting, alter reproductive development and are cancerogenic (Dickman and Aga, 2022).

The environmental burden of PFAS has shown high spatial variation. PFAS can be emitted via point sources and locally deposited, while they also have shown long-range water and atmospheric transport or even translocation via migratory wildlife, or a combination of multiple transport pathways. For example, multiple PFAS have been emitted locally at firefighting training sites with further transport of the compounds to recipients and the atmosphere (Ahrens and Bundschuh, 2014; Evich et al., 2022; Koch et al., 2019). These transportation and dispersal mechanisms have contributed to PFAS nowadays being present ubiquitously in the environment.

Most studies on PFAS have been performed in groundwater, surface water and aquatic biota and aquatic food webs (Kurwadkar et al., 2022). Our knowledge on the terrestrial fate of PFAS including occurrence and concentrations along terrestrial food chains and food webs is therefore limited. Recent findings urge us however to fill these gaps in knowledge. Partly extreme high PFAS concentrations have been detected in surface and groundwater and fish close to firefighting training sites (Axelsson and Bard, 2015). These PFAS do not stay in aquatic food webs. Plant-uptake, digestion of PFAS-coated soil particles by invertebrates and uptake of PFAS-rich dust are all mechanisms that can contribute to PFAS also entering the terrestrial food chain with risk of bioaccumulation and biomagnification in consumers and predators. In fact, PFAS have been detected in multiple terrestrial biota including earthworms (reviewed by Burkhard and Votava, 2023), voles (Ecke et al., 2020; Grønnestad et al., 2019) and their predators (Bustnes et al., 2015), reindeer (Roos et al., 2022), roedeer (Falk et al., 2012) and polar bears (Smithwick et al., 2005).

Small mammals are keystone species for the functioning of boreal ecosystems. They consume mushrooms as well as plants and their berries, and contribute to the dispersal of cryptogams and vascular plants (Ericson, 1977; Hansson, 1988). They are also important staple food for multiple mammalian and avian predators (Hörnfeldt et al., 1990; Krebs and Myers, 1974). Their key role in bottom-up and top-down ecosystem processes makes small mammals therefore suitable model species to increase our understanding of the terrestrial fate of PFAS. The bank vole (*Myodes glareolus*) is Europe’s most common mammal (Mitchell-Jones et al., 1999). Even if the species is generally regarded as granivorous and herbivorous, it frequently also consumes mushrooms and invertebrates (Hansson, 1979a; Hansson, 1985b). Due to their limited home ranges (up to ca. 0.4 ha) (Bergstedt, 1966; Löfgren, 1995), micropollutants detected in their tissue likely reflect local exposure (e.g., Ecke et al., 2020; Ecke et al., 2018). Ungulates like roedeer (*Capreolus capreolus*) and moose (*Alces alces*) are browsers mainly feeding on twigs of trees and shrubs with roedeer however also feeding on for example crops (Spitzer et al., 2020). Also roedeer and moose might consume mushrooms and berries (either deliberately or accidentally) (Cederlund et al., 1980). The home range of roedeer is indeed considerably smaller (0.5 km^2^) than that of moose (ca. 72 km^2^) (Jones et al., 2009), but compared to bank voles, it is more difficult to link uptake of pollutants to potential point sources in ungulates.

In previous studies of PFAS on the island Frösön at lake Storsjön, central Sweden, partly extreme PFAS concentrations were detected in groundwater (>400,000 ng/L) (Axelsson and Bard, 2015), and concentrations in fish (Modin, 2021) have resulted in dietary recommendations. These findings pose the opportunity and need to study the fate of PFAS in the terrestrial food web on this island.

We therefore studied PFAS in soil and biota (mushrooms, berries of vascular plants, voles, and ungulates) with mushrooms, berries, and ungulates representing important complementary local year-round food sources (either fresh or frozen) for human consumption. PFAS accumulate and damage liver tissue (reviewed by Costello et al., 2022). We therefore predicted PFAS concentrations in bank voles, roedeer and moose to be highest in liver. Based on feeding and movement ecology, we hypothesized that bank voles have higher PFAS concentrations than roedeer and moose and that their concentrations are higher than those in soil, berries, and mushrooms (see also Fig. 1).

**Figure 1.**
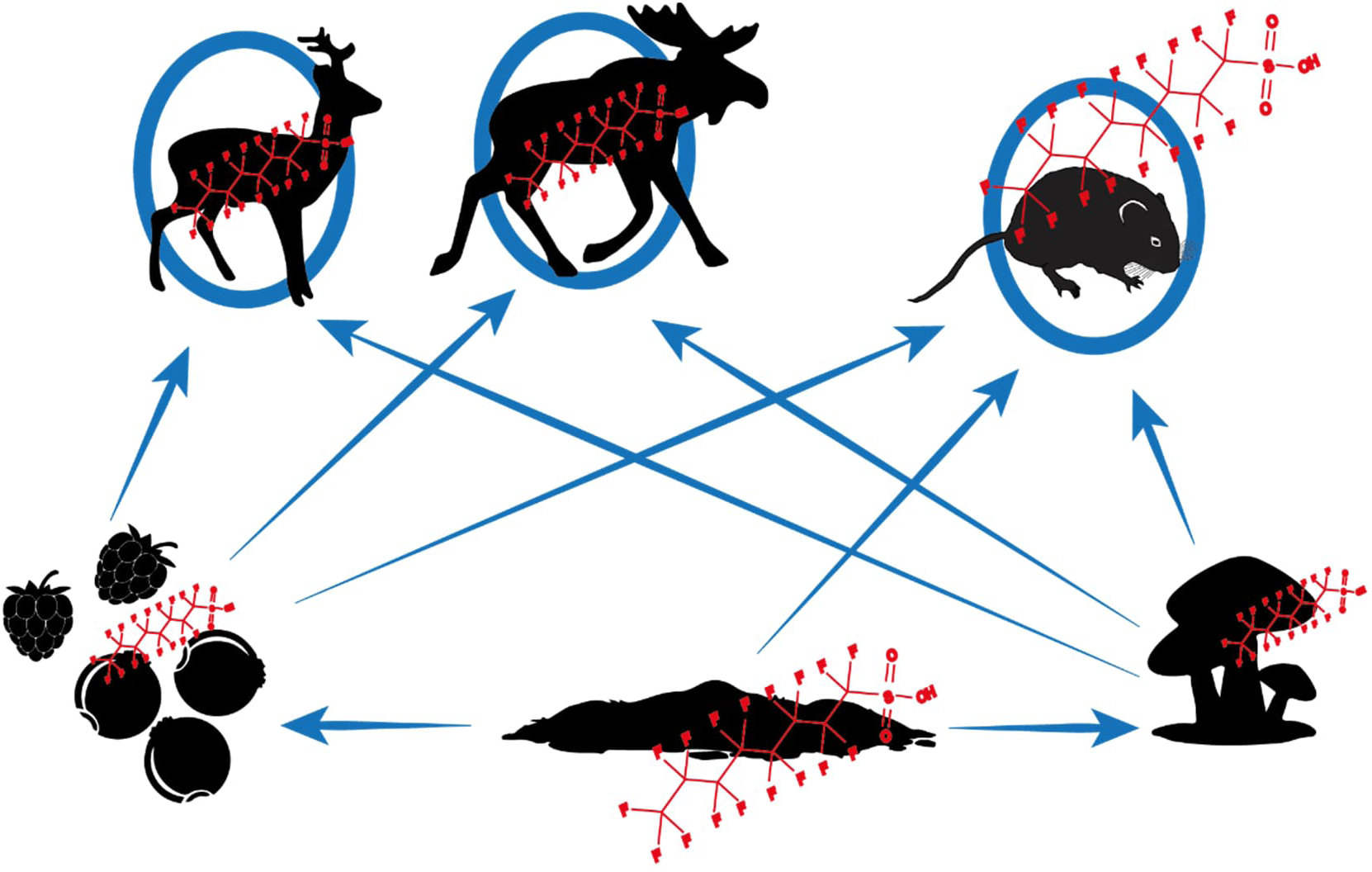
Schematic illustration of the projected bioaccumulation and biomagnification pathways of PFAS (per– and polyfluoroalkyl substances) in the studied trophic network at Frösön, northern Sweden. PFAS in soil are likely to be taken up by dwarf-shrubs including lingonberry (*Vaccinium vitis-ideae*), blueberry (*V. myrtillus*) and raspberry (*Rubus idaeus*) and mushrooms (here sweet tooth *Hydnum repandum*). Berries and mushrooms are important foodstuff for bank voles (*Myodes glareolus*) (Hansson, 1971; Hansson, 1985b) and might also be eaten (either deliberately or accidentally) by ungulates including roedeer (*Capreolus capreolus*) and moose (*Alces alces*) (Cederlund et al., 1980).

## Materials and Methods

### Field sampling

To reflect potential exposure to and bioaccumulation and biomagnification of PFAS, we sampled soil (*n =* 21 pooled samples), berries of dwarf-shrubs lingonberry (*Vaccinium vitis-ideae*) (*n =* 6 pooled samples), blueberry (*V. myrtillus*) (*n =* 6 pooled samples), and raspberry (*Rubus idaeus*) (*n =* 6 pooled samples), the edible mushroom sweet tooth (*Hydnum repandum*) (*n =* 6 pooled samples), as well as tissues and organs of bank voles (*n =* 32 specimens), roedeer (*n =* 10) and moose (*n* = 8). Samples were taken on the island of Frösön (41.6 km^2^) at lake Storsjön, boreal central Sweden, that has been identified as a hotspot of PFAS contamination in Sweden (Axelsson and Bard, 2015; Modin, 2021) (Fig. 2). As a reference, soil, berries and mushrooms were also sampled in reference areas near Umeå, northern Sweden (Fig. 2). Bank voles were snap-trapped opportunistically in forested areas during three trapping sessions (13-14 July, 11-12 August and 13-14 September 2022) using dried apples as bait. Soil, berries, and mushrooms were sampled on Frösön concurrently with the small mammal sampling in August and September 2022, and also in August and September 2022 in the reference areas. Soil samples of ca. 50 g were taken from the organic soil layer (O-horizon) and put in plastic bags. Berries from 40-80 plants/clones were sampled to generate pooled samples of ca. 200 g wet weight (ww). Pooled samples for mushrooms comprised 4-5 fruitbodies to generate 200-250 g ww. Localities had an inter-distance of at least 100 m. All samples were stored in a portable freezer (–20°C) during fieldwork and stored until processing in the laboratory at –20°C. Bank voles were dissected in a BSL-2 (biosafety level 2) laboratory and muscle, liver, heart, kidney, and spleen were extracted and stored at –20°C until further processing. The tissue and organs of the 32 bank voles where prior laboratory analyses pooled to 12 samples in a stratified-random process. First, voles were classified as males and females, and as juveniles and adults, respectively. Individual vole samples were than randomly assigned to one of these functional groups.

**Figure 2.**
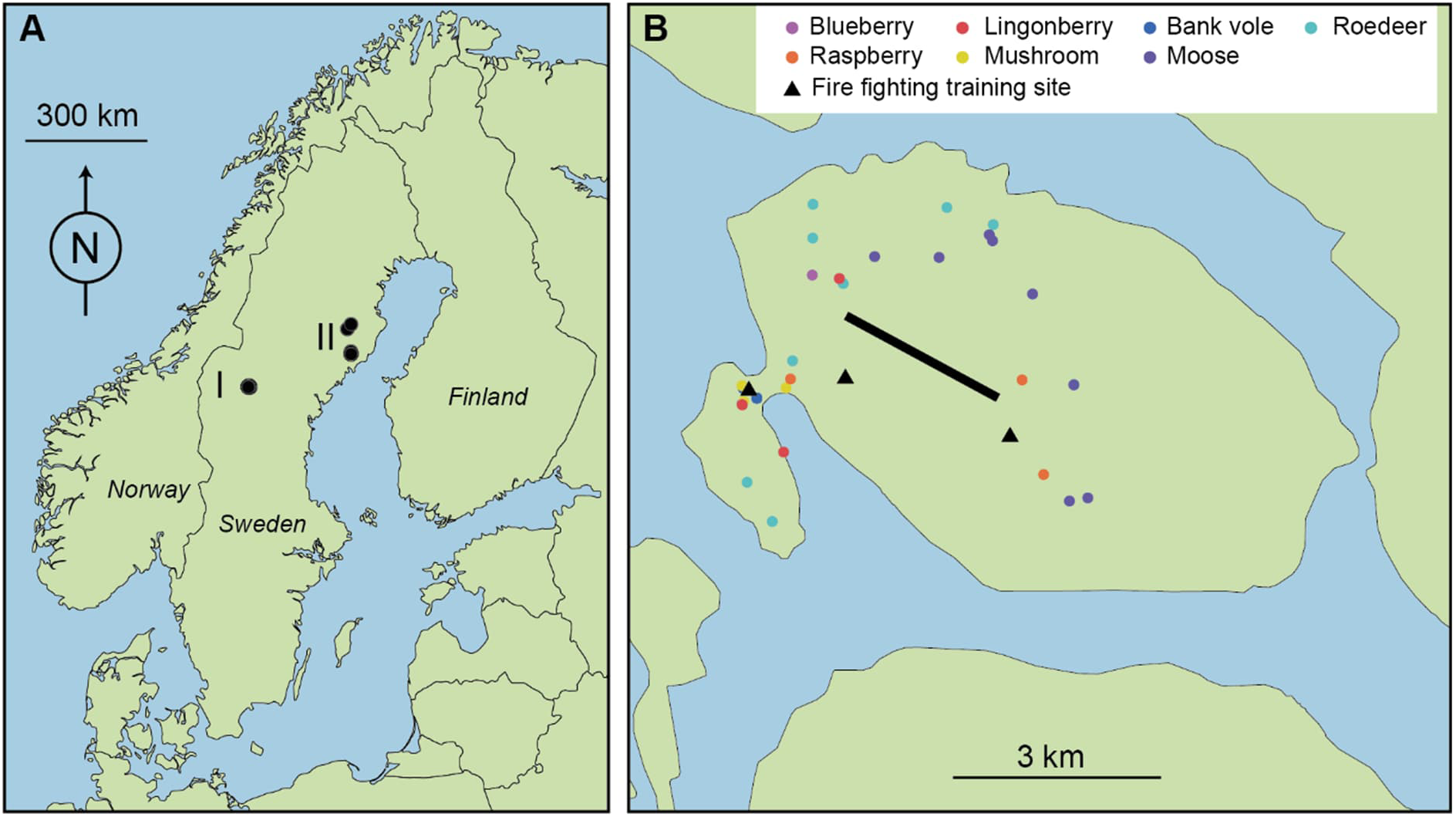
Map of Sweden and neighbouring countries showing (A) the study area on the island Frösön (I) and the reference area near the city of Umeå (II) and (B) a detailed map of the sampling localities (coloured filled circles) for the studied matrices on Frösön. The black line on Frösön in (B) represents the runway of Åre Östersund airport. Soil was sampled at the same sites where berries and mushrooms were sampled, except for some sites where only lingonberries were sampled.

Samples of roedeer and moose on Frösön were provided by local hunting teams during the hunting season 18 September 2021 – 21 January 2022 (see Appendix 1 for details on individual samples). Hunters followed a standard protocol and provided ca. 1 cm^3^ samples of muscle, lung, and liver that they put in plastic zip-bags and stored at –20°C in the premises of the municipality of Östersund. After the hunting season, samples were transported on icepacks to the laboratory.

The sampling of small mammals was approved by the Animal Ethics Committee in Umeå (A 18-2019) and by the Swedish Environmental Protection Agency (NV-07483-19). All applicable institutional and national guidelines for the use of animals were followed.

### Chemicals

In total, 22 target PFAS were included namely C_3_-C_13_ PFCA (PFBA, PFPeA, PFHxA, PFHpA, PFOA, PFNA, PFDA, PFUnDA, PFDoDA, PFTriDA, PFTeDA), C_4_-C_10_ PFSA (PFBS, PFPeS, PFHxS, PFHpS, PFOS, PFNS, PFDS), 4:2, 6:2 and 8:2 fluorotelomer sulfonates (4:2 FTSA, 6:2 FTSA, 8:2 FTSA), and perfluorooctane sulfonamide (FOSA). For PFHxS and PFOS the linear and branched isomers were quantified separately (L-PFHxS, B-PFHxS, L-PFOS, and B-PFOS, respectively). In addition, 17 mass-labeled internal standards (IS) were used, which were spiked to the samples before extraction (Wellington Laboratories): ^13^C_4_– PFBA, ^13^C_5_-PFPeA, ^13^C_5_-PFHxA, ^13^C_4_-PFHpA, ^13^C_8_-PFOA, ^13^C_9_-PFNA, ^13^C_6_-PFDA, ^13^C_7_-PFUnDA, ^13^C_3_– PFDoDA, ^13^C_2_-PFTeDA, ^13^C_3_-PFBS, ^13^C_3_-PFHxS, ^13^C_8_-PFOS, ^13^C_2_-4:2 FTSA, ^13^C_2_-6:2 FTSA, ^13^C_2_-8:2 FTSA, ^13^C_8_-FOSA.

### Sample preparation

All tissue biota samples were extracted using a sample aliquot of approximately 0.5 g homogenized tissue in a Precellys® Evolution vial, add 3 mL acetonitrile and spiked with 5 ng absolute for individual PFAS IS mixture (100 µL of 0.05 µg/mL). The samples were homogenized and extracted using Precellys® Evolution, setting the parameters as 5000 rpm, 2 x 40 sec, 20 sec break in between. Subsequently, sonicate the Precellys® Evolution vial for 30 min and then centrifuge the vial (3000 rpm for min) and transfer the supernatant into a new 15 mL polypropylene (PP) tube. Repeat sonication extraction with 3 mL acetonitrile for 30 min and centrifugation (3000 rpm for 5 min) and transfer the supernatant into the 15 mL PP tube. Put the 15 mL tube into a freezer (–20 °C) for >16 h to crush the proteins. Thereafter, the PP-tube was centrifugated at 4000 rpm at –5 °C for 15 min and the supernatant was transferred into a new PP-tube. Then, the supernatant was concentrated to 1 mL under a gentle steam of nitrogen. For clean-up, the 1 mL extracts were transferred into a 1.7 mL Eppendorf centrifuge tube containing 25 mg ENVI-Carb and 50 μL glacial acetic acid. After vortexing for 30 sec, the Eppendorf centrifuge tube was centrifuged at 4000 rpm at –5°C for 15 min. Finally, the supernatant was transferred into a 1.5 mL PP-vial and stored in a freezer (–20 °C) before analysis.

The soil, mushroom and berry samples were first freeze-dried until the samples were dry (3-7 days). Approximately 1 g dry weight (dw) sample aliquots were weighted into 15 mL PP-tubes. Then, 2.5 ng absolute for individual PFAS IS mixture (50 µL of 0.05 µg/mL) and 3 mL methanol were added and sonicated for 30 min. After centrifugation at 3000 rpm for 15 min, the supernatant was transferred into another 15 mL PP-tube. The extraction was repeated twice with 3 mL methanol with 20 min sonication. The combined extracts were run through an ENVI carb cartridge (1 g, 12 cc) and collect the sample clean tube. The extraction 15 PP-tube was three times with methanol and transferred through the cartridge and then air was pressed through the cartridge using a syringe. Then, the extract was concentrated to 100 µL and transferred to a 1.5 mL PP. Finally, 400 µL methanol was added, vortex and then transferred into a 1.5 mL PP-vial and stored in a freezer (–20 °C) before analysis (for details see Nassazzi et al. (2022)).

### Instrumental analysis

Instrumental analysis was performed using ultra-high pressure liquid-chromatography (SCIEX ExionLC AC system) coupled to tandem mass spectrometry (SCIEX Triple Quad™ 3500) (UHPLC-MS/MS). The column oven was set to 40 °C, and 10 μL of sample were injected into a Phenomenex Kinetex C18 (30 × 2.1 mm, 1.7 μm) precolumn coupled to a Phenomenex Gemini C18 (50 mm × 2 mm, 3 μm) analytical column for chromatographic separation. The mobile phase consisted of MilliQ water with 10 mM ammonium acetate and MeOH. Data evaluation was performed using SciexOS software (2.0) (for details see Nassazzi et al. (2022)).

### Data conversion from fresh weight to dry weight

PFAS in soil, mushrooms and berries were based on freeze-dried samples and hence expressed as concentration in ng/g dry weight. To enable comparison with the wildlife samples that were expressed in ng/g wet weight, we calculated factors for conversion from wet weight to dry weight. For the ungulate samples, we used 0.5-1.4 g wet weight from the samples of the 18 specimens that were included in our study, freeze-dried them (typically 5 days and up to 7 days until samples were dry) and calculated the water content.

For the vole samples, we unfortunately did not have any material left after the PFAS analyses. For the conversion of concentrations in muscle, liver, kidney and spleen, we therefore used data on tissue/organ wet and dry weight from 64 bank vole specimens that were included in Ecke et al. (2018). To achieve conversion factors between heart and dry weight in bank vole hearts, we used hearts from 19 bank voles trapped in spring and autumn 2022 near the city of Umeå, northern Sweden that we freeze-dried like the ungulate samples.

We calculated the dry weight of the samples according to

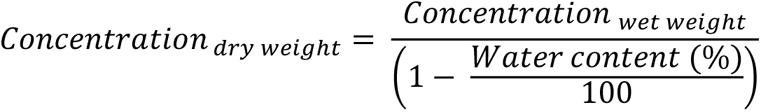

### Statistical analyses

For statistical analyses, concentrations that were below limit of quantification (LOQ) were replaced by half of LOQ. We only included PFAS in the statistical analyses that were detected in at least two samples. Based on the results of the analysed PFAS, we combined the respective PFAS to calculate the sum of all PFAS, PFAS4 (sum of PFOA, PFNA, PFHxS, and PFOS), PFCAs, PFSAs, and the sum of precursors.

To assess potential bioaccumulation of PFAS from soil to biota and from mushrooms and berries to herbivores (bank voles and ungulates), we calculated bioaccumulation factors according to

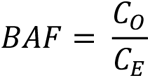

with *BAF* representing the bioaccumulation factor, *C_O_* the concentration in the organism and *C_E_* the concentration in the environment. We calculated BAFs from soil to biota and from mushrooms and berries to herbivores. BAFs >1 were interpreted as bioaccumulation with risk for biomagnification.

To meet assumptions for parametric tests, concentrations were Box-Cox-transformed prior analyses. We used principal component analysis (Jongman et al., 1995) to reduce the PFAS that had no missing values to a few essential components. Only principal components (PCs) that explained more than 10% of the variance among the PFAS variables were considered.

We computed one-way ANOVA on PFAS concentrations on Frösön to analyse if concentrations (all expressed in dry weight) varied among matrices (soil, raspberry, blueberry, lingonberry, sweet tooth, bank vole muscle, kidney, heart, liver, and spleen, as well as roedeer and moose kidney, lung, and liver). Differences in PFAS concentrations between Frösön and the reference localities were analysed by *t*-tests.

Statistical analyses were performed in R (Team, 2021) using the packages ‘car’, ‘stats’ and ‘vegan’ supported by visualisation with packages ‘ggplot2’ and ‘ggfortify’.

When relating identified PFAS concentrations to thresholds for food safety, we replaced all values below LOQ by 0.

## Results

Of the 22 analysed PFAS, all PFAS were detected in at least two samples and considered for visualization and statistical analysis, and 15 PFAS had no missing values (see Appendix 2).

Reducing the dimensions of PFAS, the first two PCs explained 66% of all variation among PFAS concentrations with the first PC that explained 46% of the variation being dominated by in ascending importance PFHpS, 8:2 FTSA, L-PFHxS, PFDS, PFDoDA, L-PFOS, PFDA, and PFNA, while PC2 that explained 20% was dominated by PFHpA, PFOA, PFBS, and PFHxA (Fig. 3a). Bank vole samples where mainly associated with PC1 and their partly high PC scores indicated high PFAS concentrations (Fig. 3b, c). In addition, one soil and one mushroom sample from Frösön indicated high PFAS concentrations (Fig. 3b). Non-vole samples separated mainly along the PFAS gradient of PC2 (Fig. 3c).

**Figure 3.**
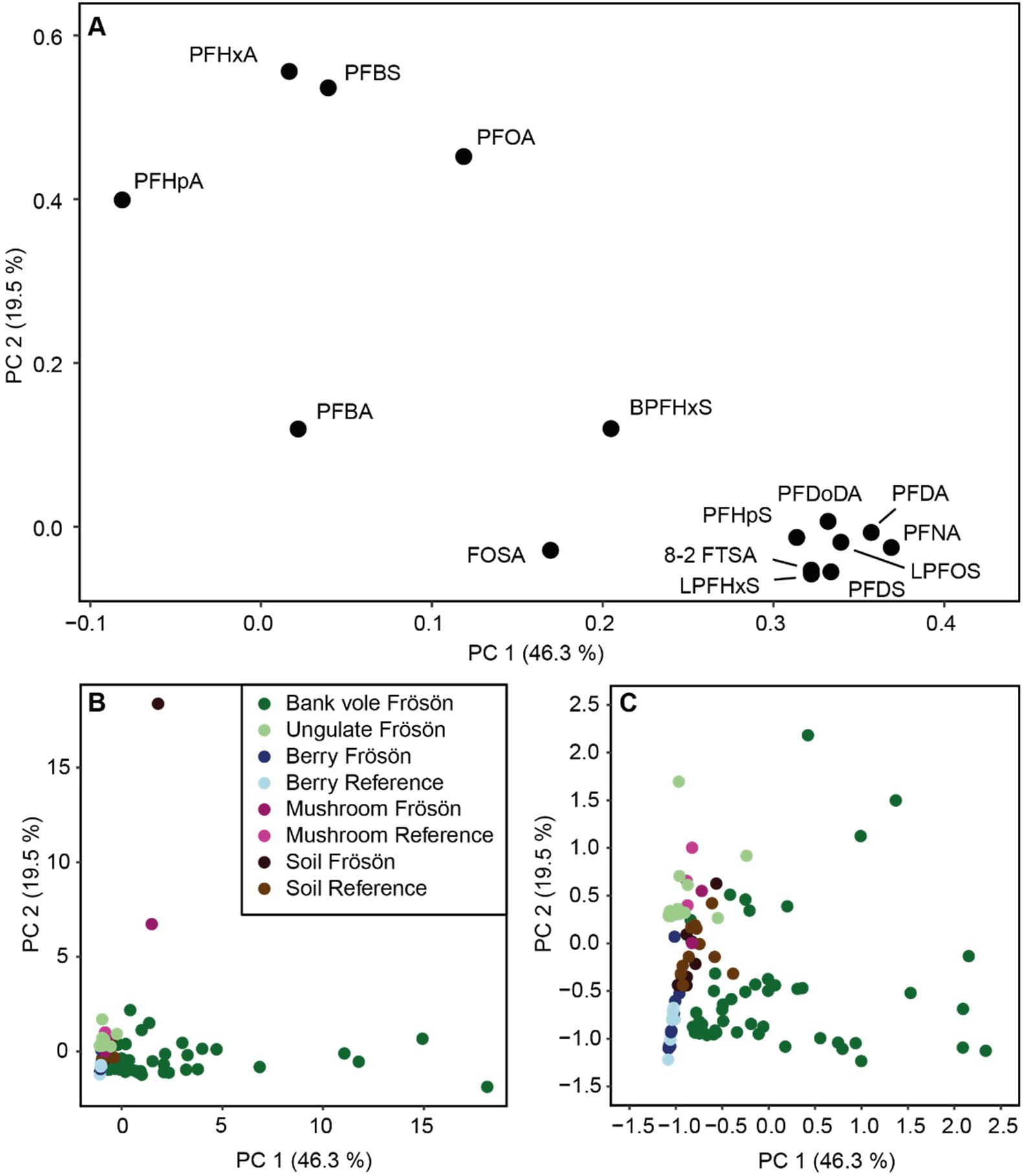
Principal component analysis (PCA) on 14 PFAS in environmental samples (soil, mushrooms, berries, ungulates, and bank voles on the island of Frösön and in the reference areas. The PC loadings (A) illustrate the compositional similarities among the different PFAS, while the PC scores (B, C) illustrate the similarities in PFAS concentrations among the different samples, with (C) representing an enlargement of the PC scores for PC1 and PC2, respectively, < 2.5. Percentage of explained variation are given for the first two PCs. PFHXs is represented by its linear and branched isomers.

The general patterns of the PCA were confirmed by further statistical analyses. PFAS concentrations on Frösön showed large variations among the studied species, tissues, and organs. The mean concentrations ± standard error (ng/g dw) of the sum of PFAS showed the following ascending pattern for soil, mushroom and berries: lingonberry (1.8 ± 1.2, *n =* 3) < raspberry (3.1 ± 1.1, *n =* 3) < blueberry (7.3 ± 1.9, *n =* 3) < sweet tooth (54.7 ± 20.6; *n =* 3) << soil (196 ± 179, *n =* 10) (see also Fig. 4, Appendix 2). For the wildlife samples that were expressed in wet weight, the mean concentrations ± standard error (ng/g ww) showed the following pattern: lung (moose) (3.2 ± 0.5, *n =* 7) ≈ muscle (moose) (3,7 ± 0,7; *n =* 7) < roedeer (muscle) (5.0 ± 0.5, *n =* 9) < lung (roedeer) (7.9 ± 3.6, *n =* 10) < liver (moose) (55.4 ± 16.3, *n =* 8) < liver (roedeer) (84.0 ± 46,7; *n =* 10) << muscle (bank vole) (238 ± 78.0, *n =* 12) << heart (bank vole) (458 ± 130, *n =* 12) << spleen (bank vole) (621 ± 210, *n =* 12) <<< kidney (bank vole) (2280 ± 829, *n =* 12) <<< liver (bank vole) (4790 ± 1230, *n =* 12) (see also Fig. 5-6, Appendix 2).

**Figure 4.**
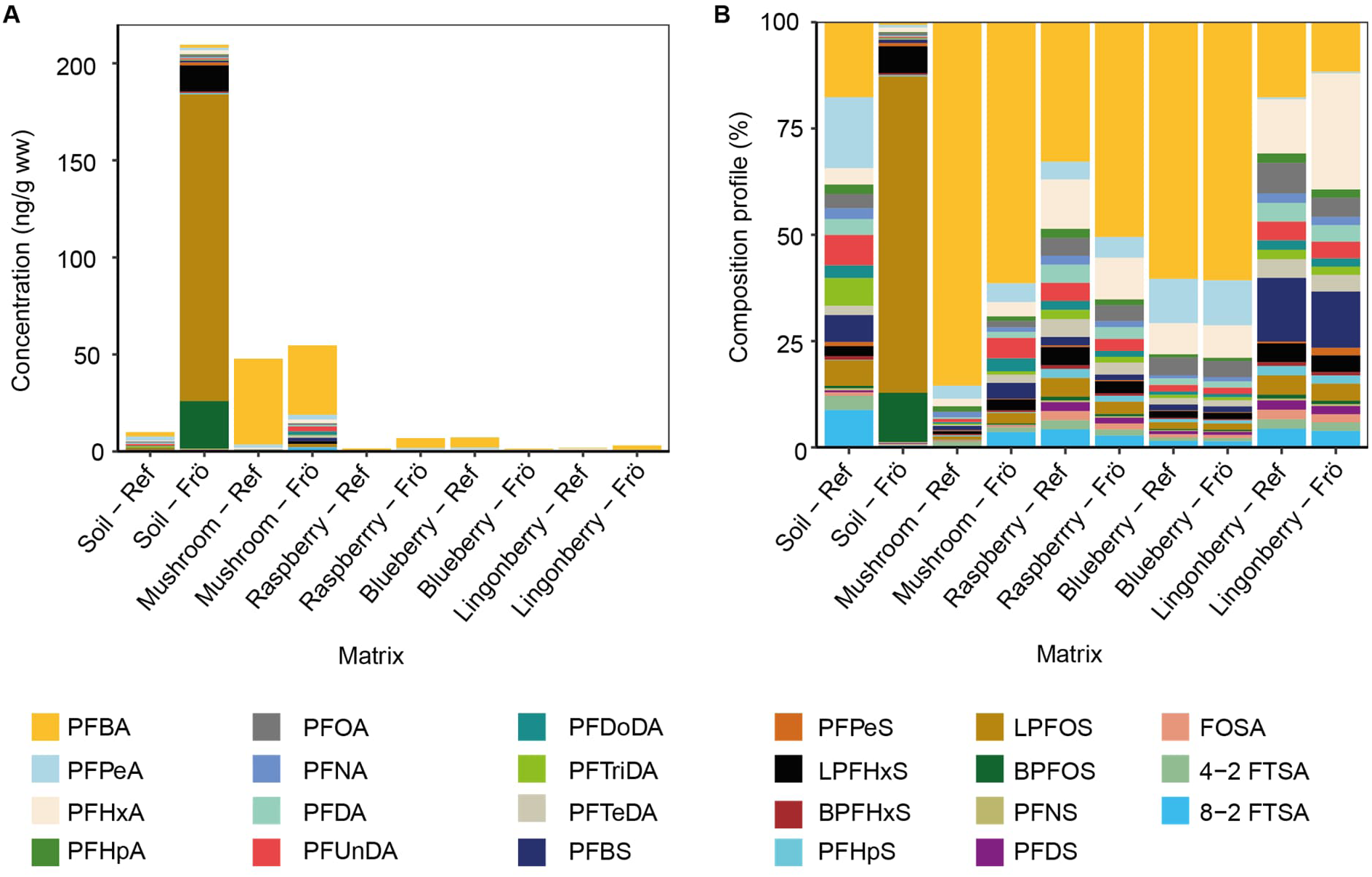
Mean concentrations of PFAS (per– and polyfluoroalkyl substances) in soil, mushrooms, berries, in the reference area (Ref; see Figure 2) and on the island of Frösön (Frö), northern Sweden (A). (B) represents the composition profile (relative concentration) of the studied PFAS.

**Figure 5.**
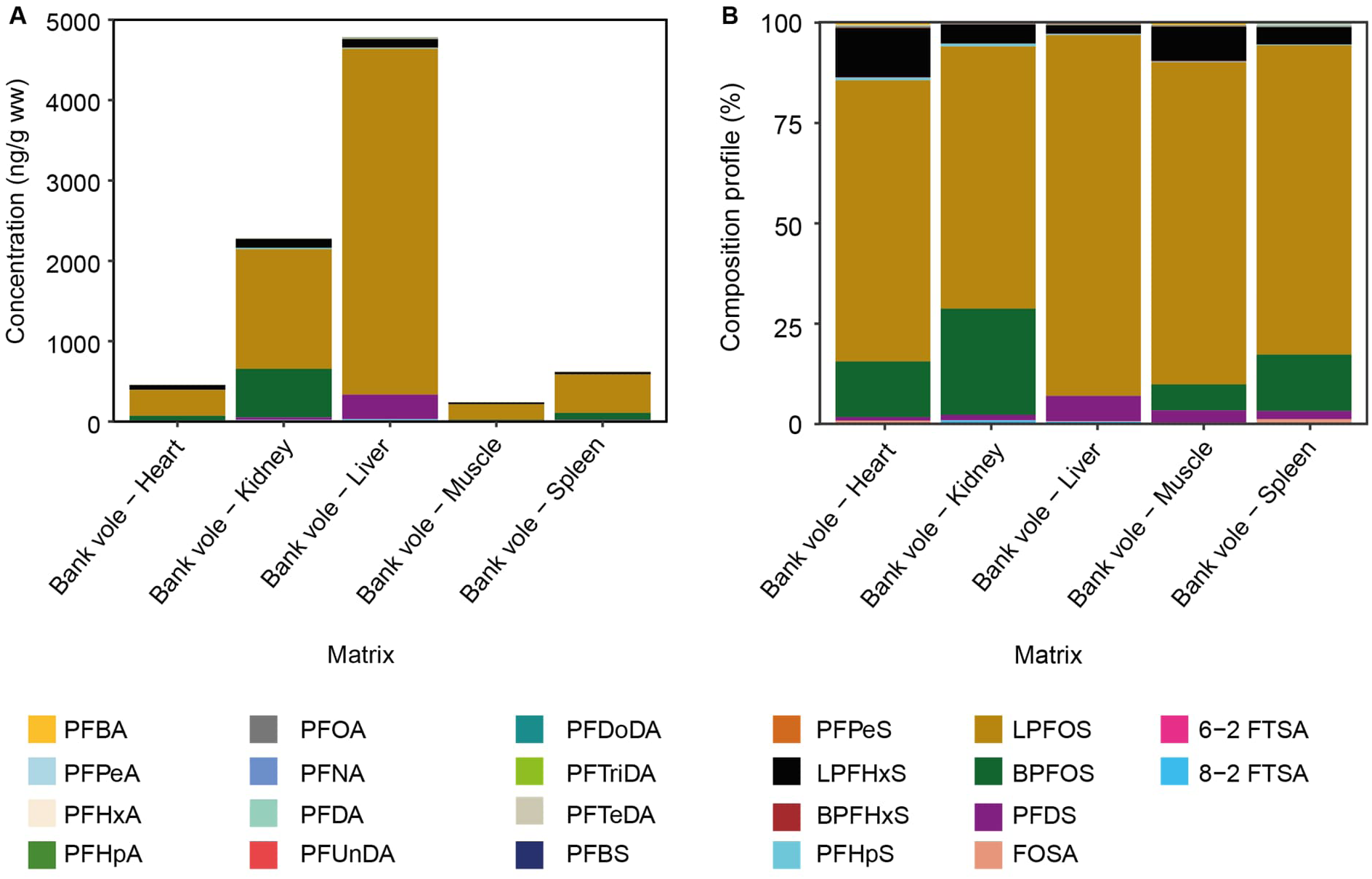
Mean concentrations of PFAS (per– and polyfluoroalkyl substances) in organs and tissue of bank voles on the island of Frösön, northern Sweden (A). (B) represents the composition profile (relative concentration) of the studied PFAS.

**Figure 6.**
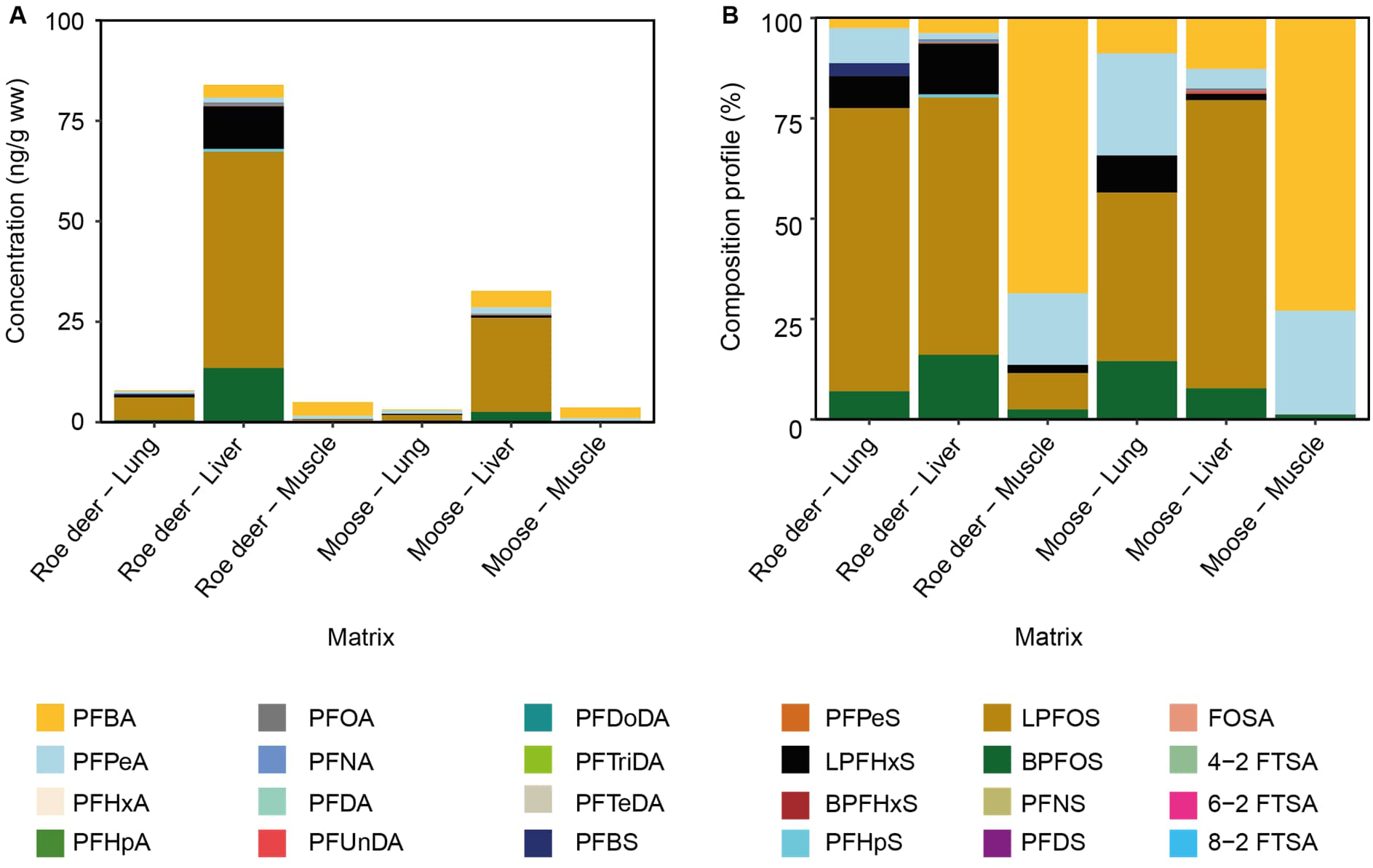
Mean concentrations of PFAS (per– and polyfluoroalkyl substances) the ungulates roedeer and moose on the island of Frösön, northern Sweden (A). (B) represents the composition profile (relative concentration) of the studied PFAS.

The concentrations of the 22 PFAS differed among studied samples (Table 1, see also Fig. 4-6). The linear and branched PFOS in soil showed higher concentrations on Frösön compared with the reference localities, while the concentrations of all other PFAS did not show any difference (Appendix 3; see also Fig. 4). PFAS in neither berries nor the mushroom differed between Frösön and the reference localities (*t*-tests; Appendix 3).

**Table 1.**
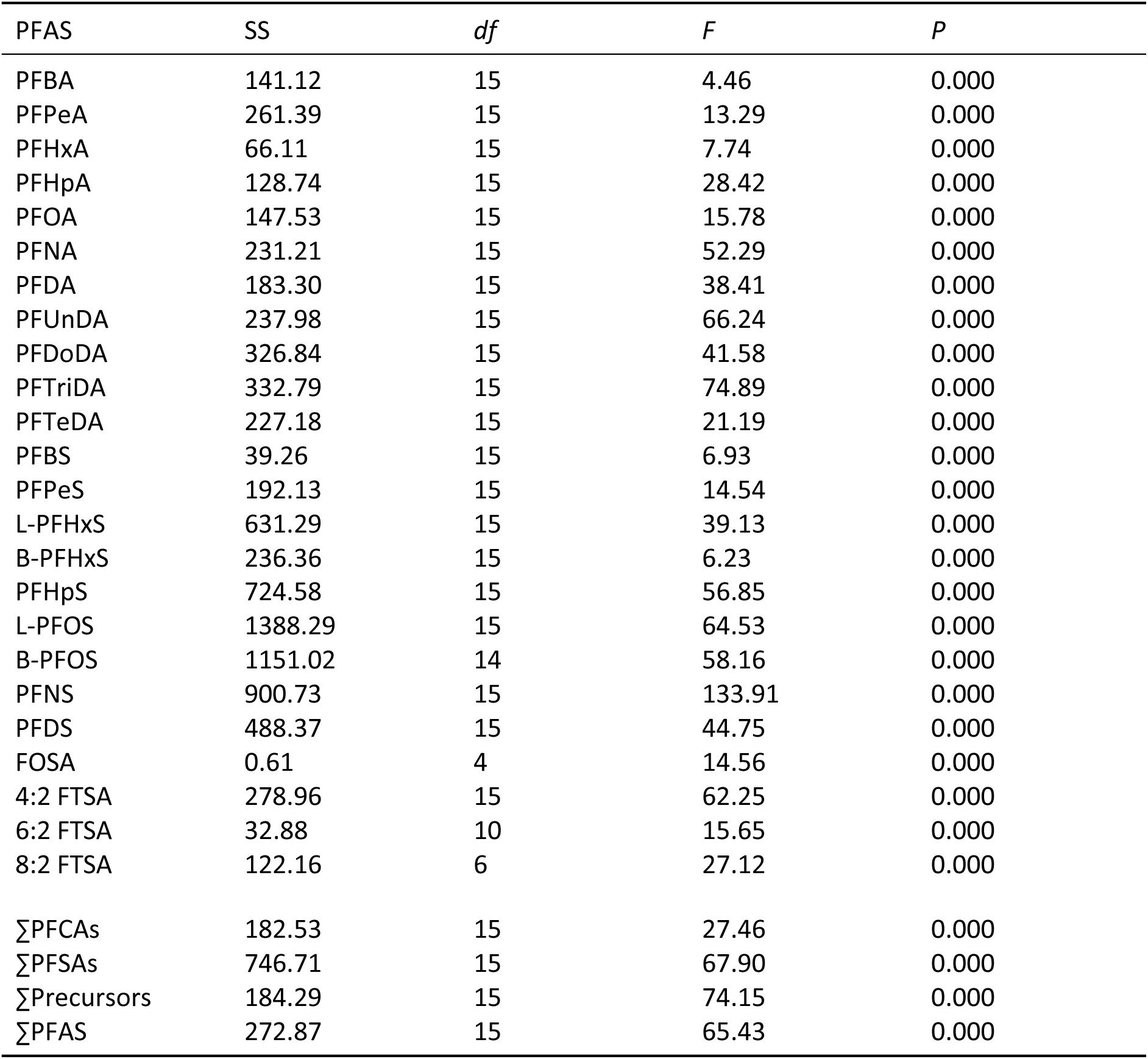
Analysis of Variance (ANOVA) of different PFAS and their sums on Frösön showing sum of squares (SS), degrees of freedom (*<u>df</u>*), *F*-statistic (*F*) and *p*-values (*P*). Differences were tested among soil, fruits of lingonberry, blueberry, and raspberry, mushrooms, and tissues in bank voles (muscle, heart, lung, liver, kidney, and spleen), roedeer and moose (muscle, liver, kidney).

The PFAS profiles varied among samples and localities. In soil on Frösön, L-PFOS was on average the dominating PFAS (∼74%), while PFAS in soil samples in the reference localities were more evenly distributed with PFBA, PFPeA, and 8:2 FTSA contributing with a total of ∼43% (Fig. 4). On Frösön, the PFAS profiles in mushroom, raspberry and blueberry were dominated by PFBA, while PFHxA followed by PFBS and PFBA dominated in lingonberry (Fig. 4). This pattern was similar in the reference samples, except that PFBA only constituted to ca. 1/3 of PFAS concentrations in raspberry, while PFBA were the dominating PFAS on Frösön (Fig. 4).

In bank voles, PFAS concentrations were highest in liver with L-PFOS being the dominating PFAS (>65%) in all tissues and organs (Fig. 5). Also in roedeer and moose, PFAS concentrations were highest in liver and like in bank voles dominated by L-PFOS in liver and lung (Fig. 6). However, muscle in the ungulates compared with bank vole were dominated by PFBA (Fig. 5, 6).

Summarizing PFAS into PFCAs, PFSAs, and precursors revealed that the reference soil samples, as well as all mushroom and berry samples and ungulate muscles were dominated by PFCAs (Fig. 7). In contrast, all vole samples, ungulate lung samples and roe deer liver were dominated by PFSAs and the proportion of PFCAs and PFSAs was similar in moose liver (52% and 48%, respectively). (Fig. 7).

**Figure 7.**
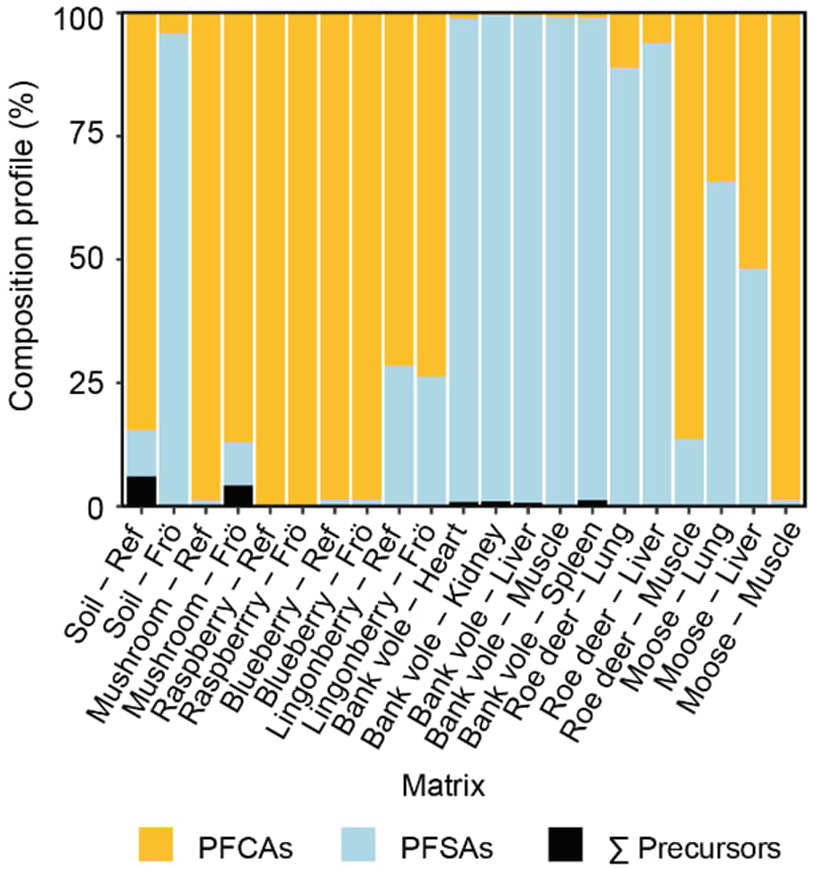
Composition profiles (relative concentration) of PFAS (per– and polyfluoroalkyl substances) divided into PFCAs (perfluoroalkyl carboxylic acids), PFSAs (perfluoroalkyl sulfonic acids) and the sum (Σ) of precursors in soil, mushrooms, berries, and organs and tissue of bank voles as well as the ungulates roedeer and moose in the reference area (Ref; see Figure 2) and on the island of Frösön (Frö), northern Sweden. Bank voles and ungulates were only sampled on Frösön.

The three roedeer with the highest PFAS4 concentrations (>45 ng/g ww) were hunted on the isthmus Bynäset or close to the isthmus, i.e., close to the firefighting training site on Bynäset (westernmost firefighting training site in Fig. 2). It is also on this isthmus all bank voles were trapped and none of these had a PFAS4 concentration lower than 474 ng/g ww (max 11 600 ng/g ww).

The bioaccumulation factors (BAFs) for several PFAS in bank voles were overall higher for mushrooms and soil than for berries and were in 27 cases >100 (Fig. 8, Appendix 4). In contrast, the bioaccumulation factors for PFAS in the ungulates were overall low for soil but were >1 for several PFAS originating from especially mushrooms and were even >100 for L-PFOS and B-PFOS in a liver of a roedeer (Fig. 8, Appendix 4). Overall, in ungulates, bioaccumulation factors were lower in lung and muscle than in liver, while in voles all studied tissues and organs showed bioaccumulation for several PFAS that originated from multiple sources (Fig. 8, Appendix 4).

**Figure 8.**
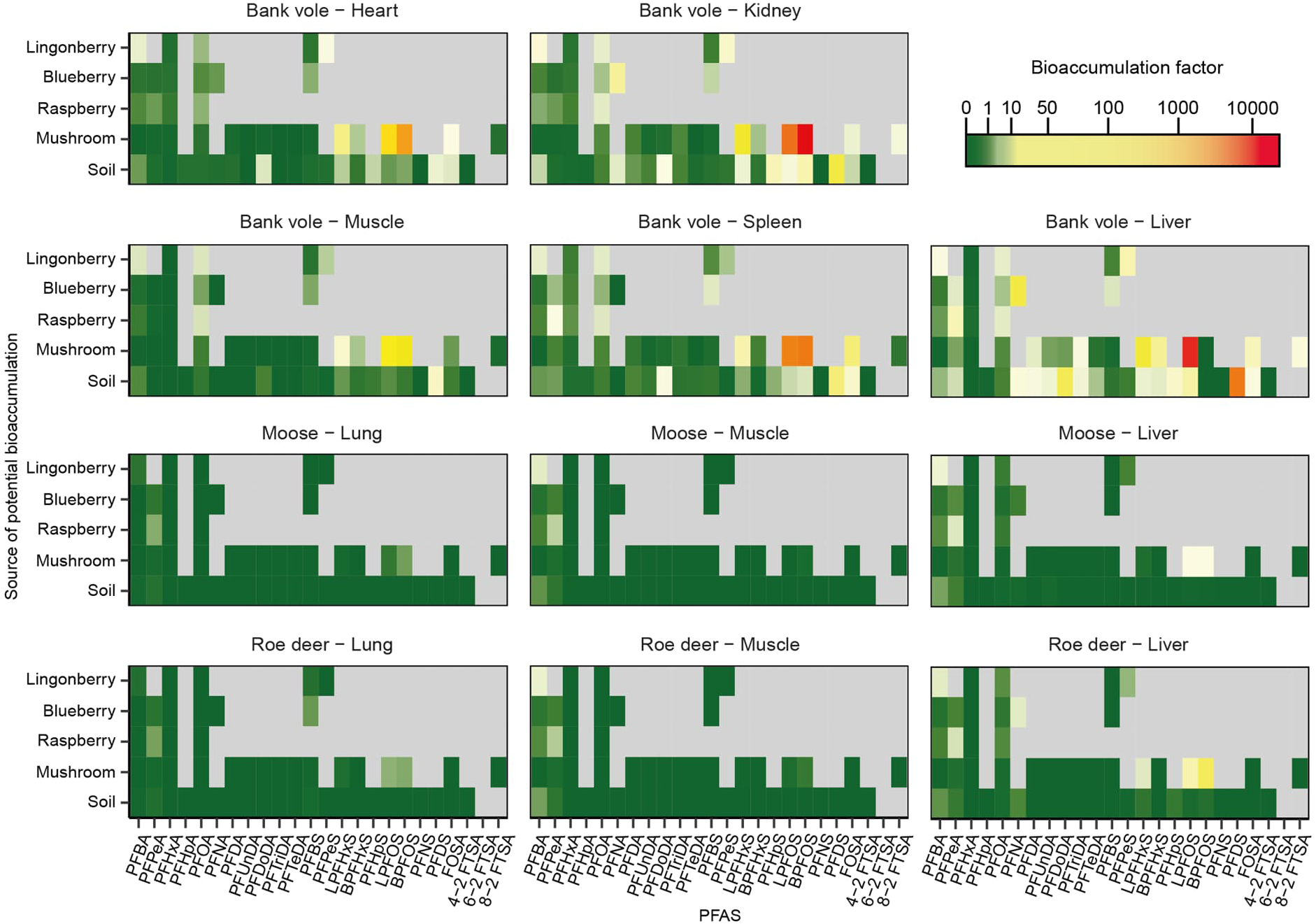
Bioaccumulation factors represented as colour ramp for 24 PFAS in tissue and organs of bank voles, roedeer and moose. The y-axis represents the potential source of bioaccumulation. Grey areas represent missing data.

## Discussion

Our study revealed partly extremely high concentrations of PFOS in biota and provides valuable insights in the potential trophic fate of PFAS from soil via mushrooms and berries to herbivores. As expected, PFAS concentrations were highest in liver of bank vole, roedeer and moose, which is in line with previous findings in rodents (Costello et al., 2022) and ungulates (Falk et al., 2012).

Despite rather high concentrations of PFOS in soil, this PFAS was not accumulated by berries, while the mushroom sweet tooth showed high affinity for PFOS, which is also supported by previous studies on oyster mushroom (*Pleurotus ostreatus*) (Golovko et al., 2022).

The high bioaccumulation factors in bank voles for several PFAS originating in soil and/or mushroom suggest bioaccumulation, which is also supported by similar PFAS profiles of soil on Frösön and bank vole tissue and organs. These findings are further supported by the feeding ecology of bank voles. Bank voles frequently feed on berries of raspberry, blueberry, and lingonberry (Hansson, 1979a; Hansson, 1985a; Hansson, 1985b). This is likely also the case in our study, and the rather low PFAS concentrations in berries in the present study did not result in bioaccumulation in bank voles. Also mushrooms are important foodstuff for bank voles (Hansson, 1979a), which in our study might explain mushroom-dependent bioaccumulation of PFAS in bank voles. However, the PFAS profiles in mushrooms differ from those in bank voles, which suggests that also other foodstuff might be involved in the bioaccumulation of PFAS in the food web on Frösön. Indeed, bioaccumulation pathways from sweet tooth to bank voles need to be assessed more carefully in future studies.

The identified PFAS concentrations in soil, sweet tooth, and wildlife – especially in bank vole – is worrisome from the perspective of ecosystem health and One Health. Our study suggests soil to be an important PFAS source for bank voles. We did not study the gut content of the bank voles. We therefore do not know their actual foodstuff and whether the here studied potential PFAS sources (soil, mushrooms, berries) are the actual sources of the high PFAS concentrations in bank voles. The calculated bioaccumulation factors however strongly suggest soil and mushrooms to be important PFAS sources for bank voles. Despite bank voles being classified as herbivores, they also consume invertebrates and mineral particles are commonly found in their stomach (Hansson, 1971; Hansson, 1979b; Hansson, 1985a; Hansson, 1985b). In fact, future studies should include the role of macroinvertebrates in general and earthworms in particular for bioaccumulation of PFAS in bank voles (sensu Grønnestad et al., 2019). Isotopic instead of foodstuff analyses of gut content can then be used to identify food sources (e.g., Magnusson et al., 2019). Bank voles dig their own burrows and dust formed during digging – resulting in either inhaling or digestion of PFAS-coated dust particles – could be an important PFAS source.

The bank vole is Europe’s most common mammal (Mitchell-Jones et al., 1999) and important staple food for many mammalian and avian predators (Hipkiss and Hörnfeldt, 2004; Hipkiss et al., 2008; Hörnfeldt et al., 1990). Tengmalm’s owl (*Aegulius funereus*) depends to up to 90% on bank voles as prey species (unpublished data). The partly identified extreme PFAS concentrations found in bank voles (>4000 ng/g ww) compared with for example 16 ng/g ww in a PFAS contaminated skiing area in Norway (Grønnestad et al., 2019) are therefore not only a potential health risk for the bank voles, but also for their predators. Since the predators have broader home ranges than their prey (Tengmalm’s owl for example ca. 2 km^2^; Kouba et al., 2017), predators could in addition contribute to secondary distribution of PFAS.

PFAS compromises the immune system of mammals (Beans, 2021). The high PFAS concentrations in bank voles are therefore a potential health risk for the specimens and could induce increased susceptibility to infection with Puumala orthohantavirus (PUUV). PUUV causes the zoonotic disease nephropathia epidemica (vole fever) in humans (Khalil et al., 2019; Khalil et al., 2014; Vapalahti et al., 2003). PFAS in bank voles could therefore potentially also pose a threat to human health.

Mountain hare (*Lepus timidus*) is a common and popular game species in northern Sweden. The species was not included in our study. Ecologically, mountain hare resembles bank voles. The high PFAS concentrations found in bank voles therefore urge to gain knowledge on PFAS in mountain hare, which is also important from a food safety perspective.

Currently, there are no maximum levels for human consumption of PFAS in mushrooms and berries. For the sum of PFAS, the concentrations in sweet tooth on Frösön were all higher than 21 ng/g dw (max 92.6 ng/g dw; PFAS4 max 11.3 ng/g dw) suggesting that consumption of this species should be limited. The non-difference of PFAS concentrations in mushroom and berries between Frösön and the reference areas was surprising. This might either suggest that mushrooms and the studied plant species do not accumulate PFAS or that also the reference areas were impacted by PFAS originating from diffuse sources.

The maximum levels for PFOS, PFOA, PFNA, PFHxS, and their sum (PFAS4) in meat (5.0, 3.5, 1.5, 0.6, and 9.0, respectively) (European Commission, 2022) were not exceeded in any of the samples. However, the muscle of one roedeer was close to the threshold level of 5.0 ng/g ww for PFOS (4.3 ng/g ww). The maximum level for PFOS in offal (50 ng/g ww) were however exceeded in liver of two roedeer.

The European Food Safety Authority (EFSA) recommends a tolerable weekly intake (TWI) for PFAS4 of 4.4 ng/kg body weight per week^1^. This threshold will be exceeded if regularly consuming roedeer meat from the island Frösön. Upon consumption of for example 200 g roedeer meat, a person with a body weight of 70 kg will have an intake of 860 ng PFOS corresponding to 12.3 ng/kg. Based on our results, authorities should therefore consider making recommendations for consumption of game meat that originates from or has parts of its home range on Frösön.

## Acknowledgements

We thank the hunting teams on Frösön for supporting our study with samples from roedeer and moose, Åke Nordström for field and lab assistance, and Mikael Marberg for lab assistance. Hanna Modin provided important insight into PFAS on Frösön, administrated the distribution of sampling kits to hunters and handled storage of samples from game species. The study was funded by Future One Health, Swedish University of Agricultural Sciences (SLU.ua.2021.4.1-3058) and the Swedish Environmental Protection Agency (NV-06350-22).

---

**Appendix 1.**
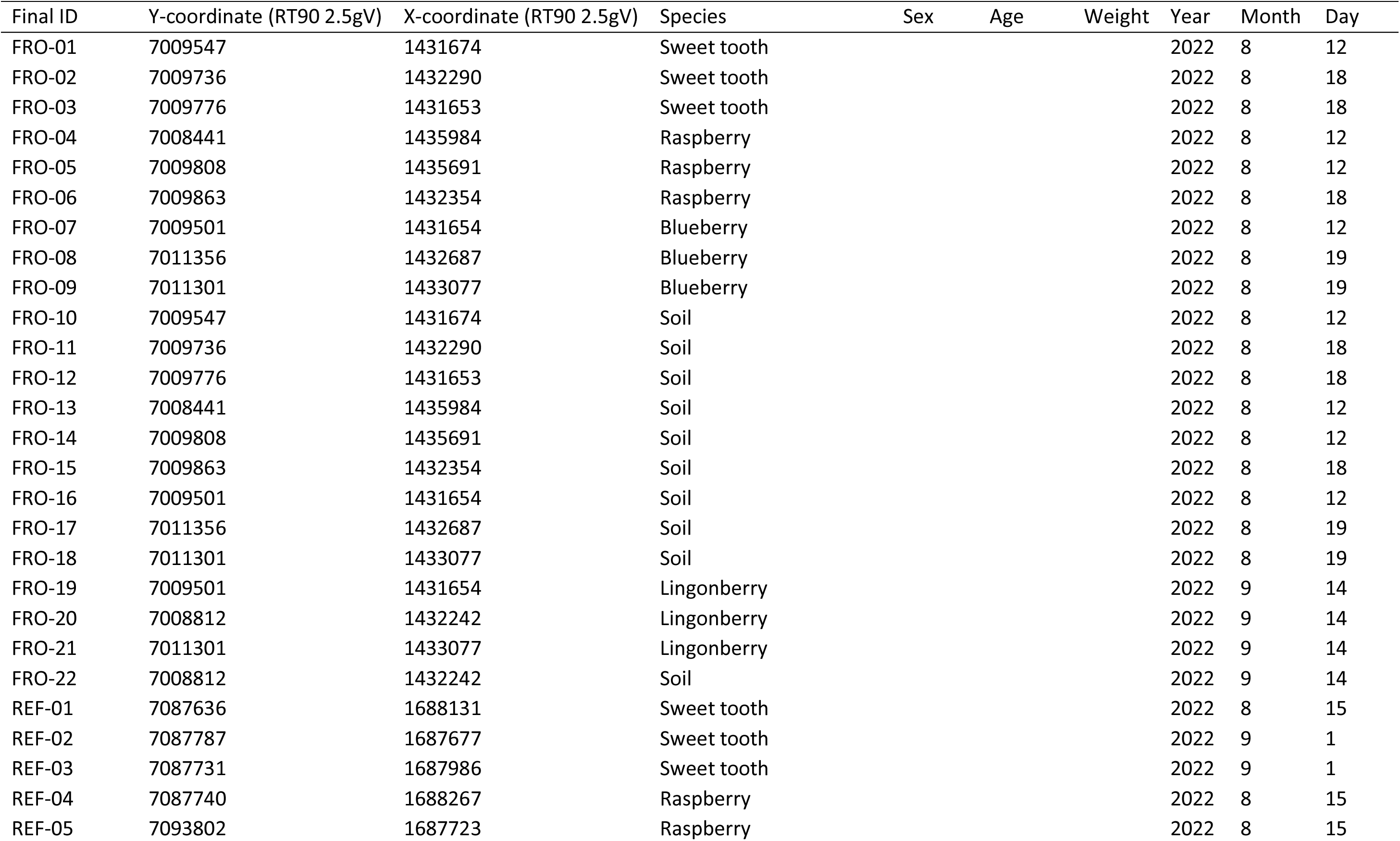
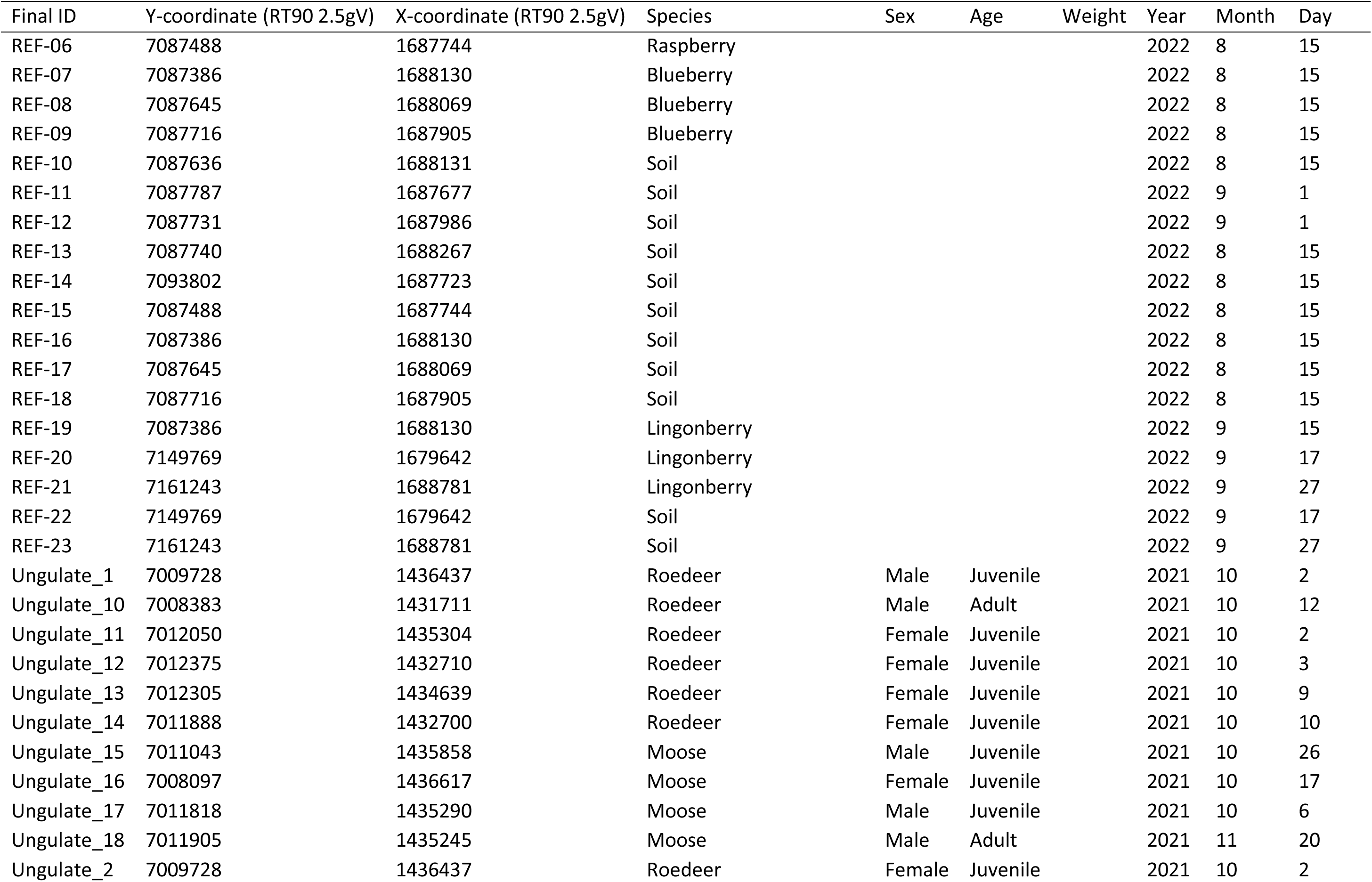
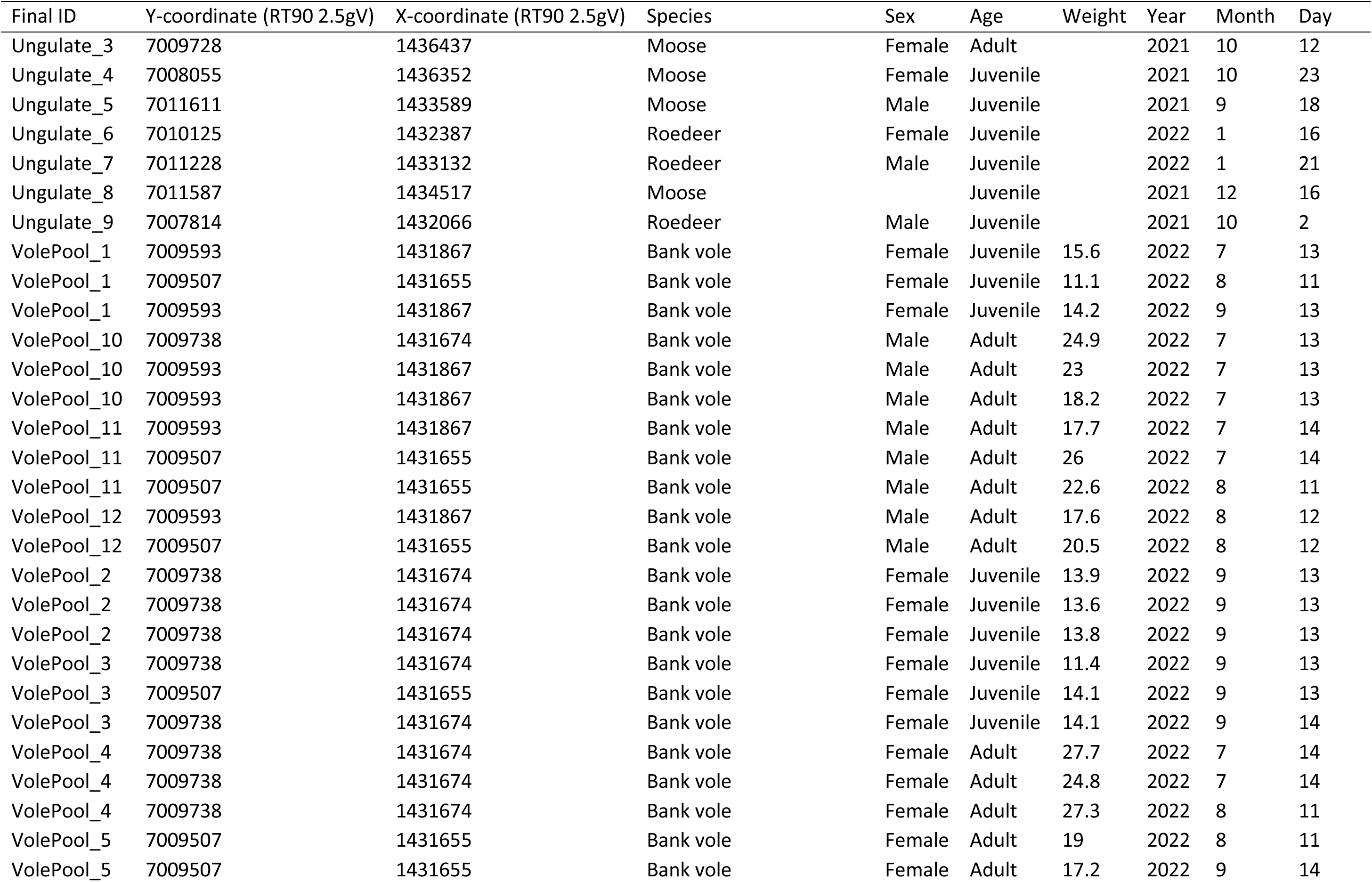
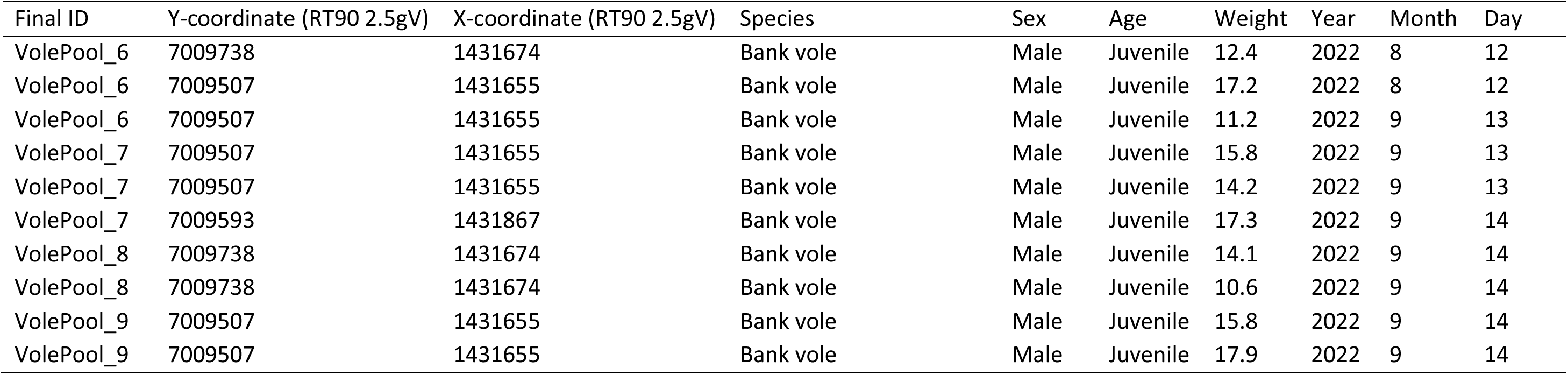
Details for each sample included in the study.

| Final ID | Y-coordinate (RT90 2.5gV) | X-coordinate (RT90 2.5gV) | Species | Sex | Age | Weight | Year | Month | Day |
| --- | --- | --- | --- | --- | --- | --- | --- | --- | --- |
| FRO-01 | 7009547 | 1431674 | Sweet tooth |  |  |  | 2022 | 8 | 12 |
| FRO-02 | 7009736 | 1432290 | Sweet tooth |  |  |  | 2022 | 8 | 18 |
| FRO-03 | 7009776 | 1431653 | Sweet tooth |  |  |  | 2022 | 8 | 18 |
| FRO-04 | 7008441 | 1435984 | Raspberry |  |  |  | 2022 | 8 | 12 |
| FRO-05 | 7009808 | 1435691 | Raspberry |  |  |  | 2022 | 8 | 12 |
| FRO-06 | 7009863 | 1432354 | Raspberry |  |  |  | 2022 | 8 | 18 |
| FRO-07 | 7009501 | 1431654 | Blueberry |  |  |  | 2022 | 8 | 12 |
| FRO-08 | 7011356 | 1432687 | Blueberry |  |  |  | 2022 | 8 | 19 |
| FRO-09 | 7011301 | 1433077 | Blueberry |  |  |  | 2022 | 8 | 19 |
| FRO-10 | 7009547 | 1431674 | Soil |  |  |  | 2022 | 8 | 12 |
| FRO-11 | 7009736 | 1432290 | Soil |  |  |  | 2022 | 8 | 18 |
| FRO-12 | 7009776 | 1431653 | Soil |  |  |  | 2022 | 8 | 18 |
| FRO-13 | 7008441 | 1435984 | Soil |  |  |  | 2022 | 8 | 12 |
| FRO-14 | 7009808 | 1435691 | Soil |  |  |  | 2022 | 8 | 12 |
| FRO-15 | 7009863 | 1432354 | Soil |  |  |  | 2022 | 8 | 18 |
| FRO-16 | 7009501 | 1431654 | Soil |  |  |  | 2022 | 8 | 12 |
| FRO-17 | 7011356 | 1432687 | Soil |  |  |  | 2022 | 8 | 19 |
| FRO-18 | 7011301 | 1433077 | Soil |  |  |  | 2022 | 8 | 19 |
| FRO-19 | 7009501 | 1431654 | Lingonberry |  |  |  | 2022 | 9 | 14 |
| FRO-20 | 7008812 | 1432242 | Lingonberry |  |  |  | 2022 | 9 | 14 |
| FRO-21 | 7011301 | 1433077 | Lingonberry |  |  |  | 2022 | 9 | 14 |
| FRO-22 | 7008812 | 1432242 | Soil |  |  |  | 2022 | 9 | 14 |
| REF-01 | 7087636 | 1688131 | Sweet tooth |  |  |  | 2022 | 8 | 15 |
| REF-02 | 7087787 | 1687677 | Sweet tooth |  |  |  | 2022 | 9 | 1 |
| REF-03 | 7087731 | 1687986 | Sweet tooth |  |  |  | 2022 | 9 | 1 |
| REF-04 | 7087740 | 1688267 | Raspberry |  |  |  | 2022 | 8 | 15 |
| REF-05 | 7093802 | 1687723 | Raspberry |  |  |  | 2022 | 8 | 15 |
| REF-06 | 7087488 | 1687744 | Raspberry |  |  |  | 2022 | 8 | 15 |
| REF-07 | 7087386 | 1688130 | Blueberry |  |  |  | 2022 | 8 | 15 |
| REF-08 | 7087645 | 1688069 | Blueberry |  |  |  | 2022 | 8 | 15 |
| REF-09 | 7087716 | 1687905 | Blueberry |  |  |  | 2022 | 8 | 15 |
| REF-10 | 7087636 | 1688131 | Soil |  |  |  | 2022 | 8 | 15 |
| REF-11 | 7087787 | 1687677 | Soil |  |  |  | 2022 | 9 | 1 |
| REF-12 | 7087731 | 1687986 | Soil |  |  |  | 2022 | 9 | 1 |
| REF-13 | 7087740 | 1688267 | Soil |  |  |  | 2022 | 8 | 15 |
| REF-14 | 7093802 | 1687723 | Soil |  |  |  | 2022 | 8 | 15 |
| REF-15 | 7087488 | 1687744 | Soil |  |  |  | 2022 | 8 | 15 |
| REF-16 | 7087386 | 1688130 | Soil |  |  |  | 2022 | 8 | 15 |
| REF-17 | 7087645 | 1688069 | Soil |  |  |  | 2022 | 8 | 15 |
| REF-18 | 7087716 | 1687905 | Soil |  |  |  | 2022 | 8 | 15 |
| REF-19 | 7087386 | 1688130 | Lingonberry |  |  |  | 2022 | 9 | 15 |
| REF-20 | 7149769 | 1679642 | Lingonberry |  |  |  | 2022 | 9 | 17 |
| REF-21 | 7161243 | 1688781 | Lingonberry |  |  |  | 2022 | 9 | 27 |
| REF-22 | 7149769 | 1679642 | Soil |  |  |  | 2022 | 9 | 17 |
| REF-23 | 7161243 | 1688781 | Soil |  |  |  | 2022 | 9 | 27 |
| Ungulate_1 | 7009728 | 1436437 | Roedeer | Male | Juvenile |  | 2021 | 10 | 2 |
| Ungulate_10 | 7008383 | 1431711 | Roedeer | Male | Adult |  | 2021 | 10 | 12 |
| Ungulate_11 | 7012050 | 1435304 | Roedeer | Female | Juvenile |  | 2021 | 10 | 2 |
| Ungulate_12 | 7012375 | 1432710 | Roedeer | Female | Juvenile |  | 2021 | 10 | 3 |
| Ungulate_13 | 7012305 | 1434639 | Roedeer | Female | Juvenile |  | 2021 | 10 | 9 |
| Ungulate_14 | 7011888 | 1432700 | Roedeer | Female | Juvenile |  | 2021 | 10 | 10 |
| Ungulate_15 | 7011043 | 1435858 | Moose | Male | Juvenile |  | 2021 | 10 | 26 |
| Ungulate_16 | 7008097 | 1436617 | Moose | Female | Juvenile |  | 2021 | 10 | 17 |
| Ungulate_17 | 7011818 | 1435290 | Moose | Male | Juvenile |  | 2021 | 10 | 6 |
| Ungulate_18 | 7011905 | 1435245 | Moose | Male | Adult |  | 2021 | 11 | 20 |
| Ungulate_2 | 7009728 | 1436437 | Roedeer | Female | Juvenile |  | 2021 | 10 | 2 |
| Ungulate_3 | 7009728 | 1436437 | Moose | Female | Adult |  | 2021 | 10 | 12 |
| Ungulate_4 | 7008055 | 1436352 | Moose | Female | Juvenile |  | 2021 | 10 | 23 |
| Ungulate_5 | 7011611 | 1433589 | Moose | Male | Juvenile |  | 2021 | 9 | 18 |
| Ungulate_6 | 7010125 | 1432387 | Roedeer | Female | Juvenile |  | 2022 | 1 | 16 |
| Ungulate_7 | 7011228 | 1433132 | Roedeer | Male | Juvenile |  | 2022 | 1 | 21 |
| Ungulate_8 | 7011587 | 1434517 | Moose |  | Juvenile |  | 2021 | 12 | 16 |
| Ungulate_9 | 7007814 | 1432066 | Roedeer | Male | Juvenile |  | 2021 | 10 | 2 |
| VolePool_1 | 7009593 | 1431867 | Bank vole | Female | Juvenile | 15.6 | 2022 | 7 | 13 |
| VolePool_1 | 7009507 | 1431655 | Bank vole | Female | Juvenile | 11.1 | 2022 | 8 | 11 |
| VolePool_1 | 7009593 | 1431867 | Bank vole | Female | Juvenile | 14.2 | 2022 | 9 | 13 |
| VolePool_10 | 7009738 | 1431674 | Bank vole | Male | Adult | 24.9 | 2022 | 7 | 13 |
| VolePool_10 | 7009593 | 1431867 | Bank vole | Male | Adult | 23 | 2022 | 7 | 13 |
| VolePool_10 | 7009593 | 1431867 | Bank vole | Male | Adult | 18.2 | 2022 | 7 | 13 |
| VolePool_11 | 7009593 | 1431867 | Bank vole | Male | Adult | 17.7 | 2022 | 7 | 14 |
| VolePool_11 | 7009507 | 1431655 | Bank vole | Male | Adult | 26 | 2022 | 7 | 14 |
| VolePool_11 | 7009507 | 1431655 | Bank vole | Male | Adult | 22.6 | 2022 | 8 | 11 |
| VolePool_12 | 7009593 | 1431867 | Bank vole | Male | Adult | 17.6 | 2022 | 8 | 12 |
| VolePool_12 | 7009507 | 1431655 | Bank vole | Male | Adult | 20.5 | 2022 | 8 | 12 |
| VolePool_2 | 7009738 | 1431674 | Bank vole | Female | Juvenile | 13.9 | 2022 | 9 | 13 |
| VolePool_2 | 7009738 | 1431674 | Bank vole | Female | Juvenile | 13.6 | 2022 | 9 | 13 |
| VolePool_2 | 7009738 | 1431674 | Bank vole | Female | Juvenile | 13.8 | 2022 | 9 | 13 |
| VolePool_3 | 7009738 | 1431674 | Bank vole | Female | Juvenile | 11.4 | 2022 | 9 | 13 |
| VolePool_3 | 7009507 | 1431655 | Bank vole | Female | Juvenile | 14.1 | 2022 | 9 | 13 |
| VolePool_3 | 7009738 | 1431674 | Bank vole | Female | Juvenile | 14.1 | 2022 | 9 | 14 |
| VolePool_4 | 7009738 | 1431674 | Bank vole | Female | Adult | 27.7 | 2022 | 7 | 14 |
| VolePool_4 | 7009738 | 1431674 | Bank vole | Female | Adult | 24.8 | 2022 | 7 | 14 |
| VolePool_4 | 7009738 | 1431674 | Bank vole | Female | Adult | 27.3 | 2022 | 8 | 11 |
| VolePool_5 | 7009507 | 1431655 | Bank vole | Female | Adult | 19 | 2022 | 8 | 11 |
| VolePool_5 | 7009507 | 1431655 | Bank vole | Female | Adult | 17.2 | 2022 | 9 | 14 |
| VolePool_6 | 7009738 | 1431674 | Bank vole | Male | Juvenile | 12.4 | 2022 | 8 | 12 |
| VolePool_6 | 7009507 | 1431655 | Bank vole | Male | Juvenile | 17.2 | 2022 | 8 | 12 |
| VolePool_6 | 7009507 | 1431655 | Bank vole | Male | Juvenile | 11.2 | 2022 | 9 | 13 |
| VolePool_7 | 7009507 | 1431655 | Bank vole | Male | Juvenile | 15.8 | 2022 | 9 | 13 |
| VolePool_7 | 7009507 | 1431655 | Bank vole | Male | Juvenile | 14.2 | 2022 | 9 | 13 |
| VolePool_7 | 7009593 | 1431867 | Bank vole | Male | Juvenile | 17.3 | 2022 | 9 | 14 |
| VolePool_8 | 7009738 | 1431674 | Bank vole | Male | Juvenile | 14.1 | 2022 | 9 | 14 |
| VolePool_8 | 7009738 | 1431674 | Bank vole | Male | Juvenile | 10.6 | 2022 | 9 | 14 |
| VolePool_9 | 7009507 | 1431655 | Bank vole | Male | Juvenile | 15.8 | 2022 | 9 | 14 |
| VolePool_9 | 7009507 | 1431655 | Bank vole | Male | Juvenile | 17.9 | 2022 | 9 | 14 |

**Appendix 2.**
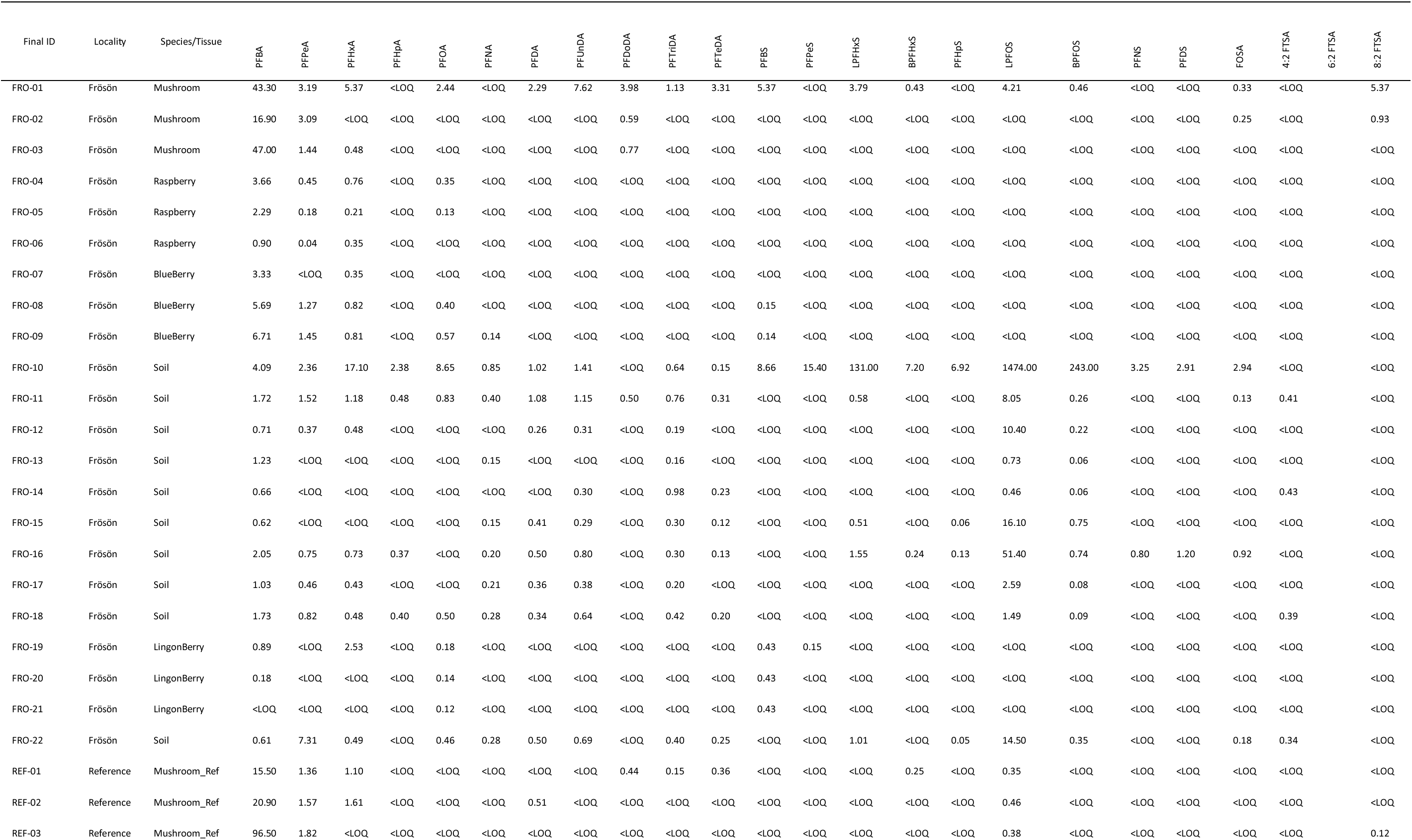
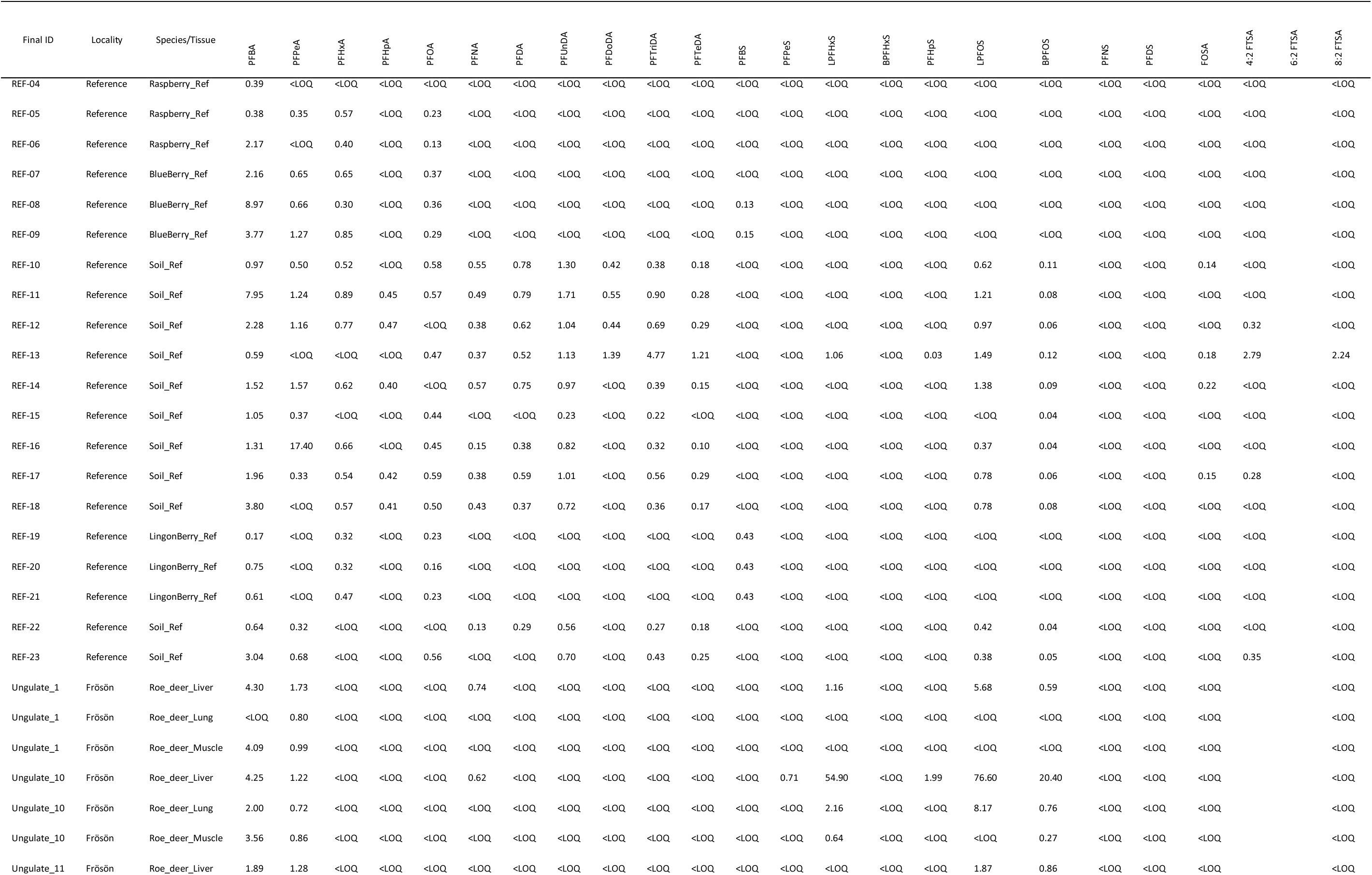
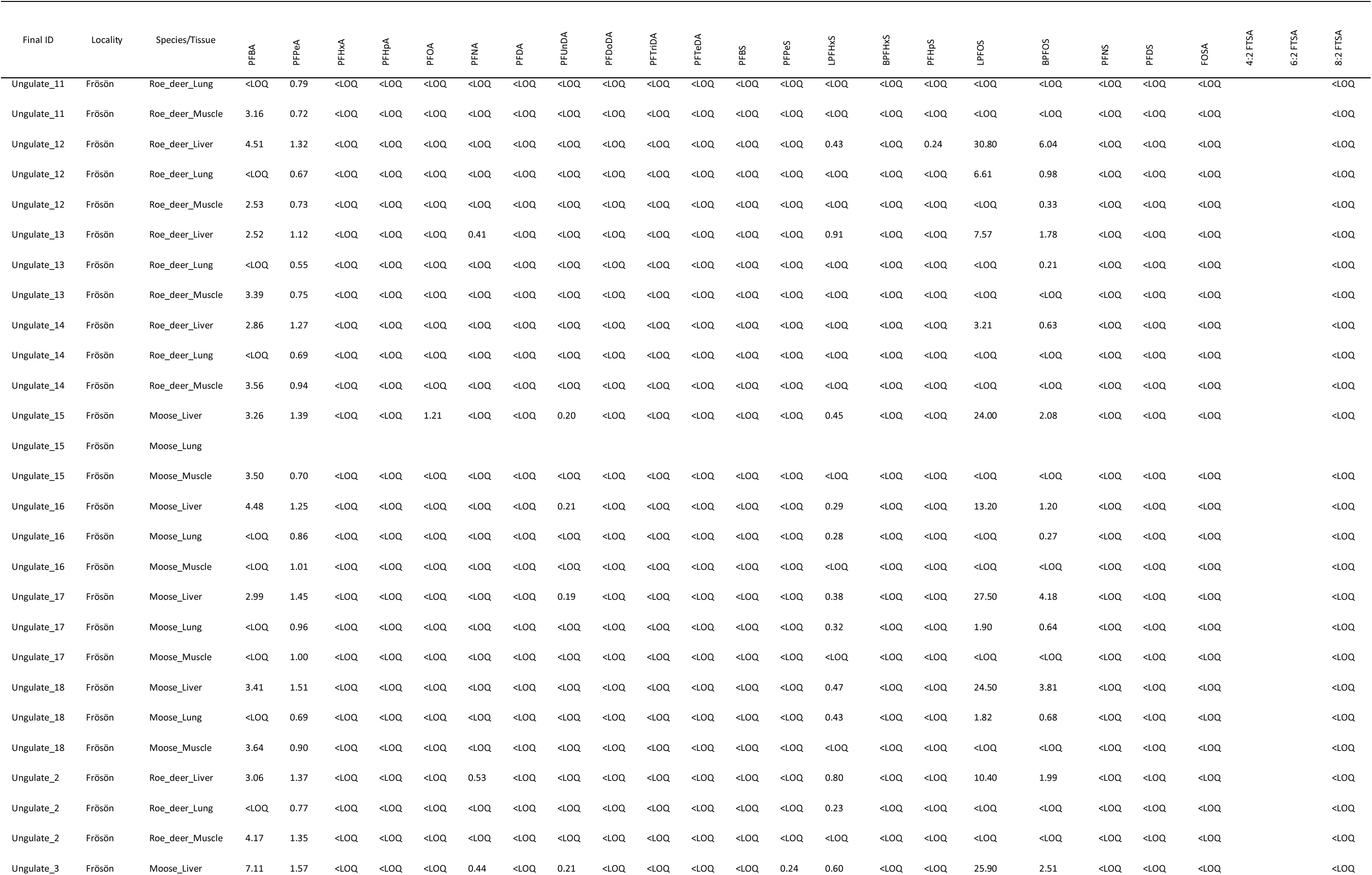
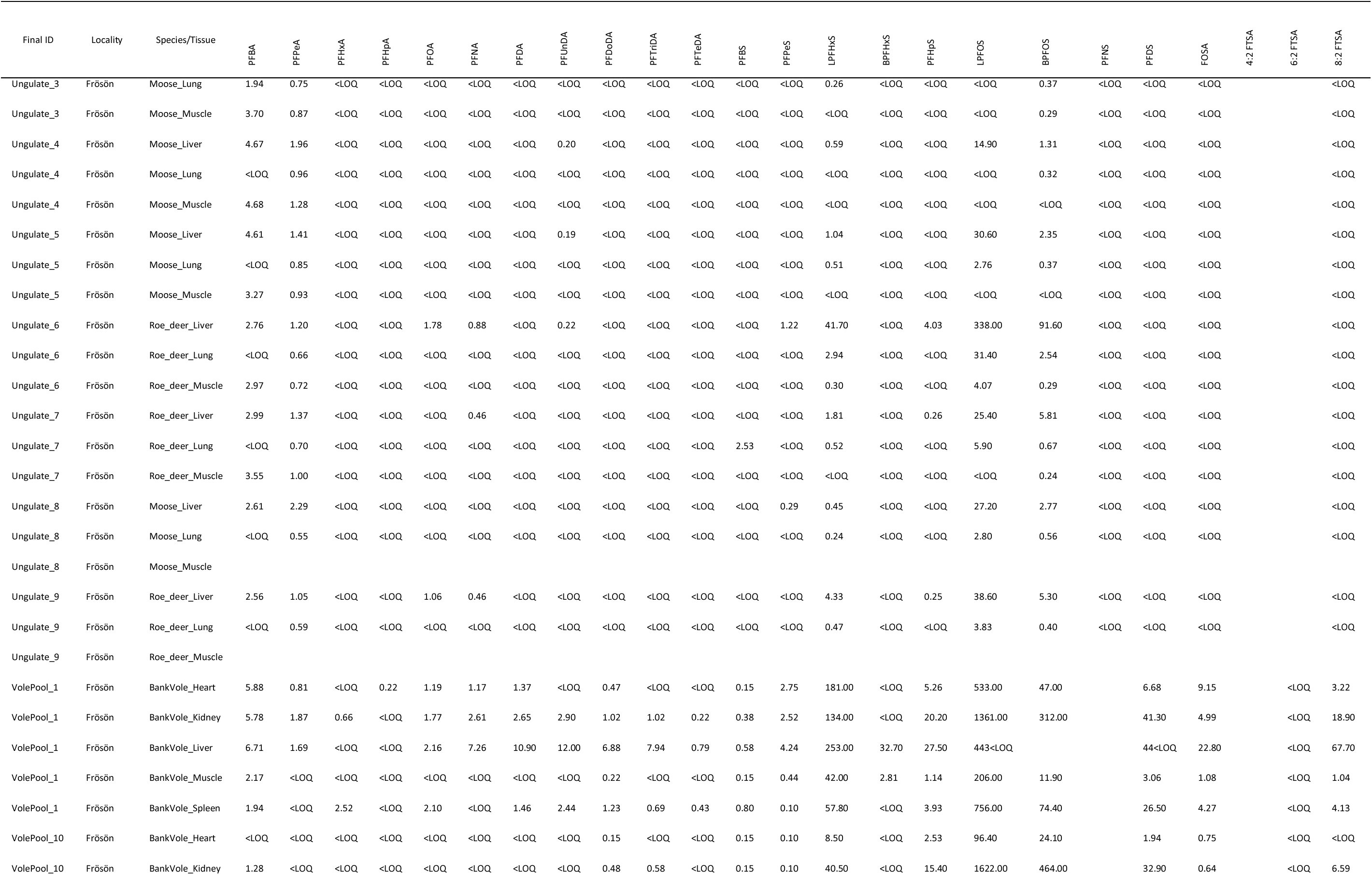
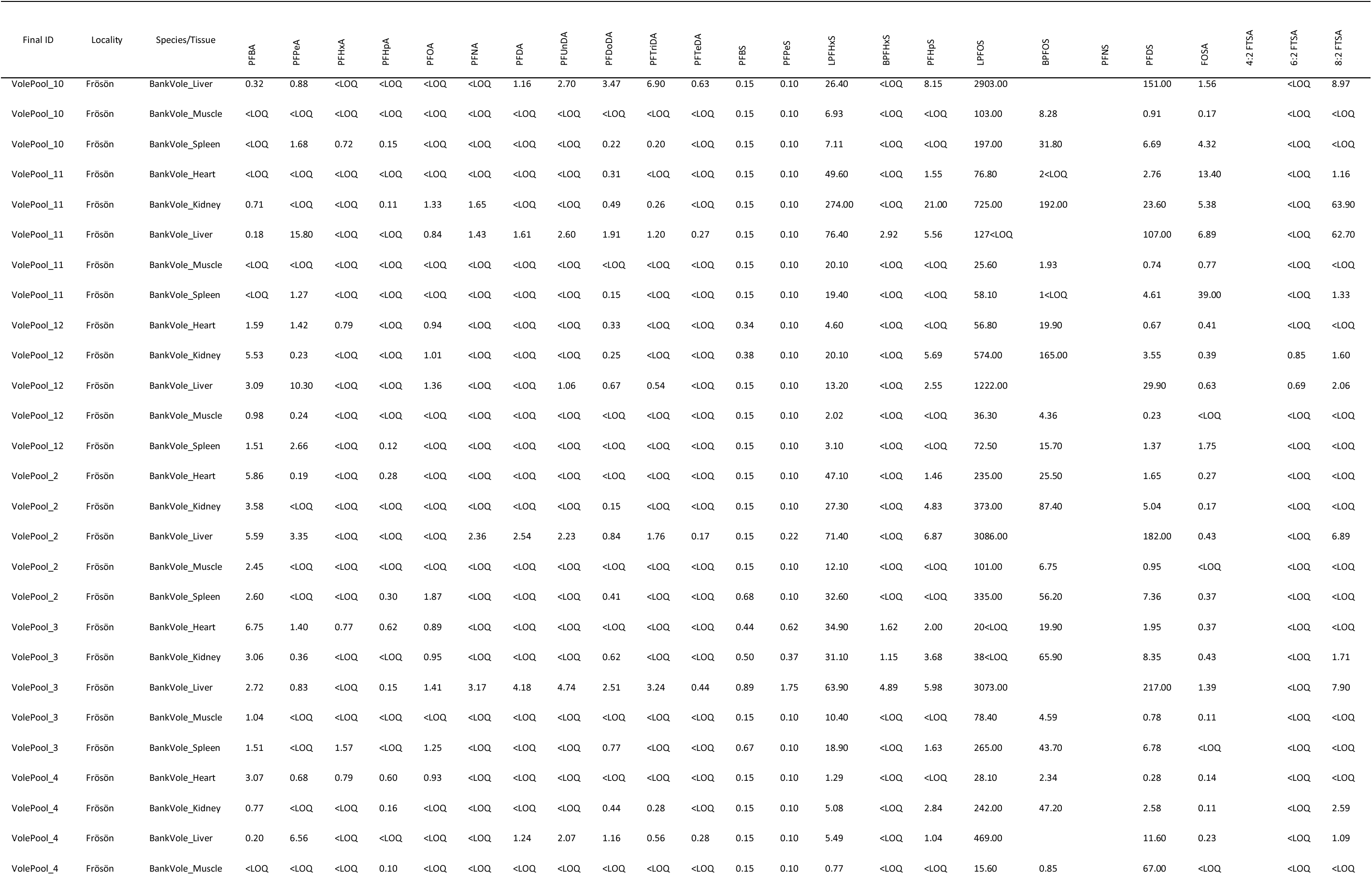
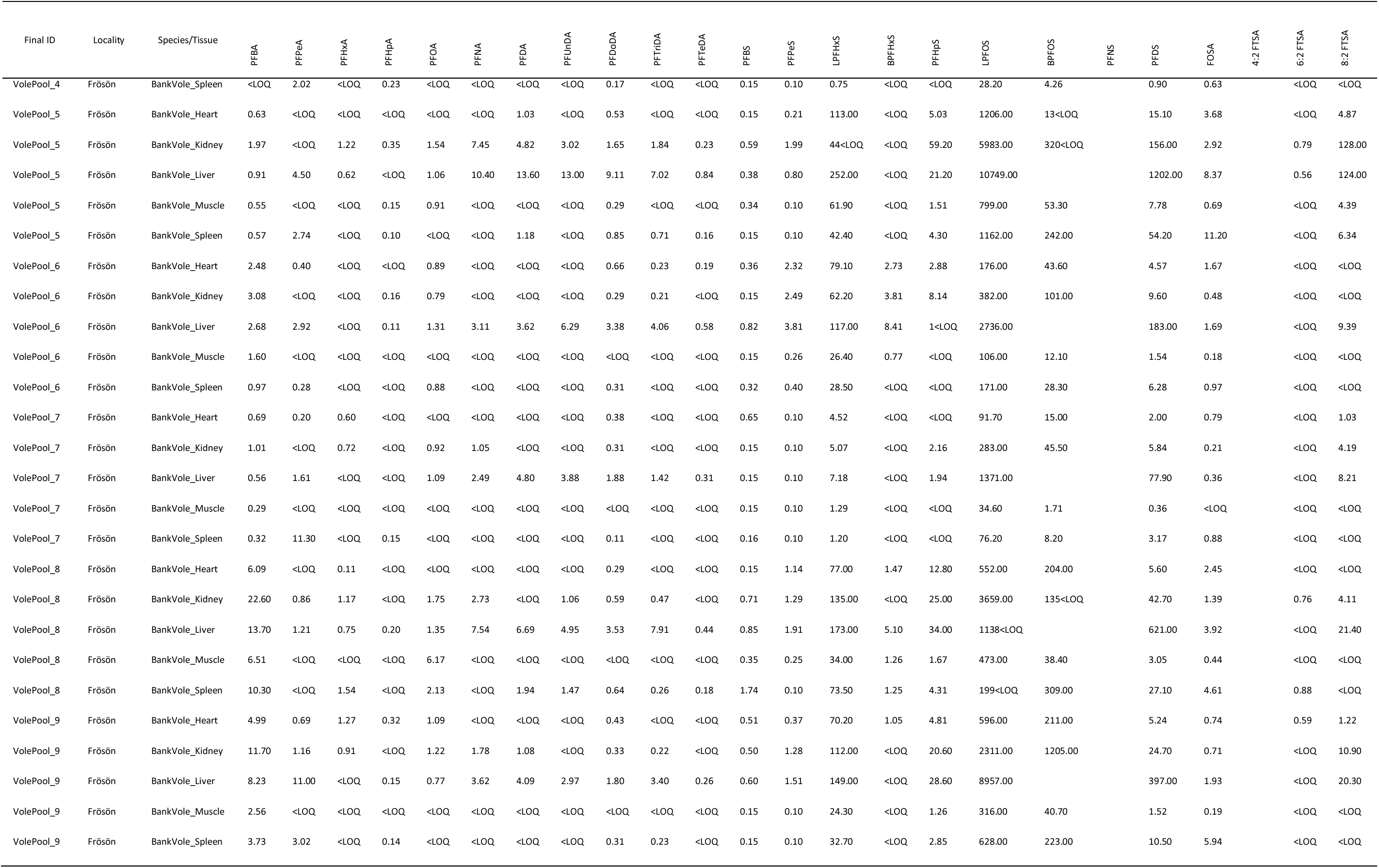
Raw data for 22 PFAS in soil and biota on the island of Frösön and in reference areas near Umeå, northern Sweden. For wildlife (bank vole, roedeer, and moose), concentrations are given in ng/g dw, for other samples (soil, mushroom, and berries) in ng/g ww.

**Appendix 3.**
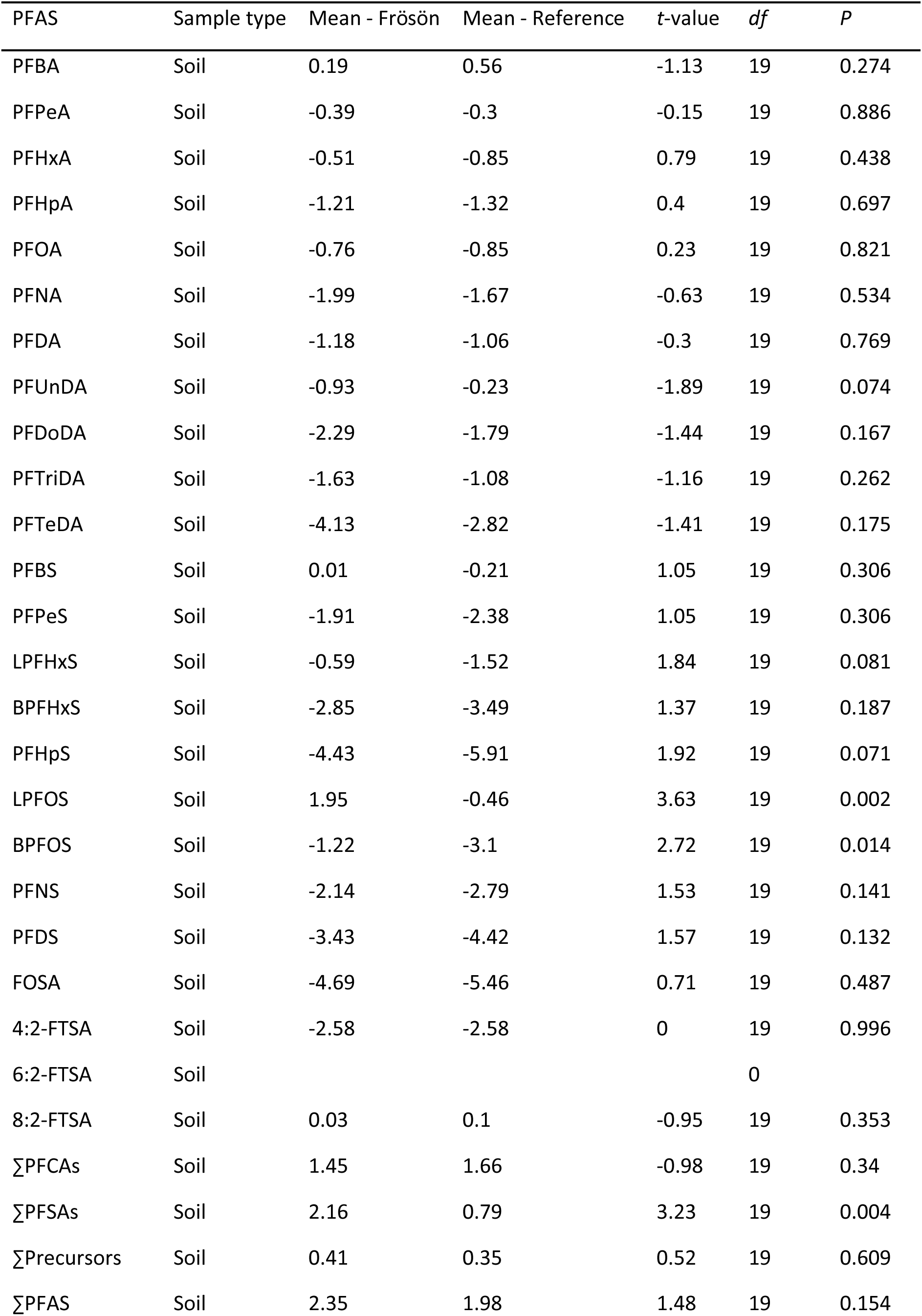
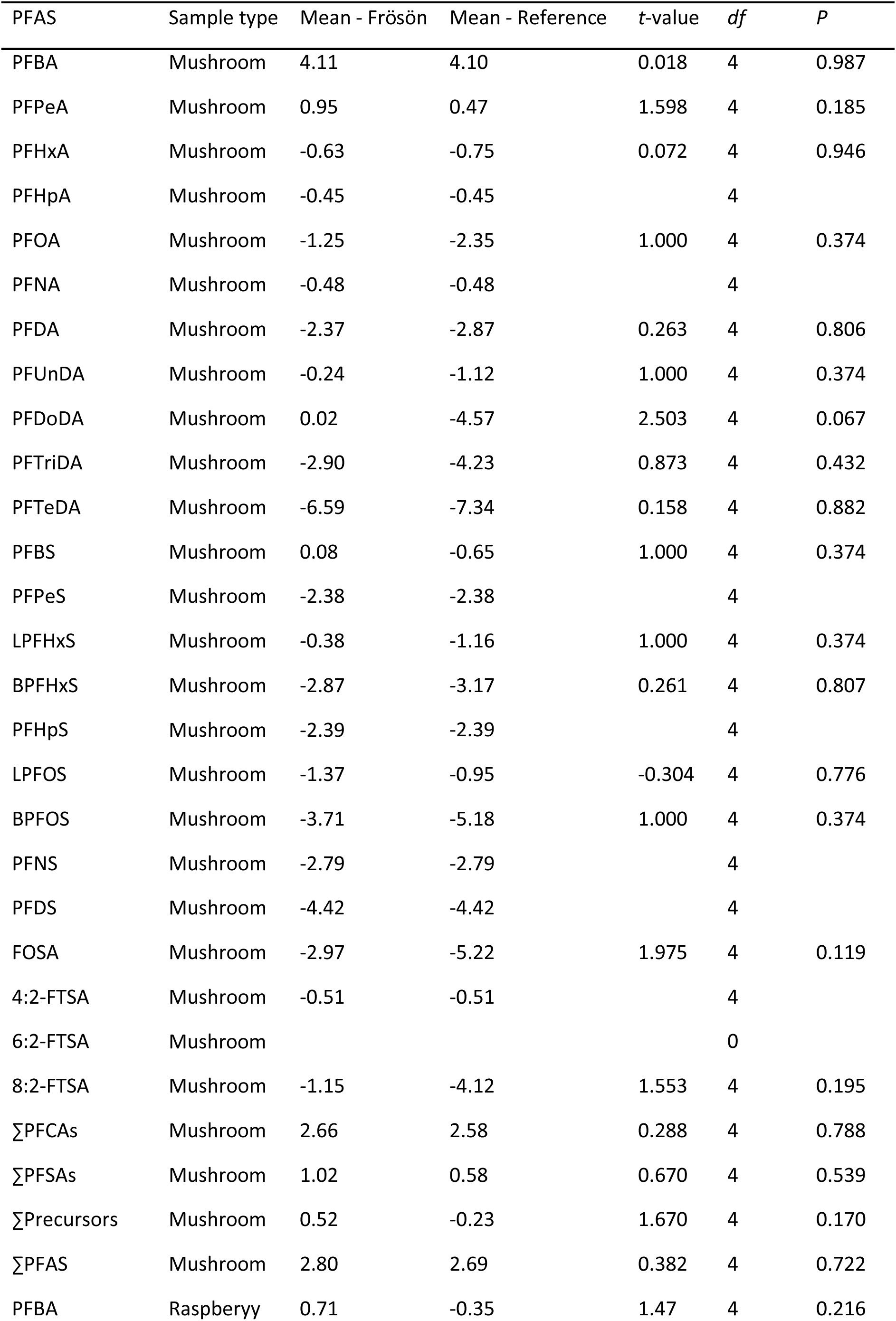
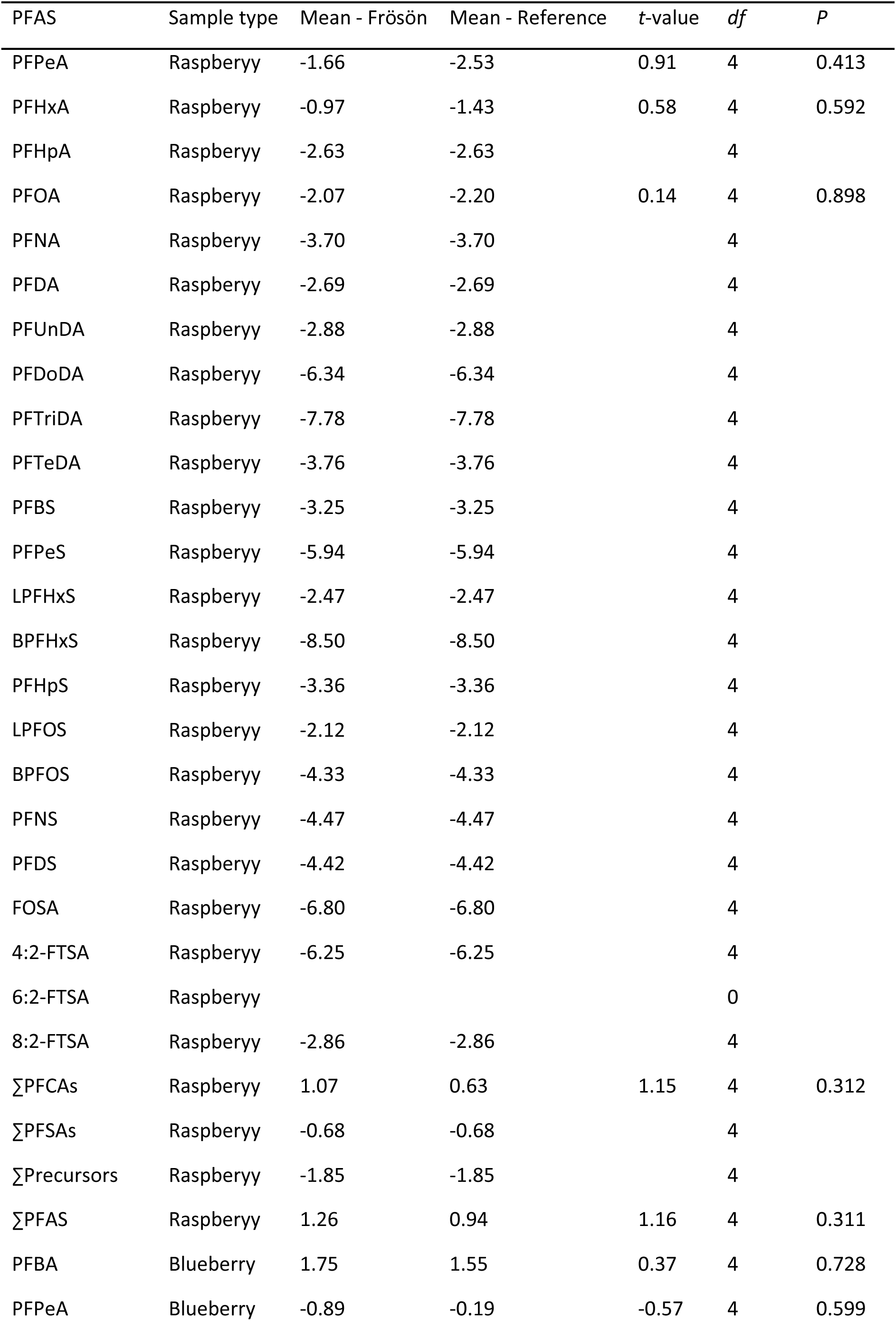
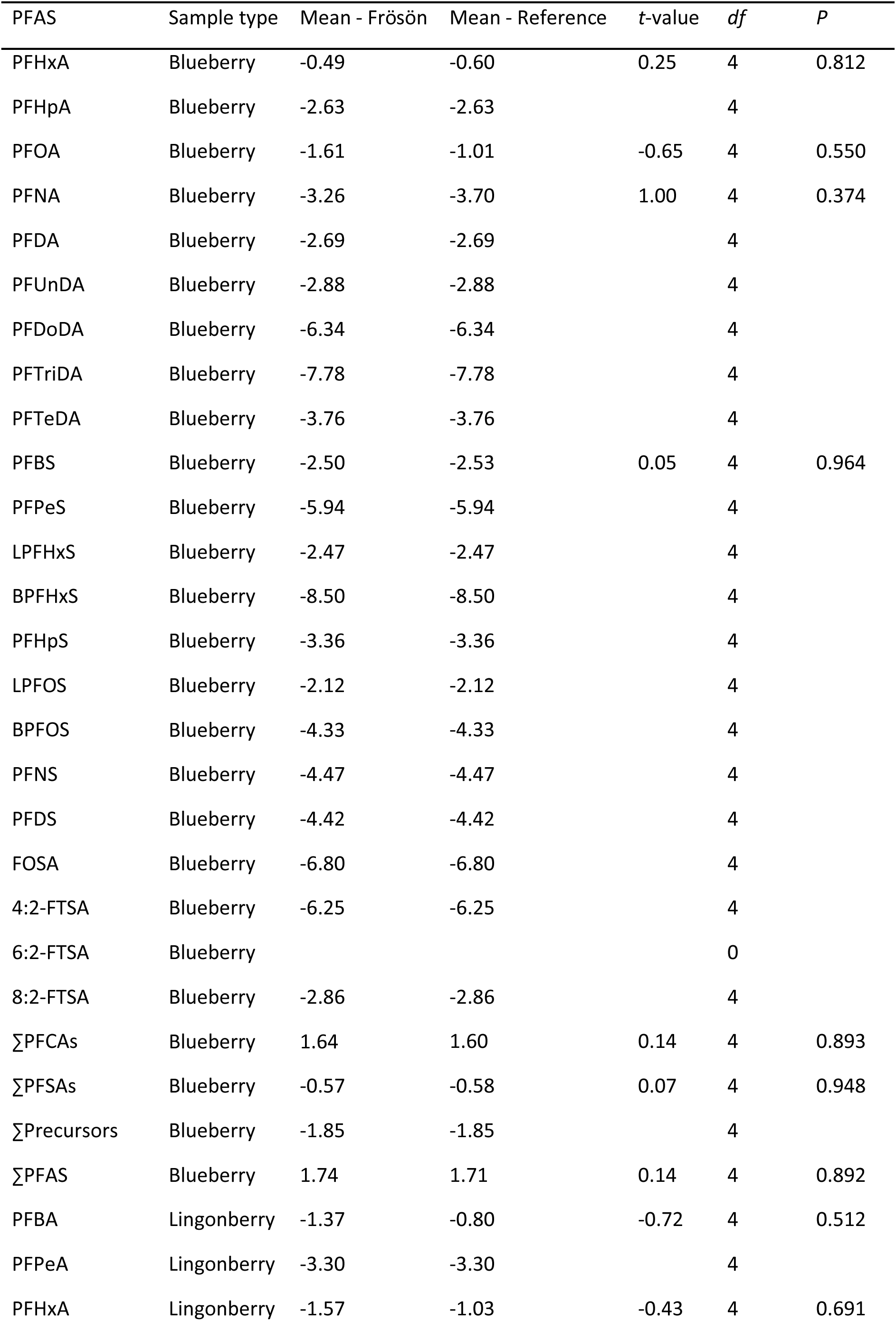
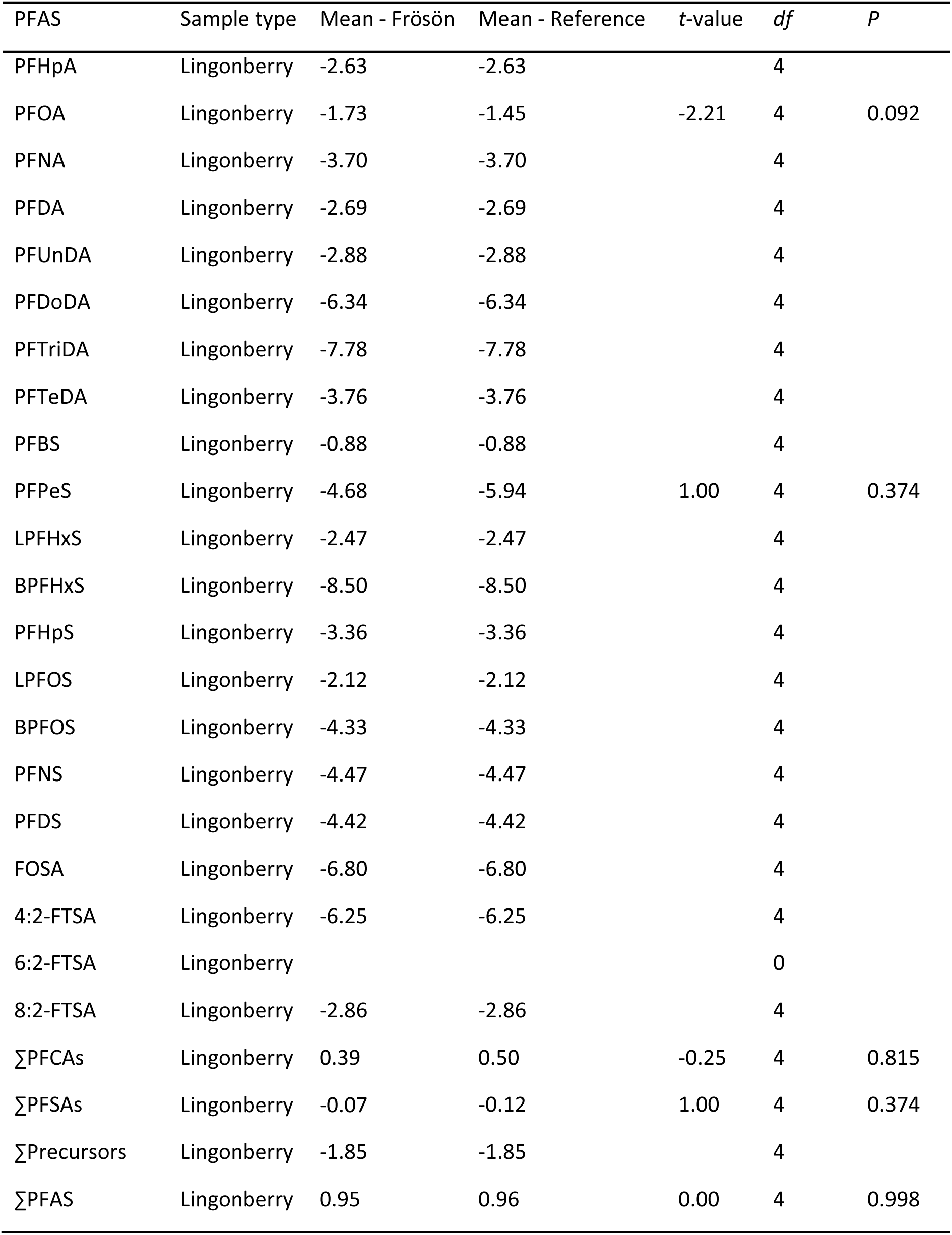
Results of t-tests between PFAS concentrations on the Island of Frösön and the reference localities near Umeå, northern Sweden.

**Appendix 4.**
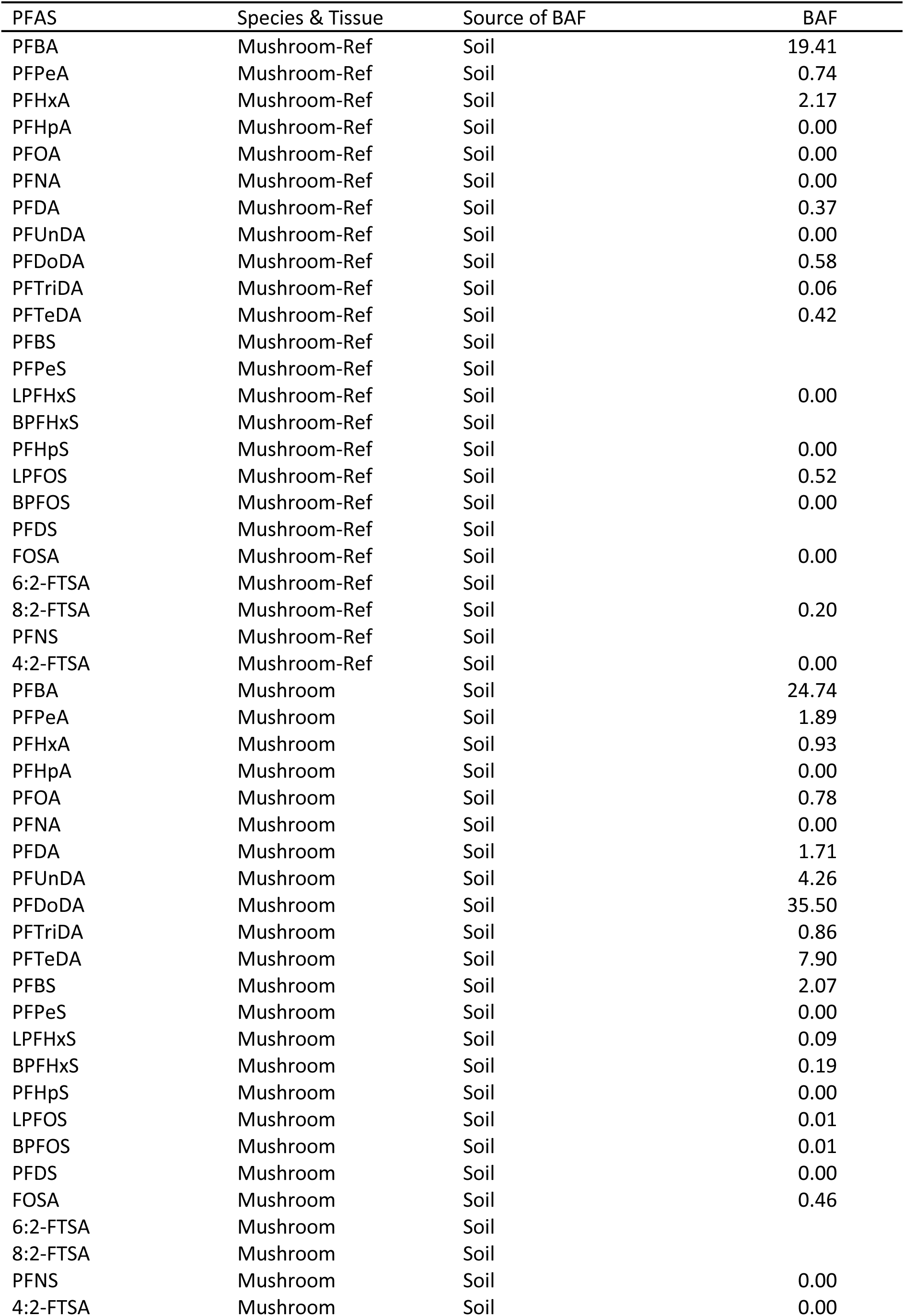
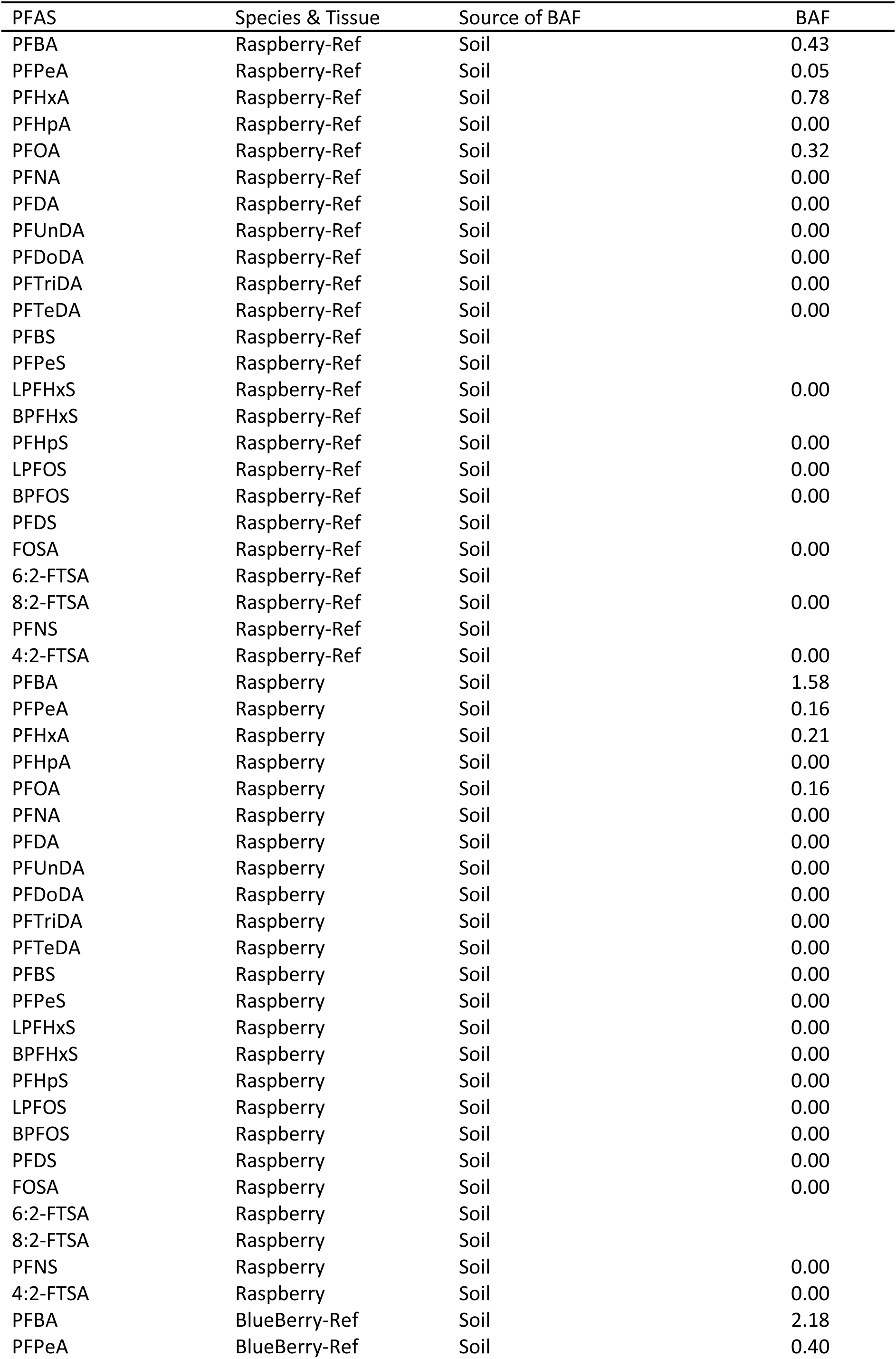
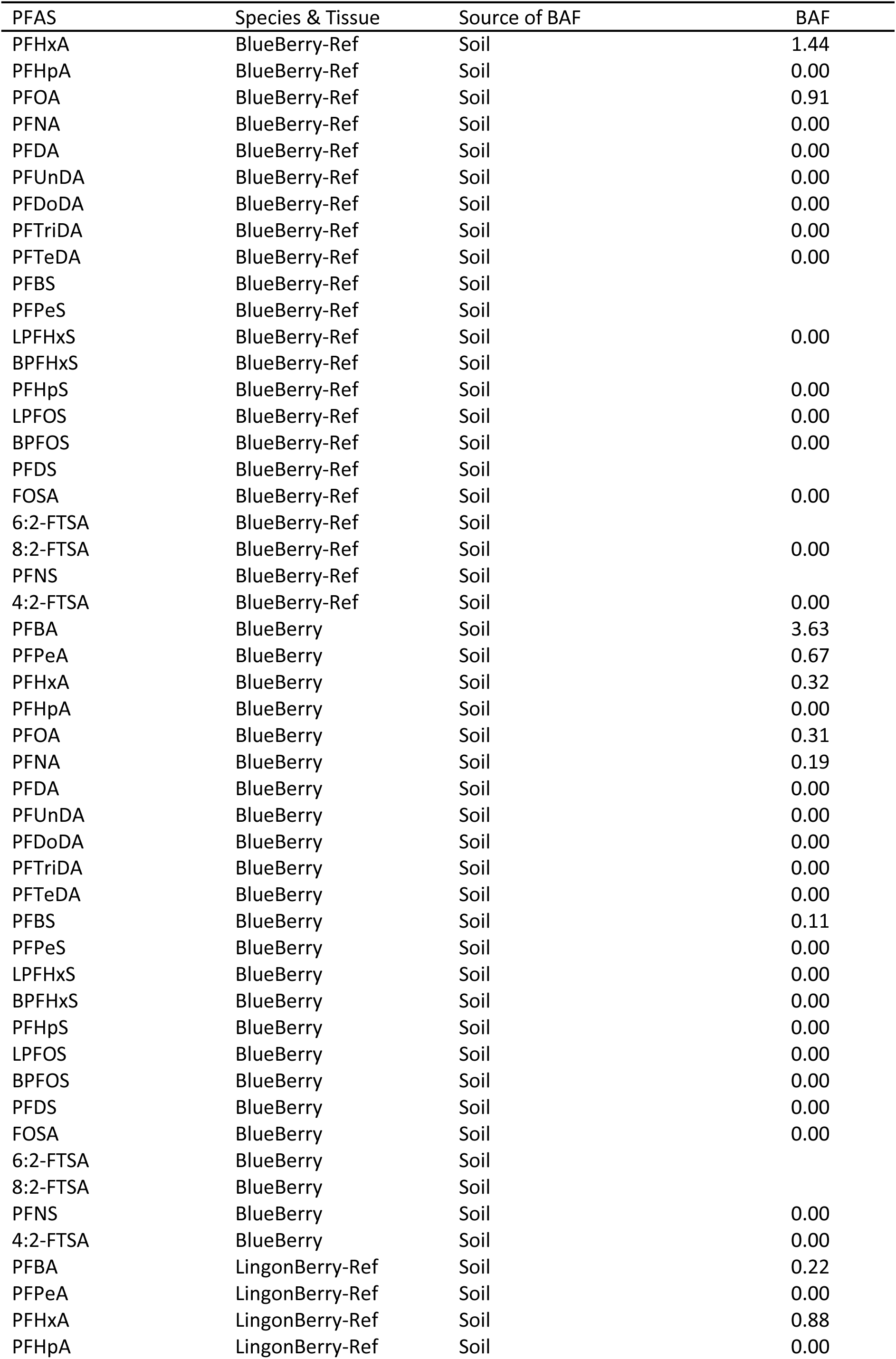
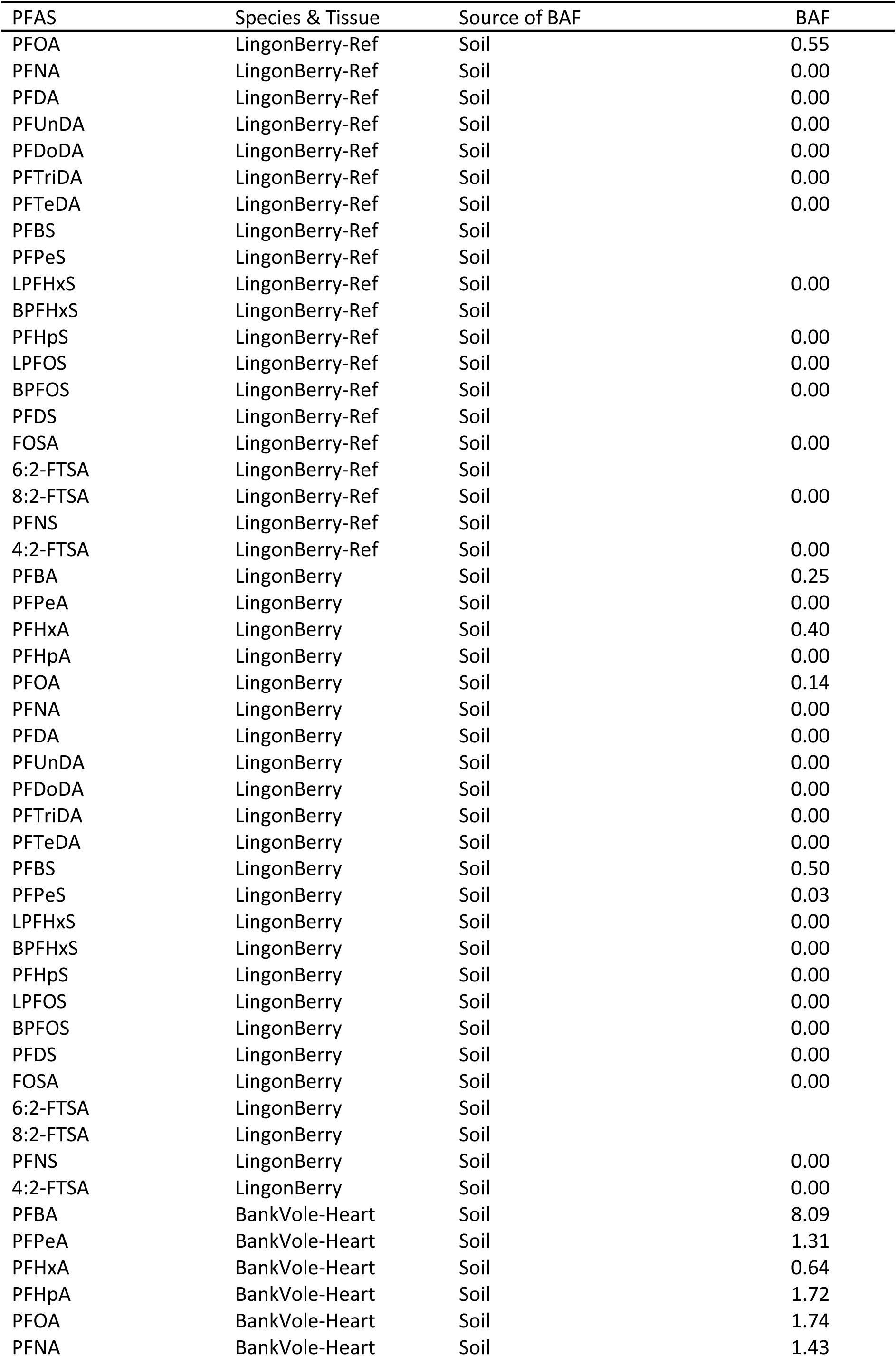
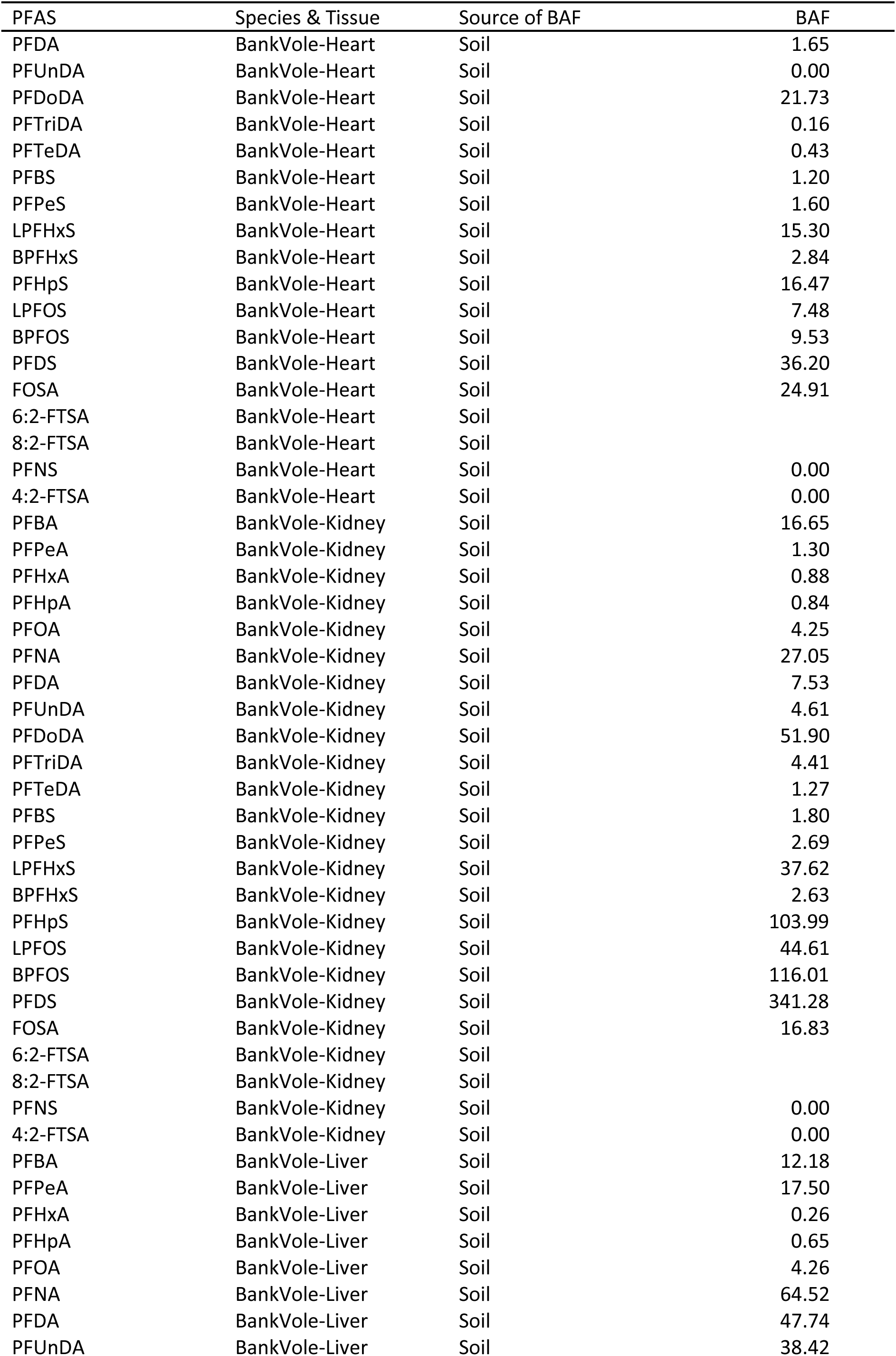
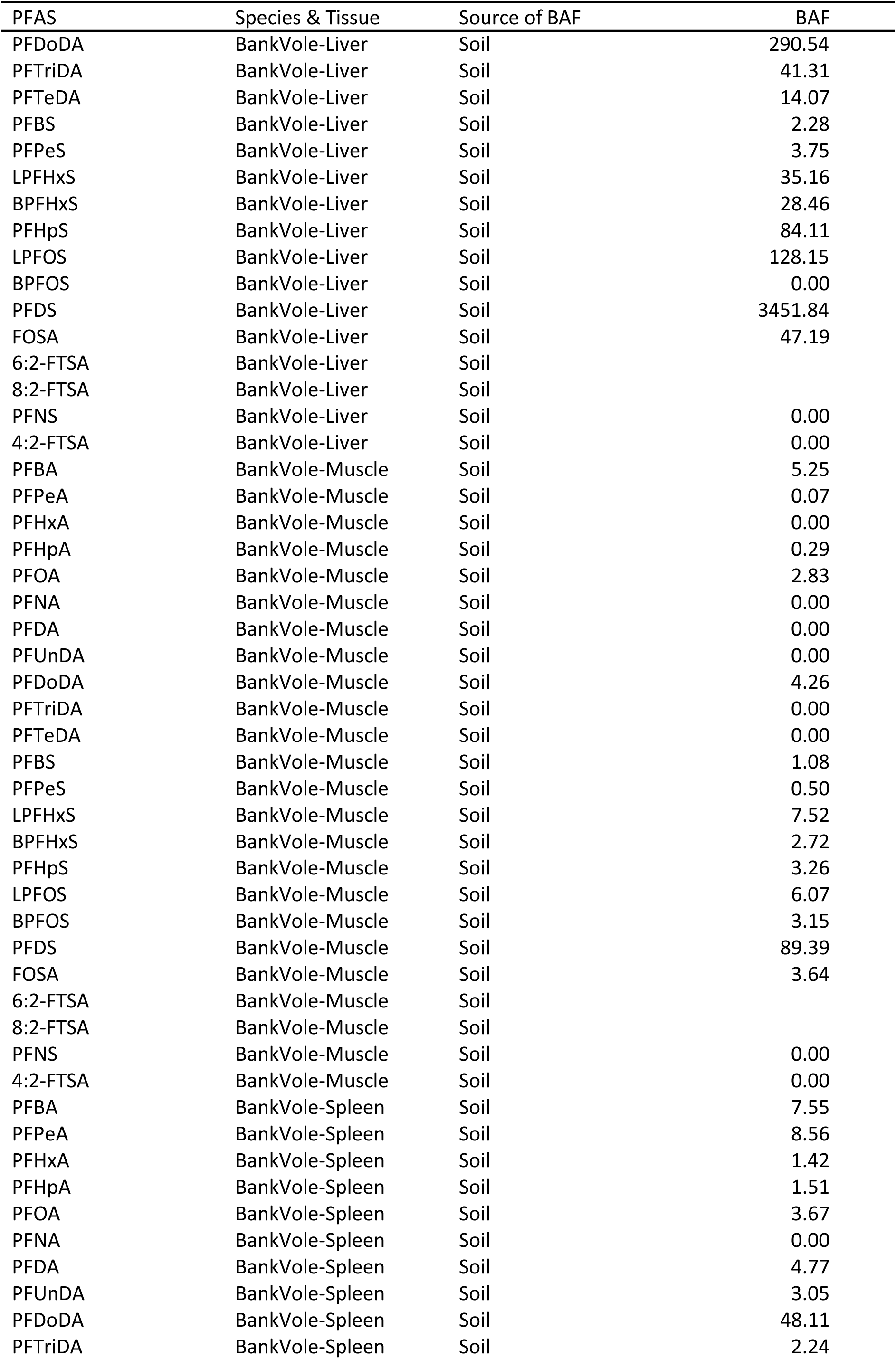
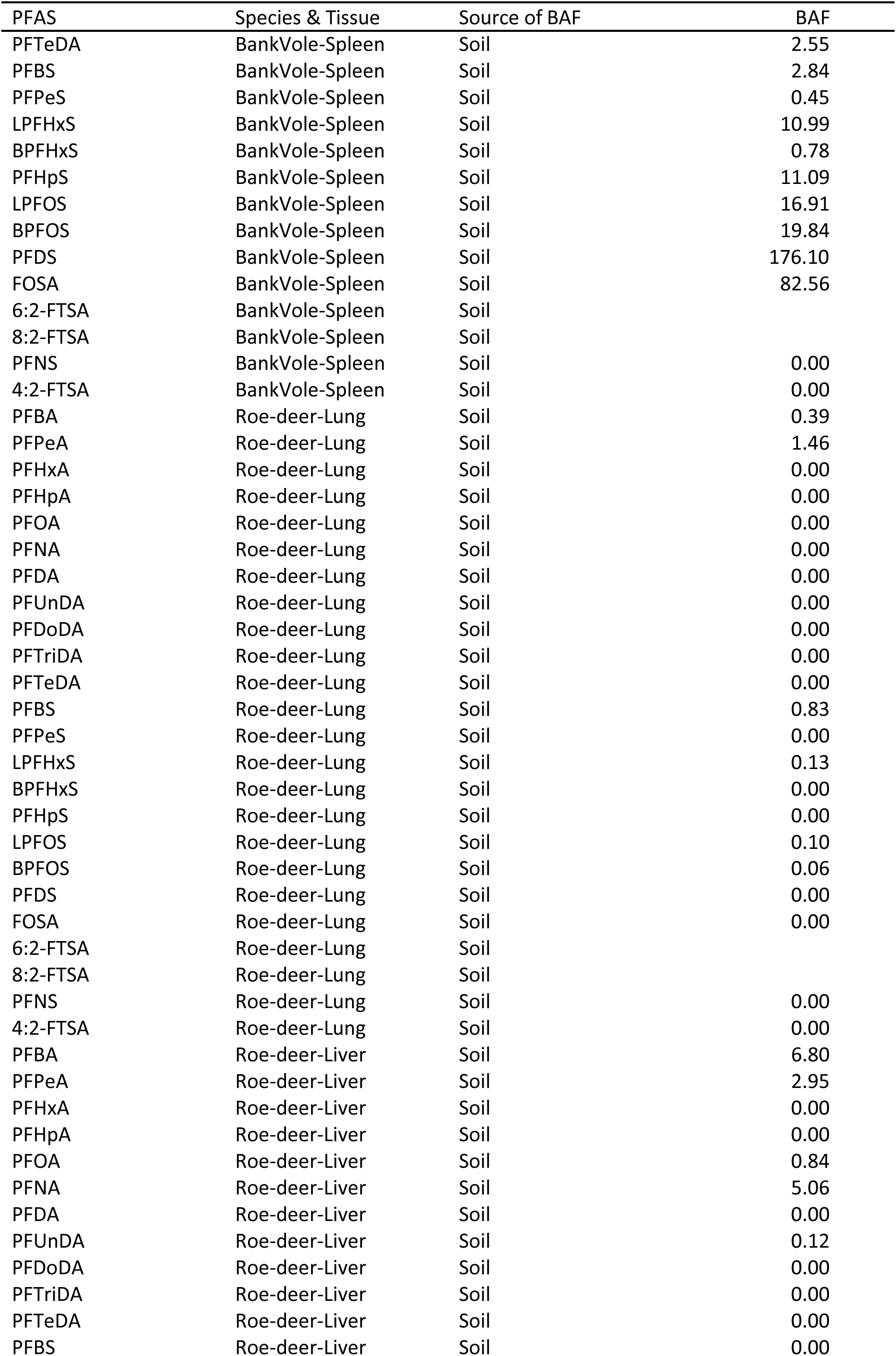
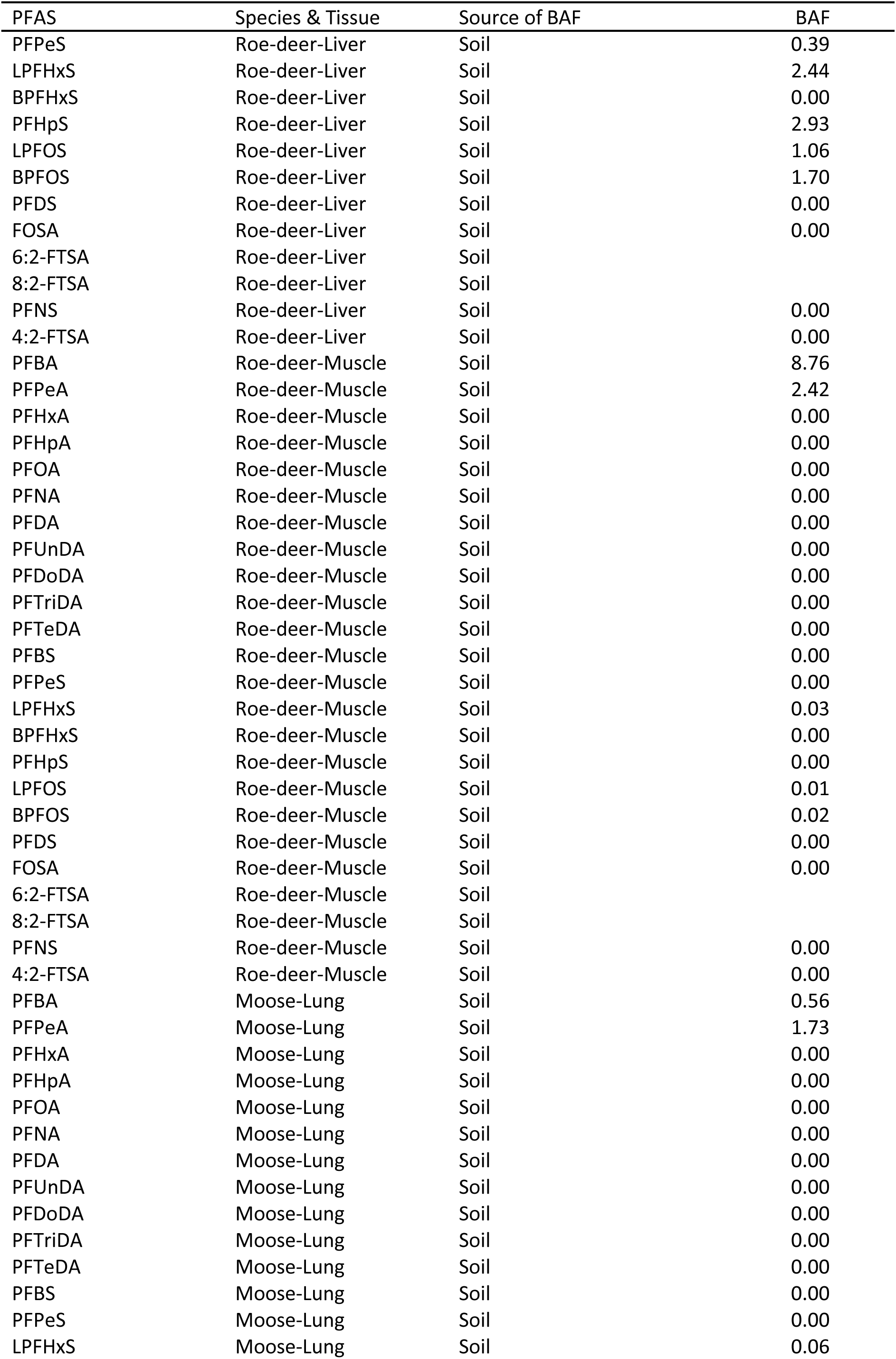
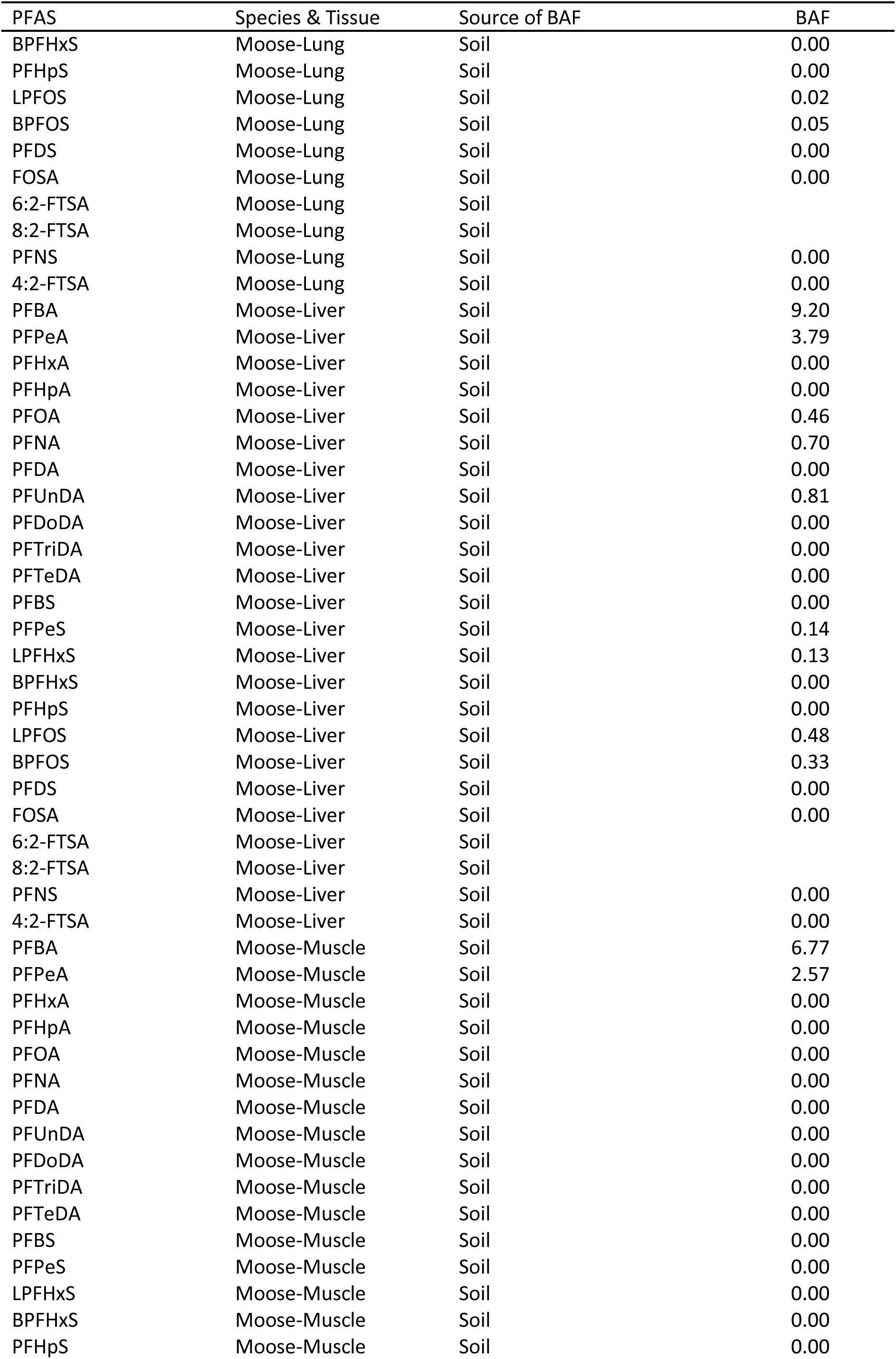
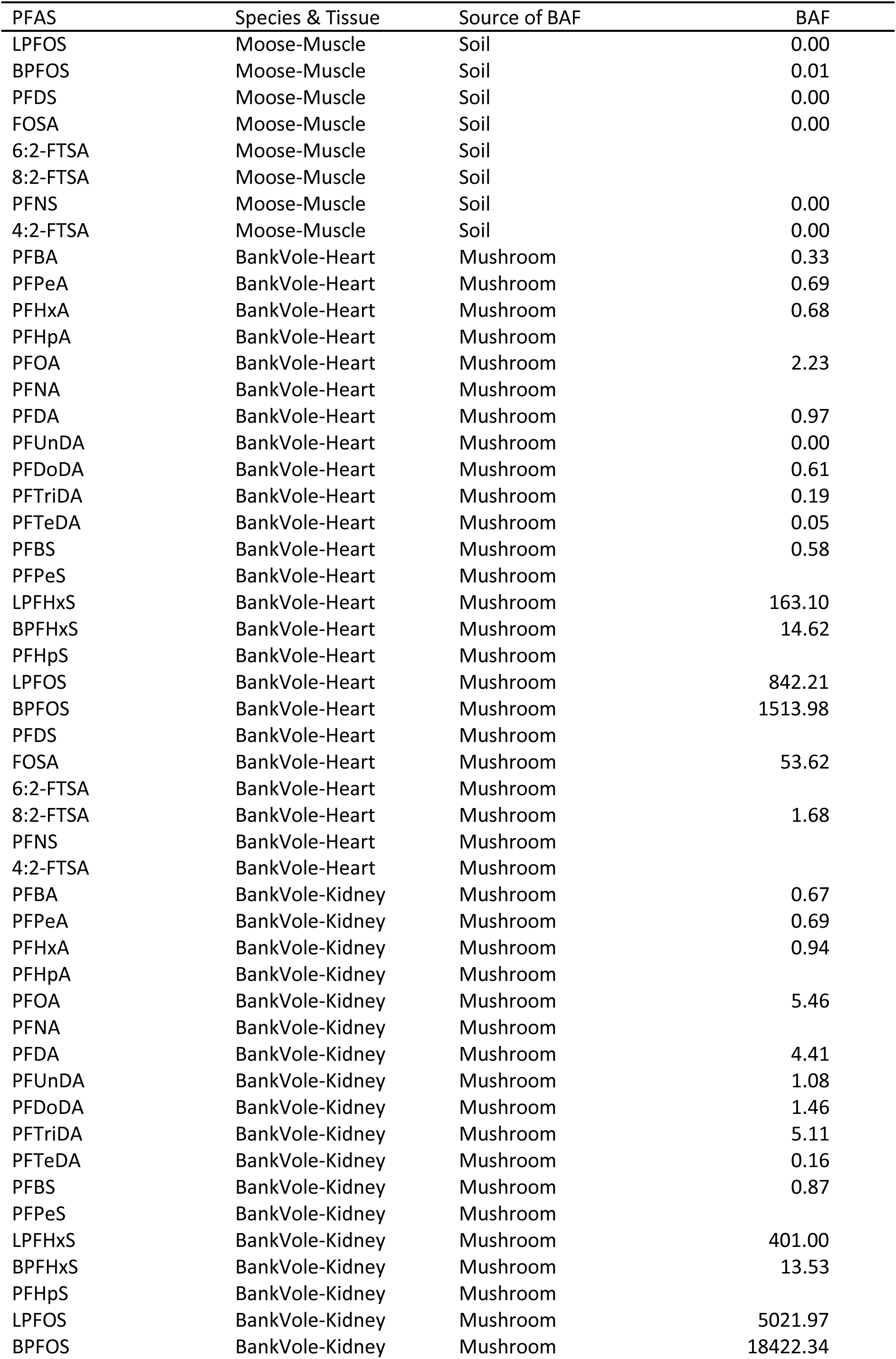
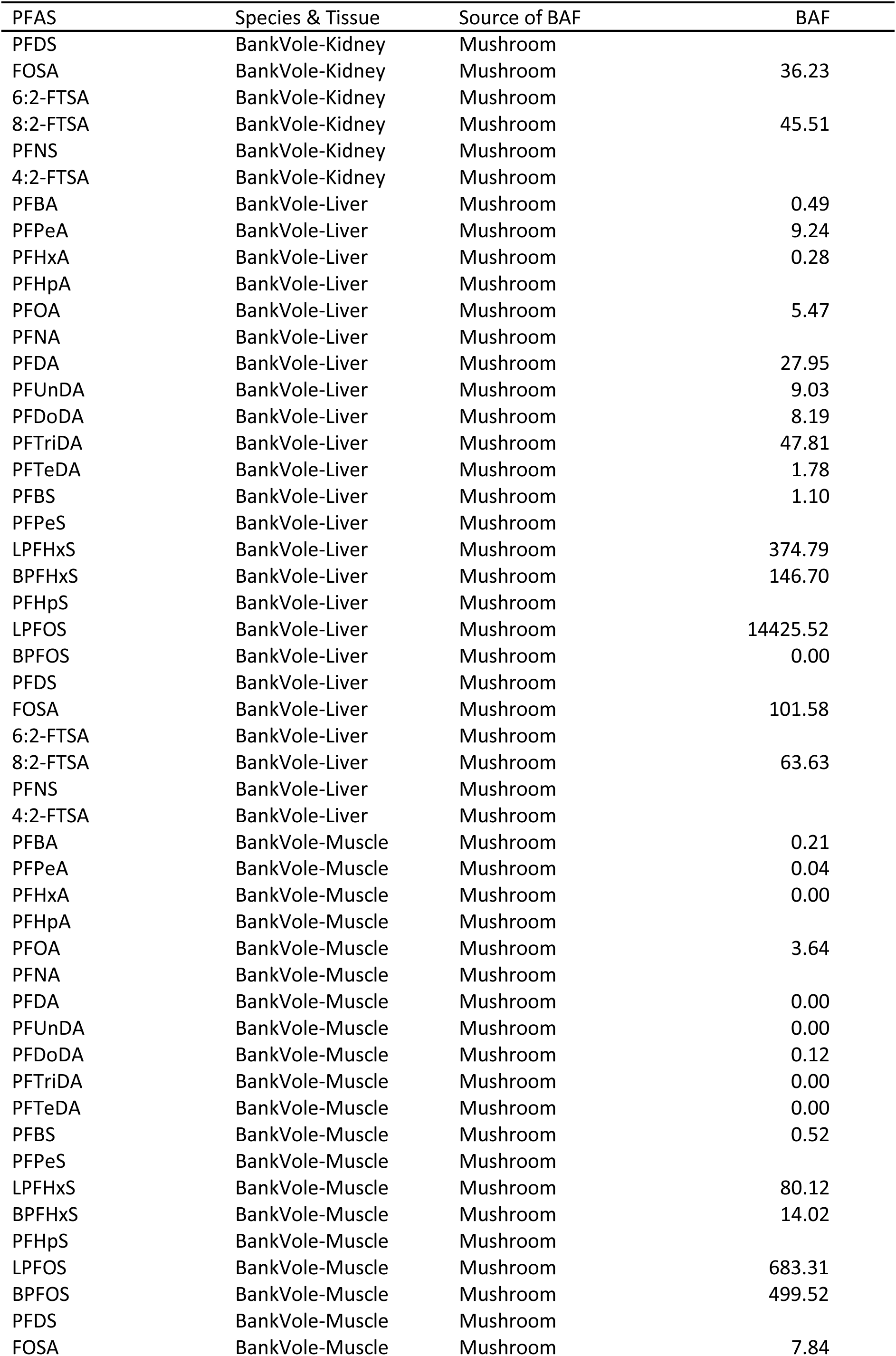
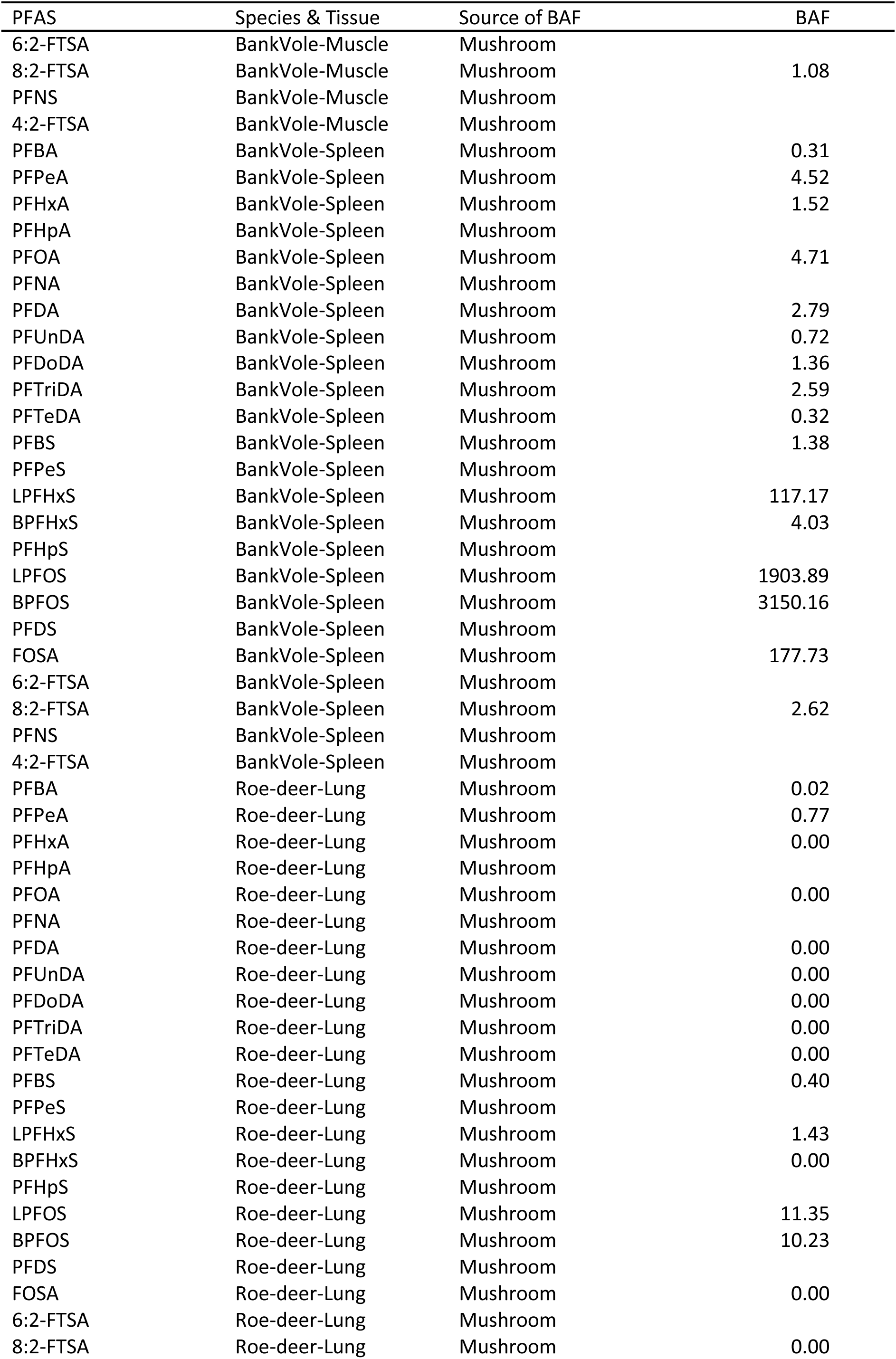
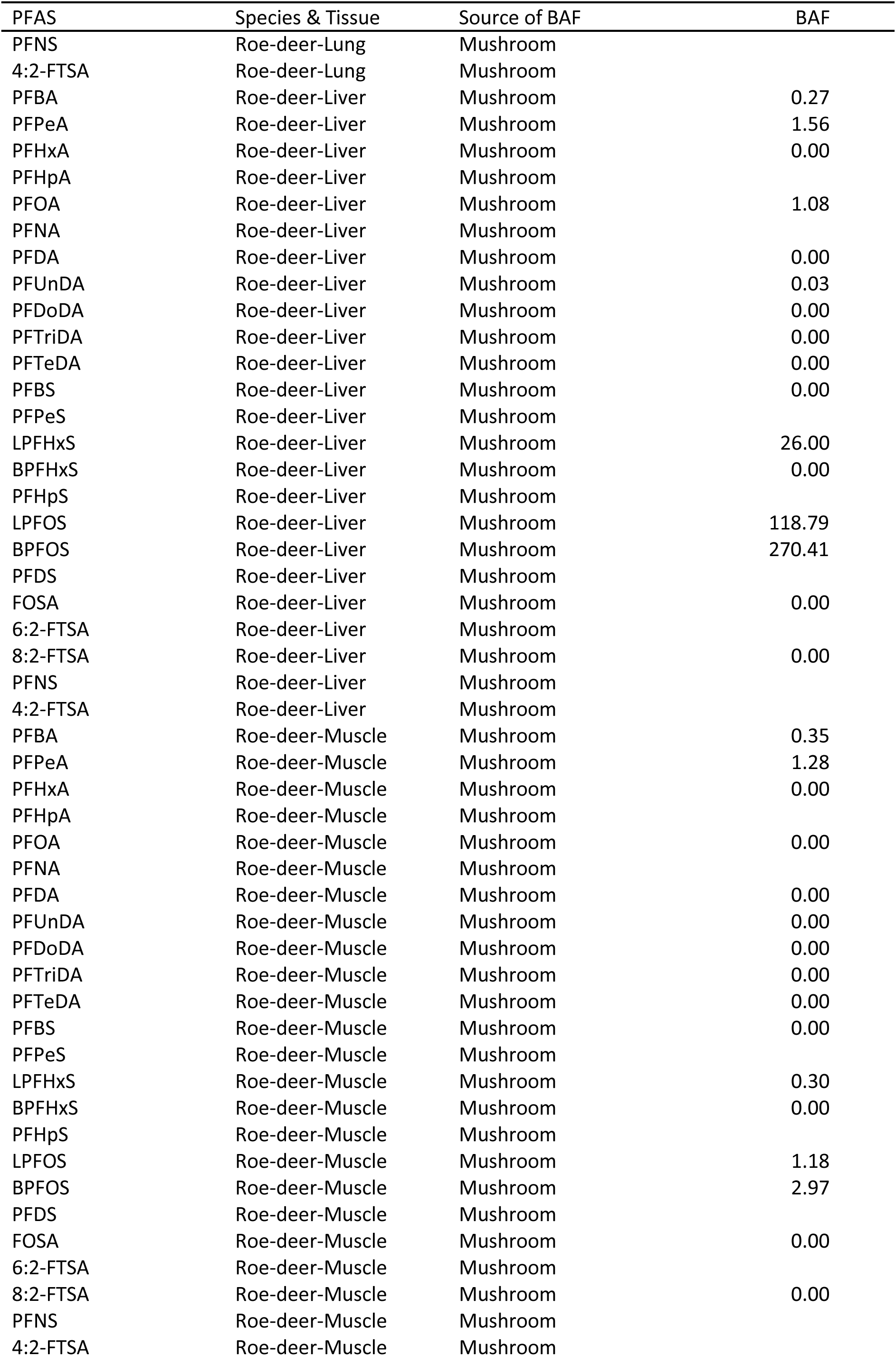
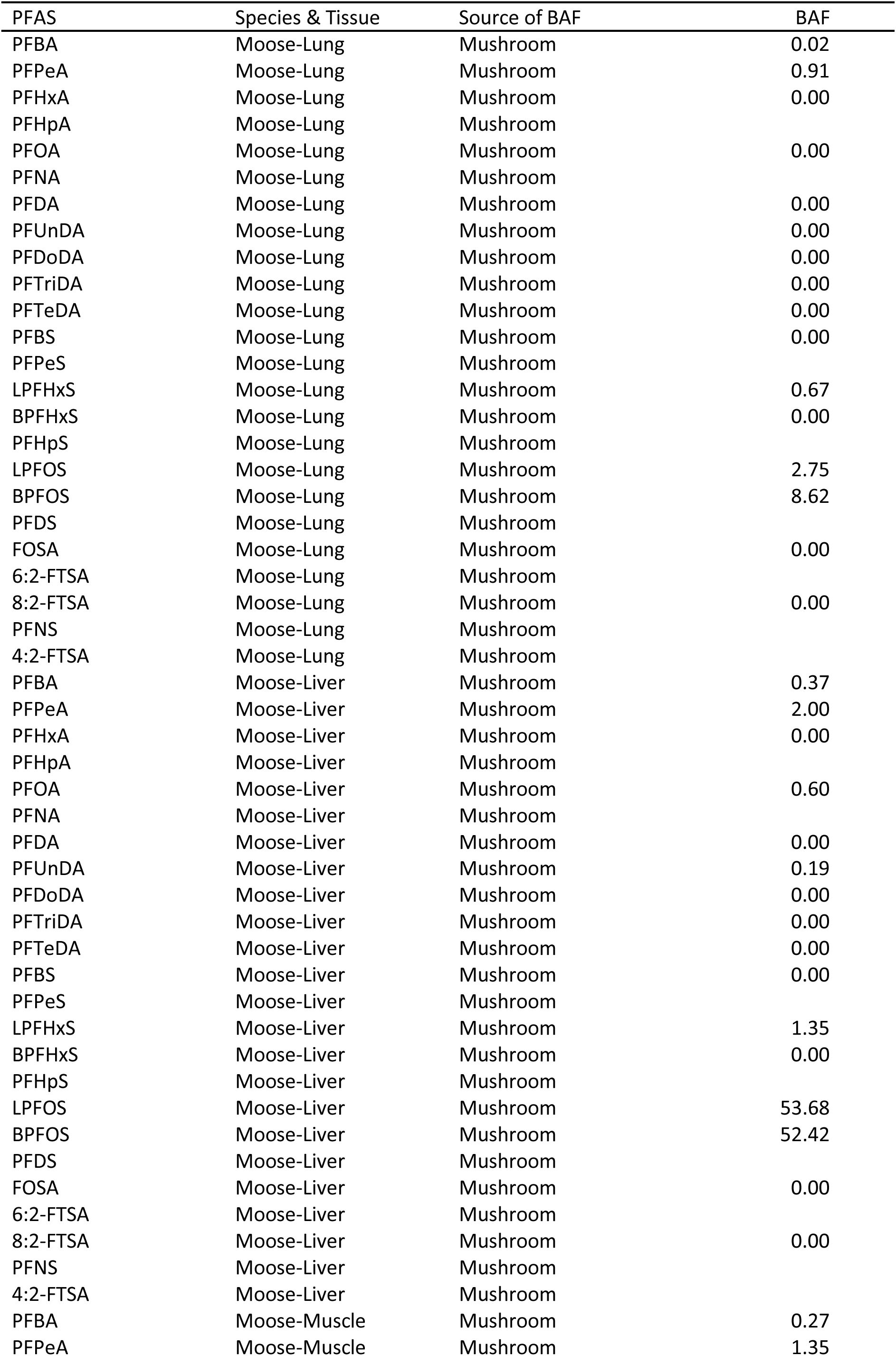
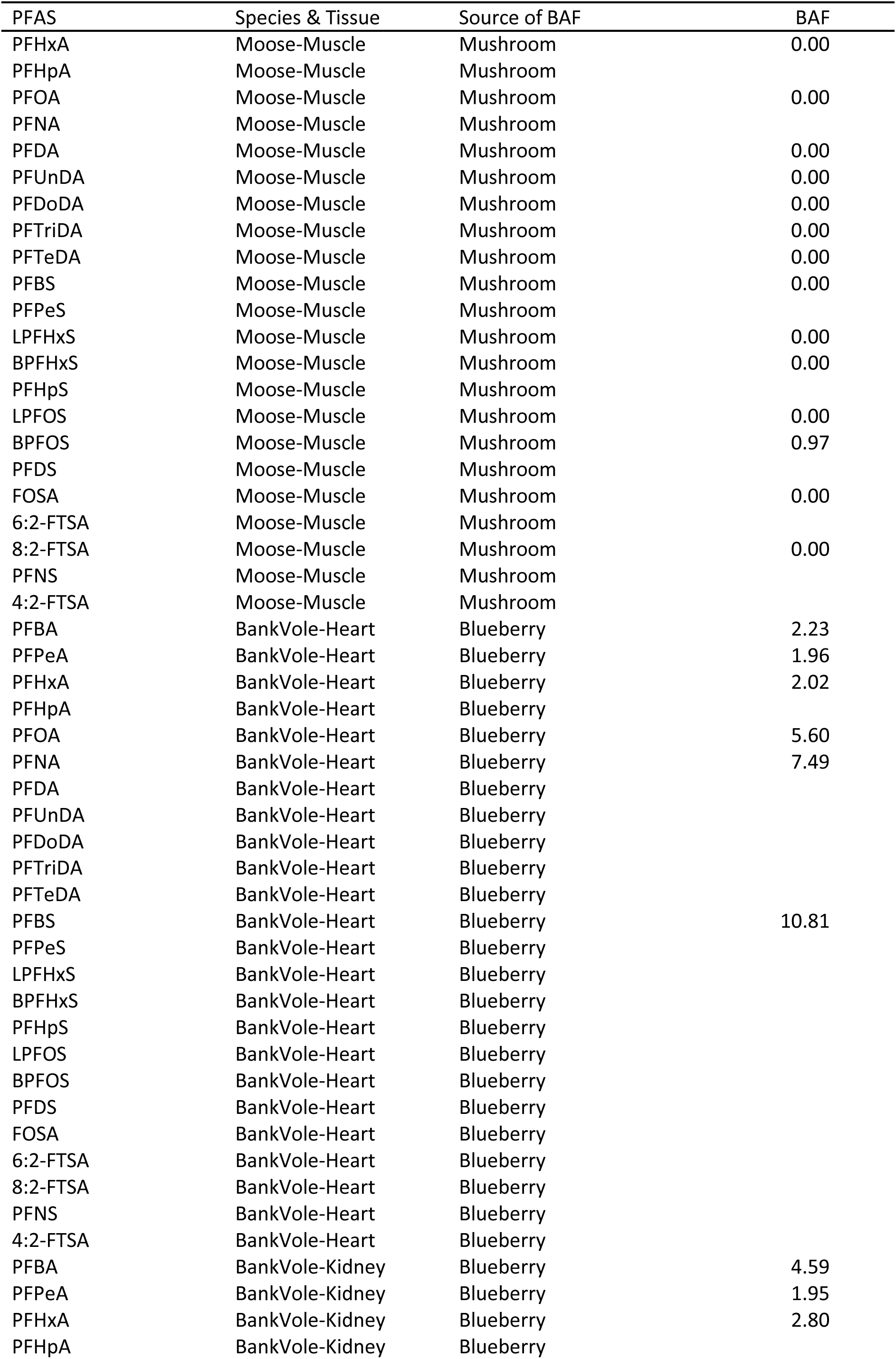
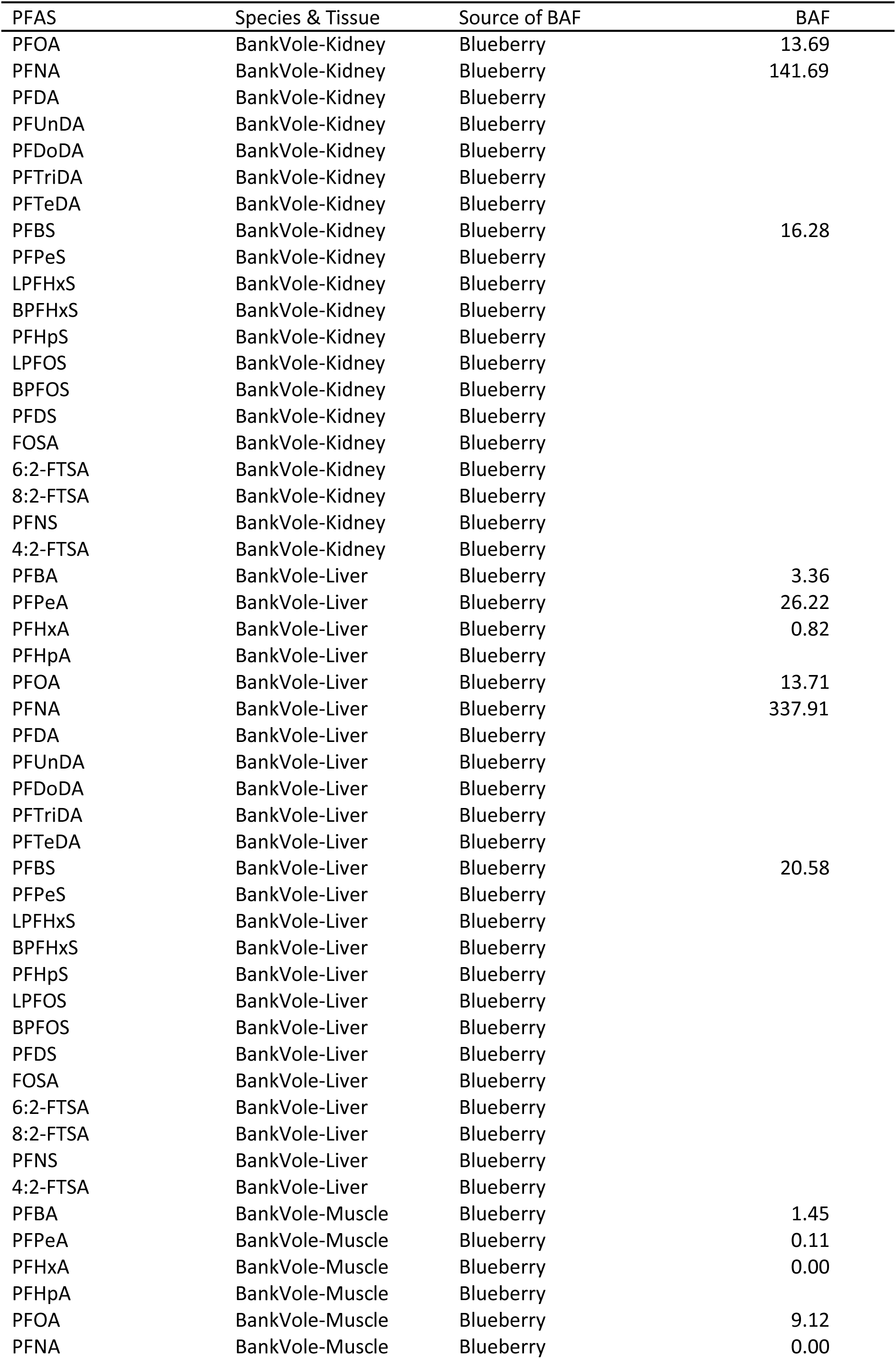
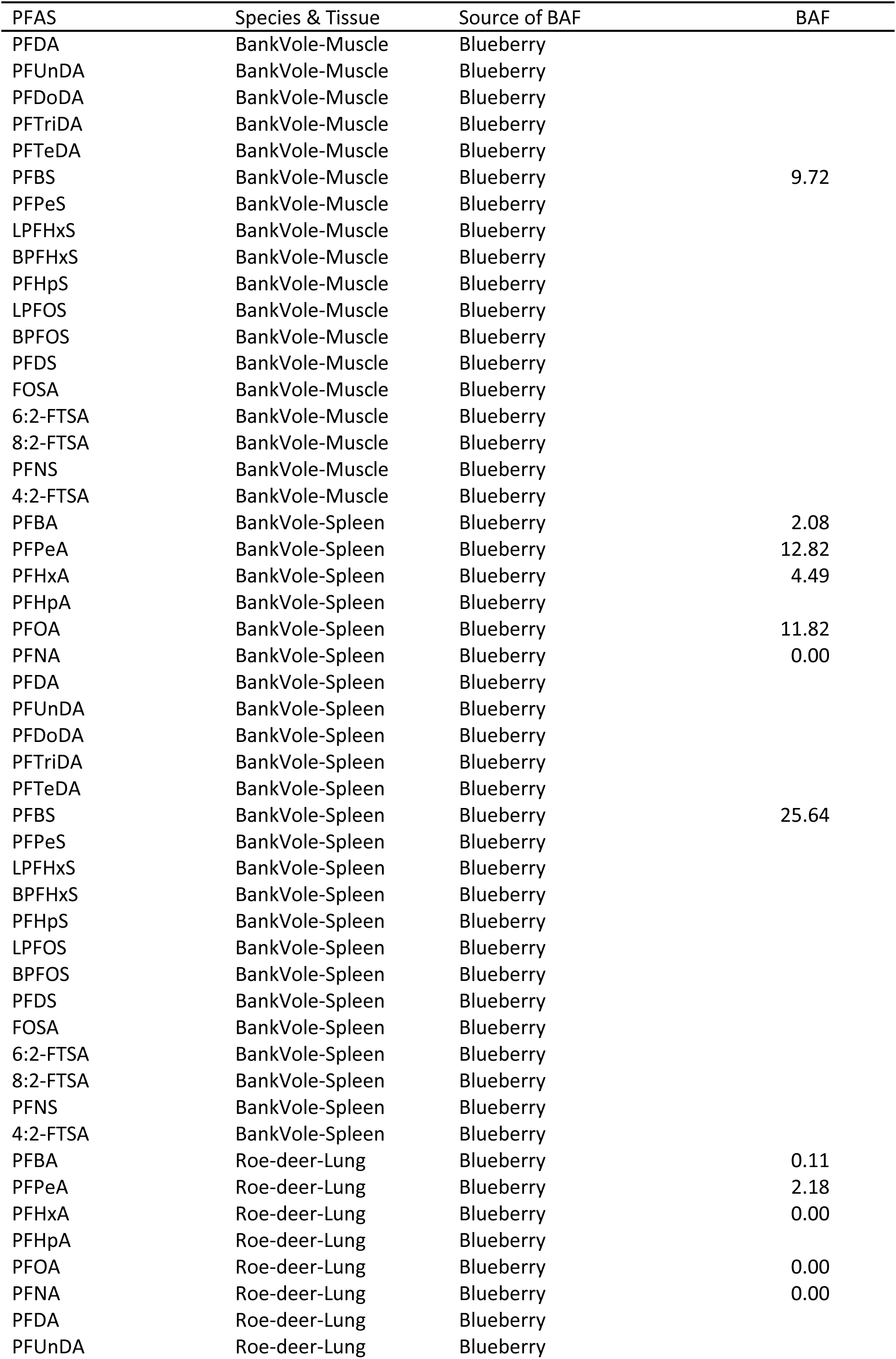
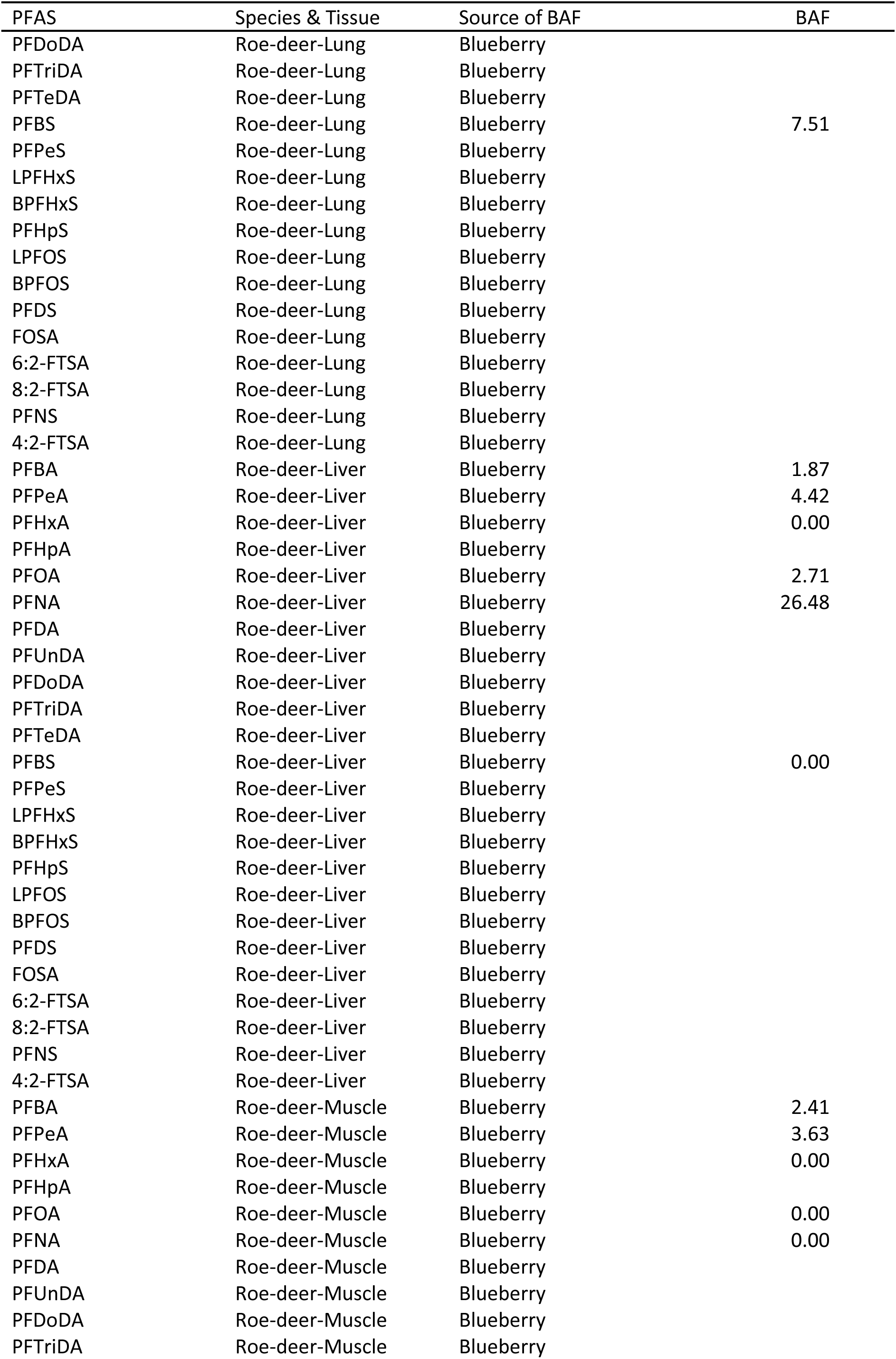
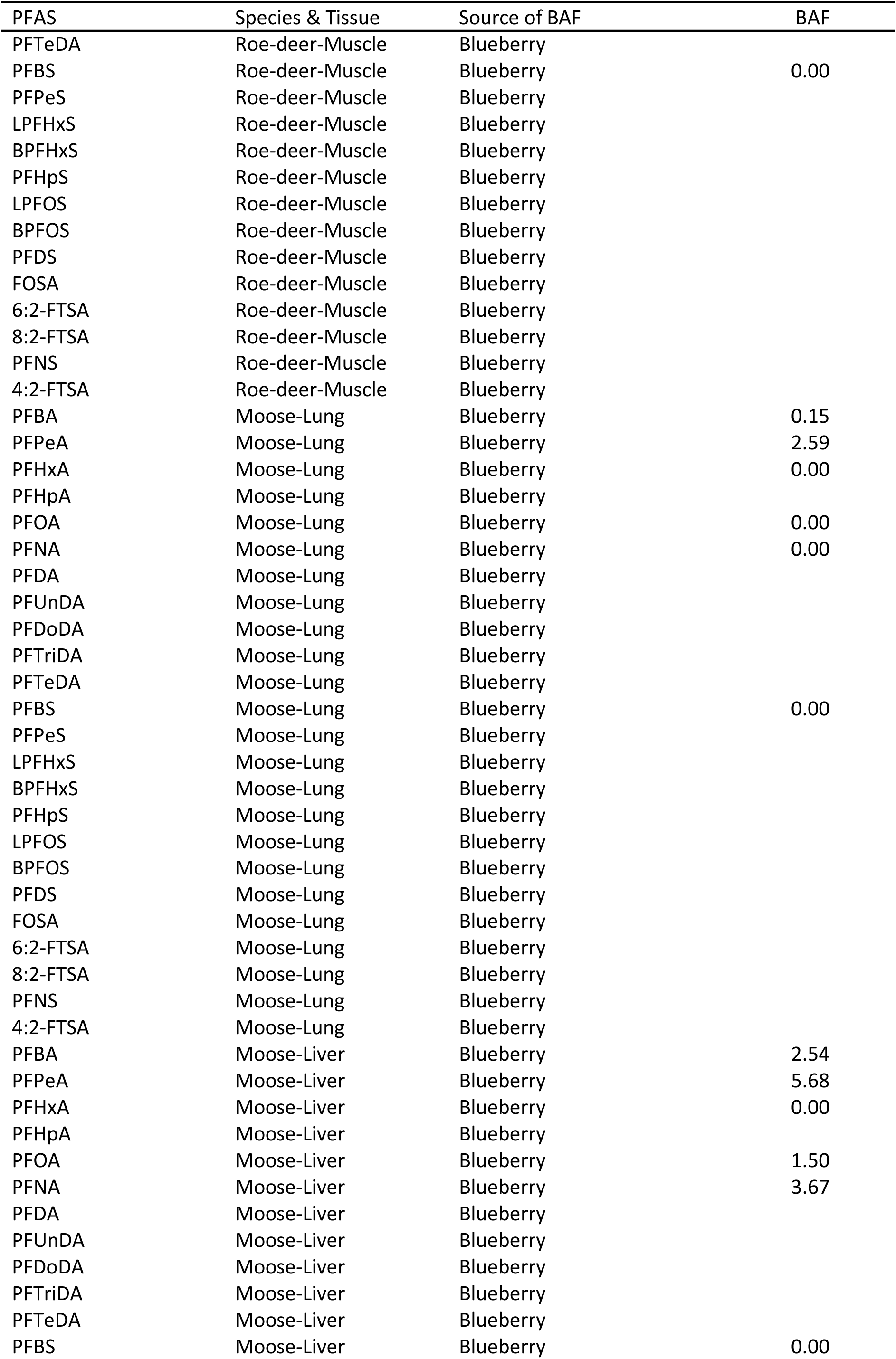
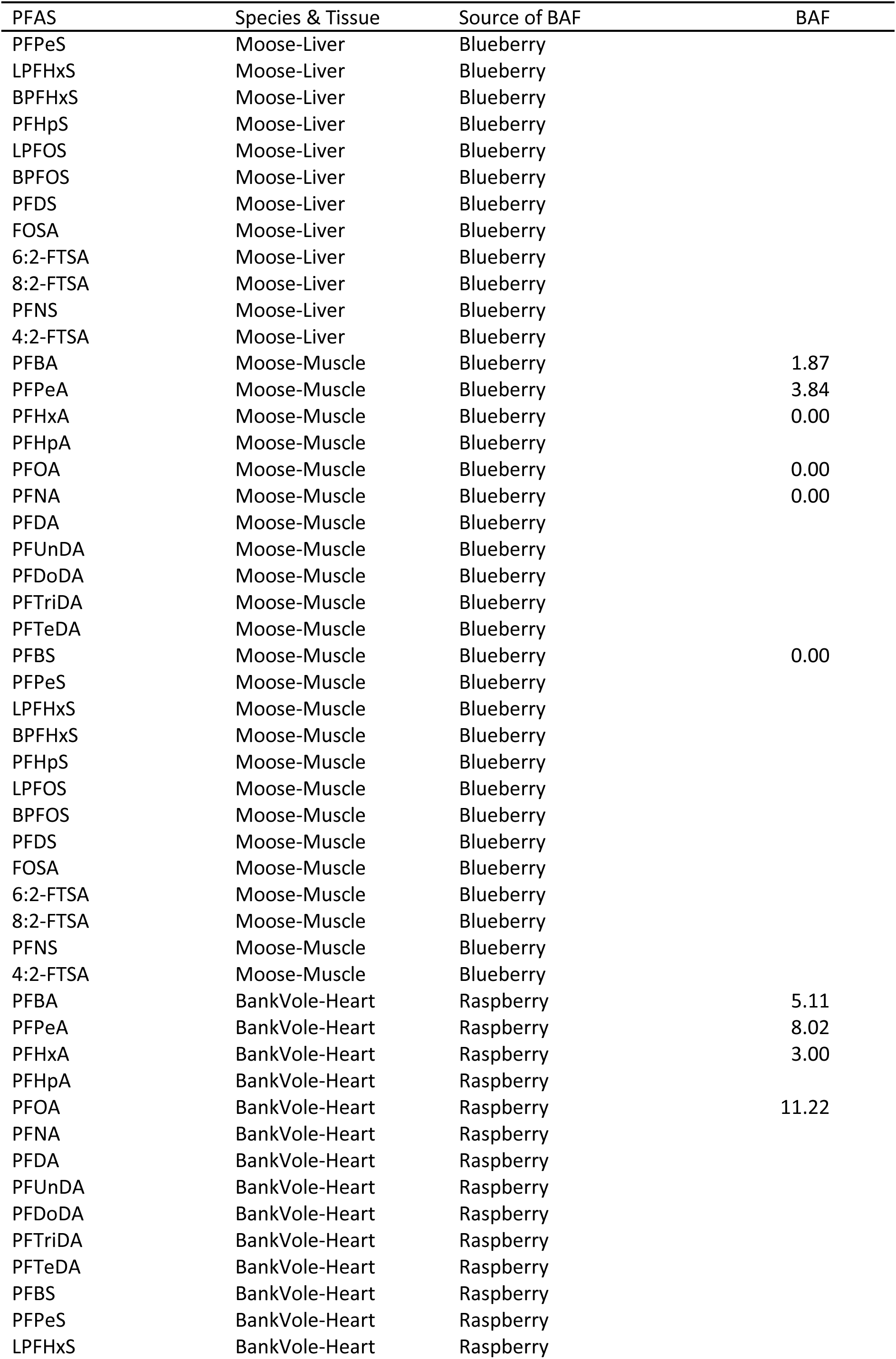
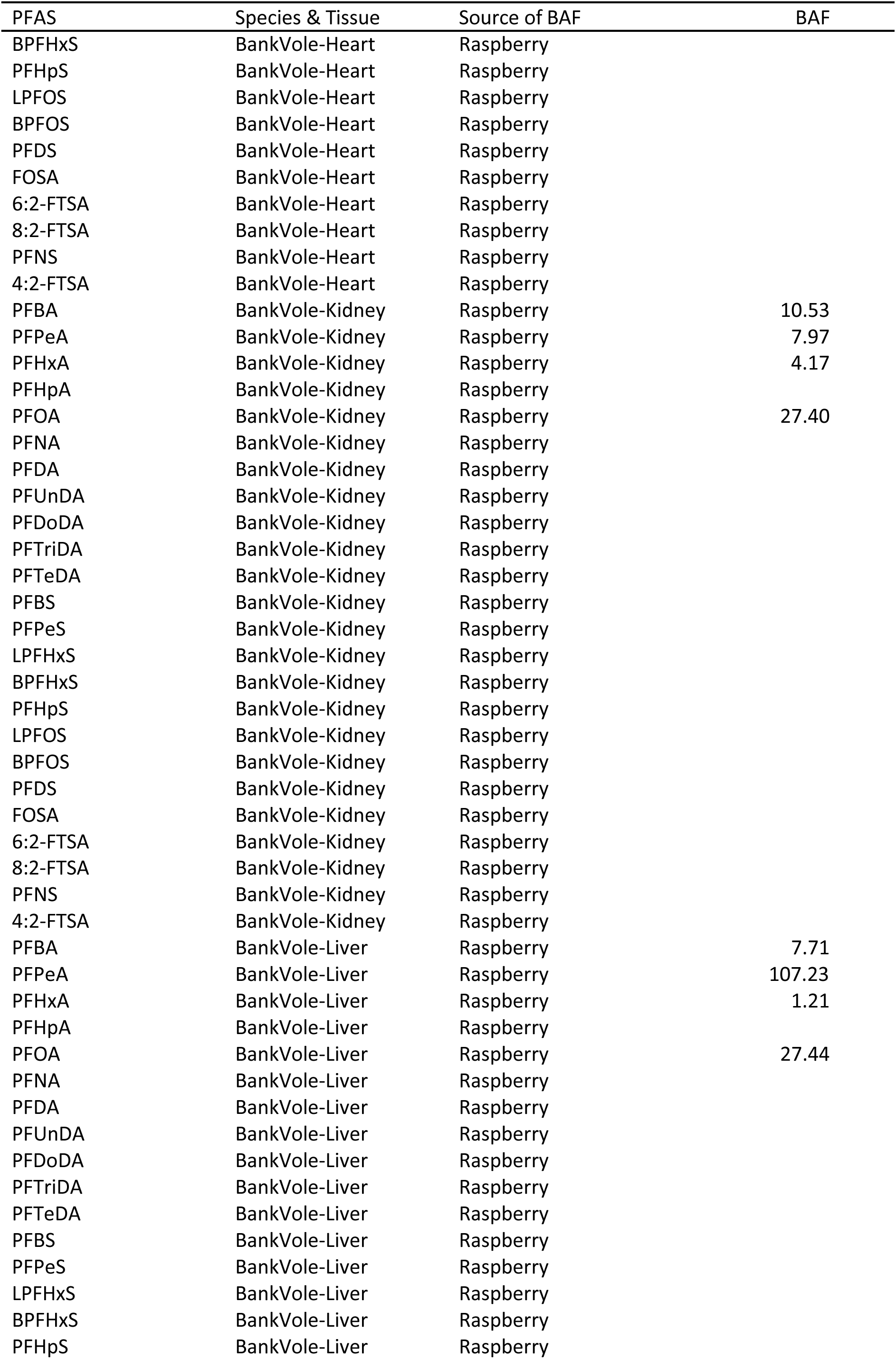
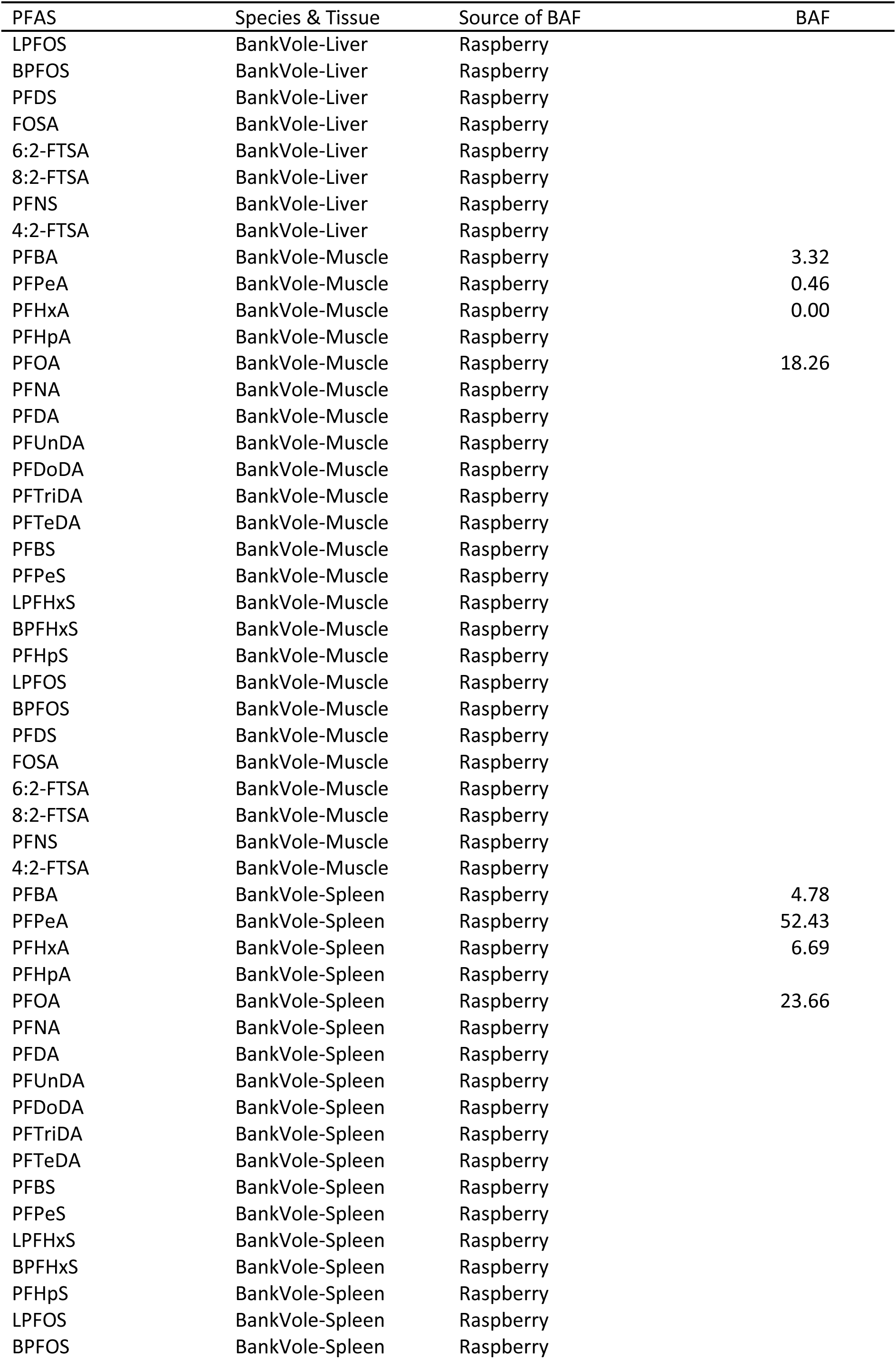
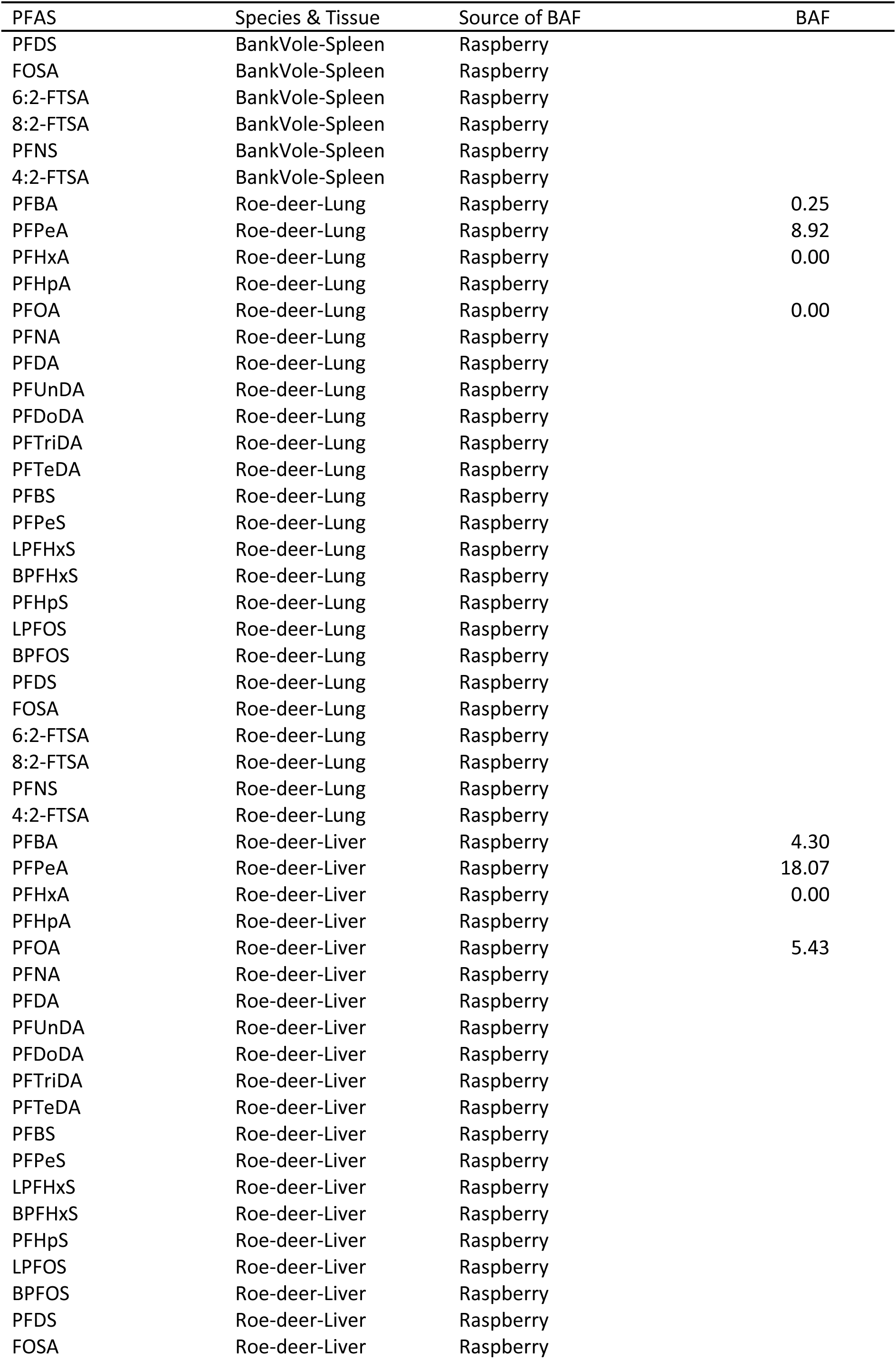
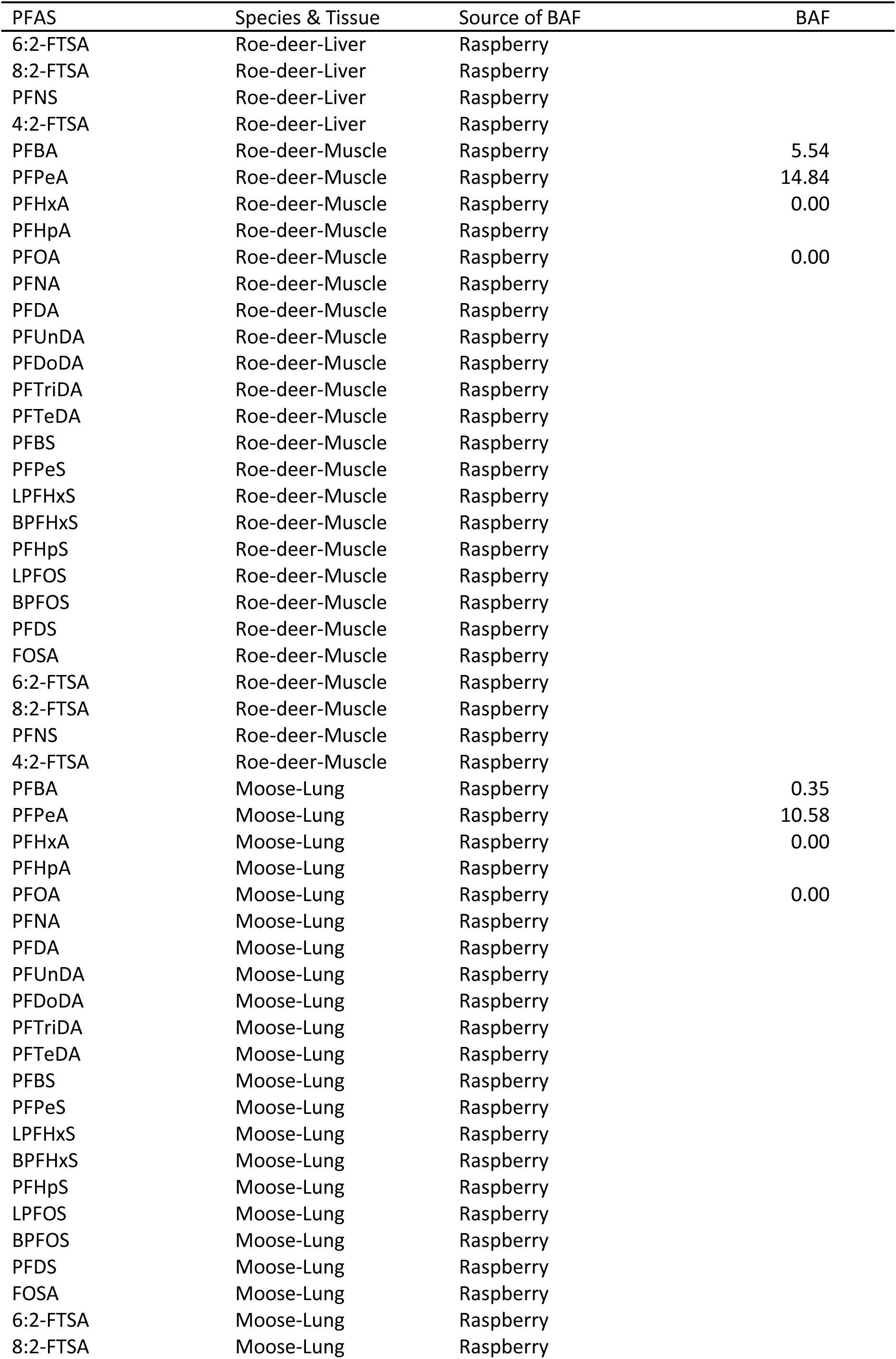
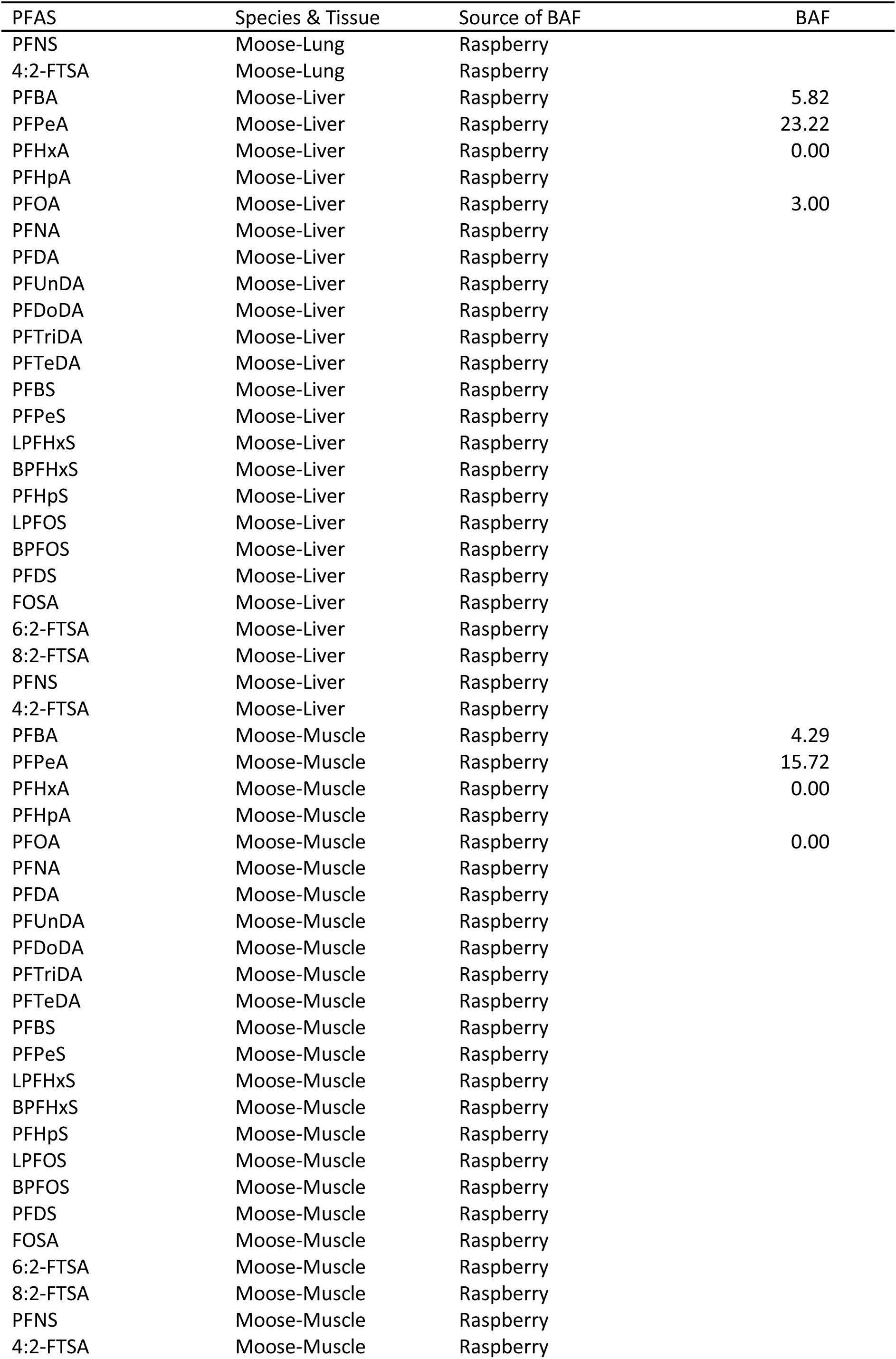
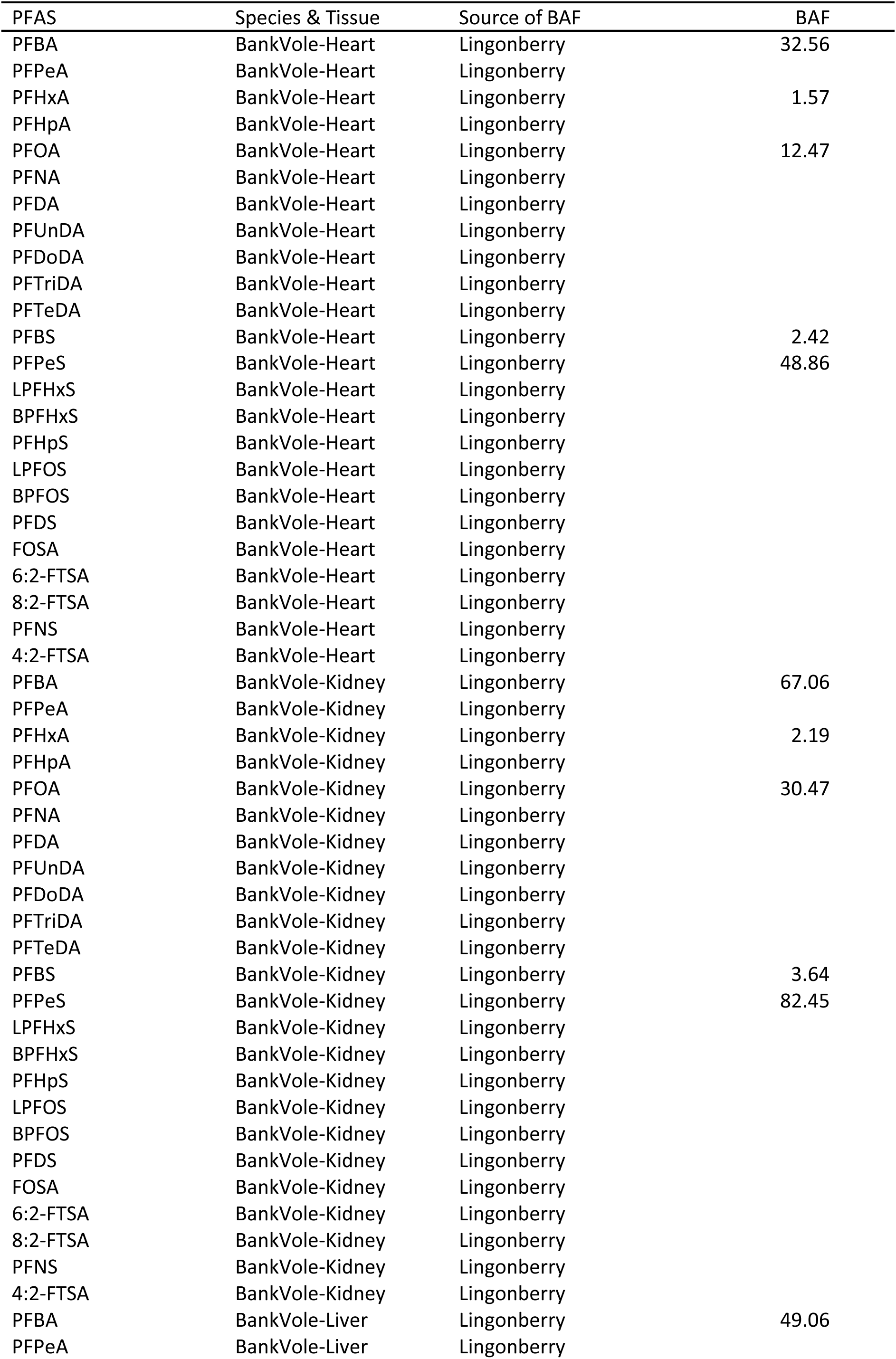
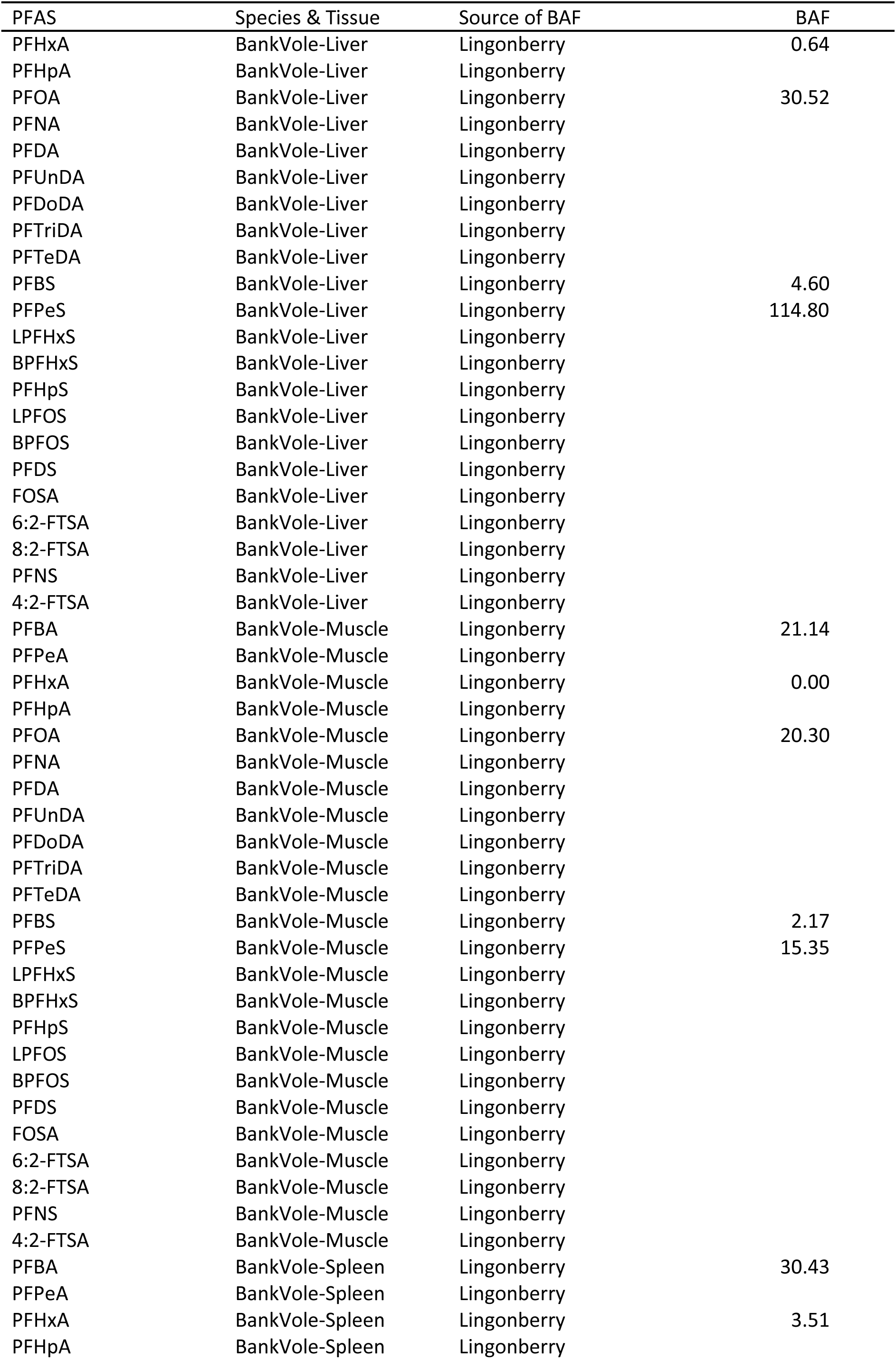
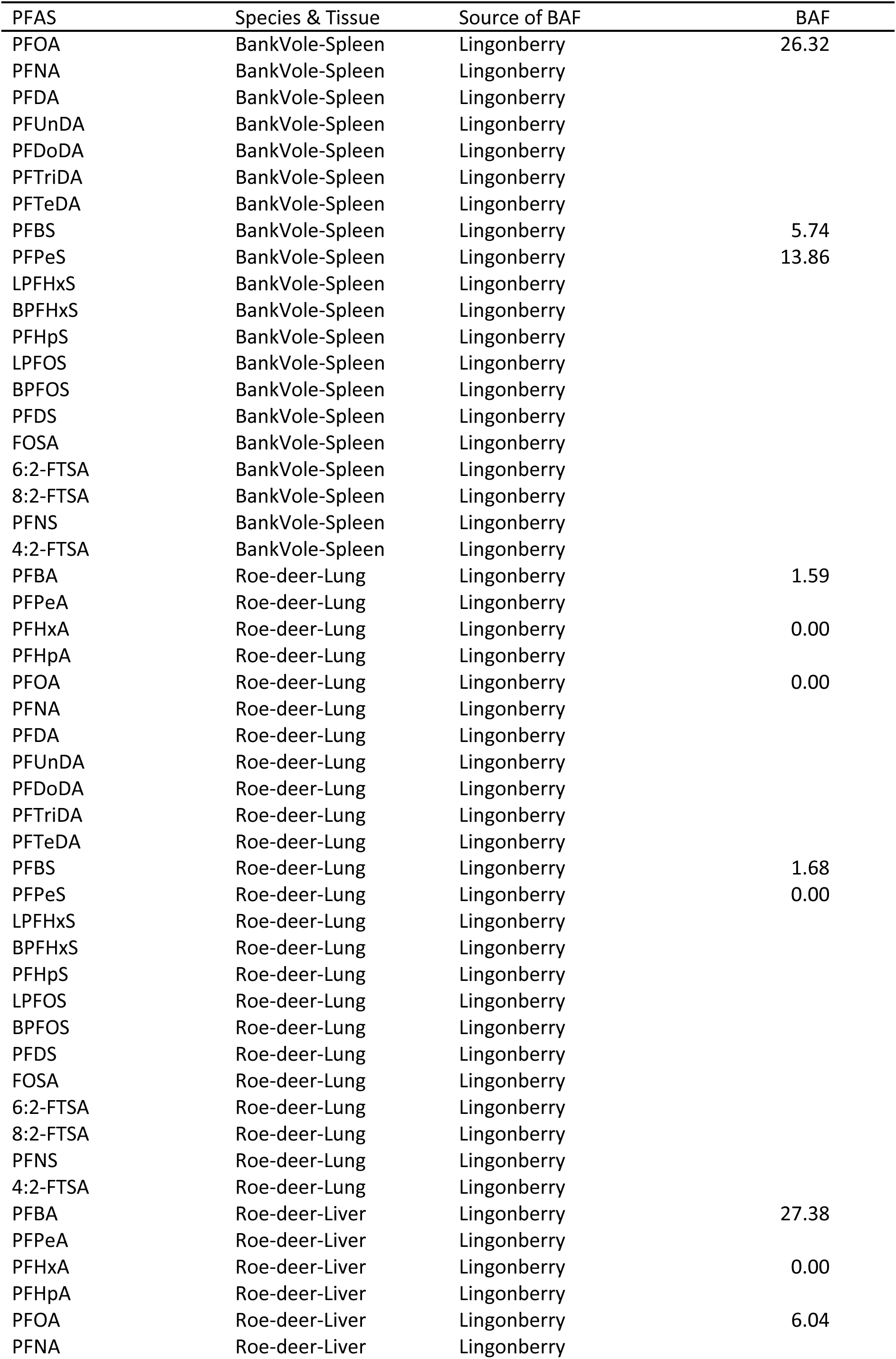
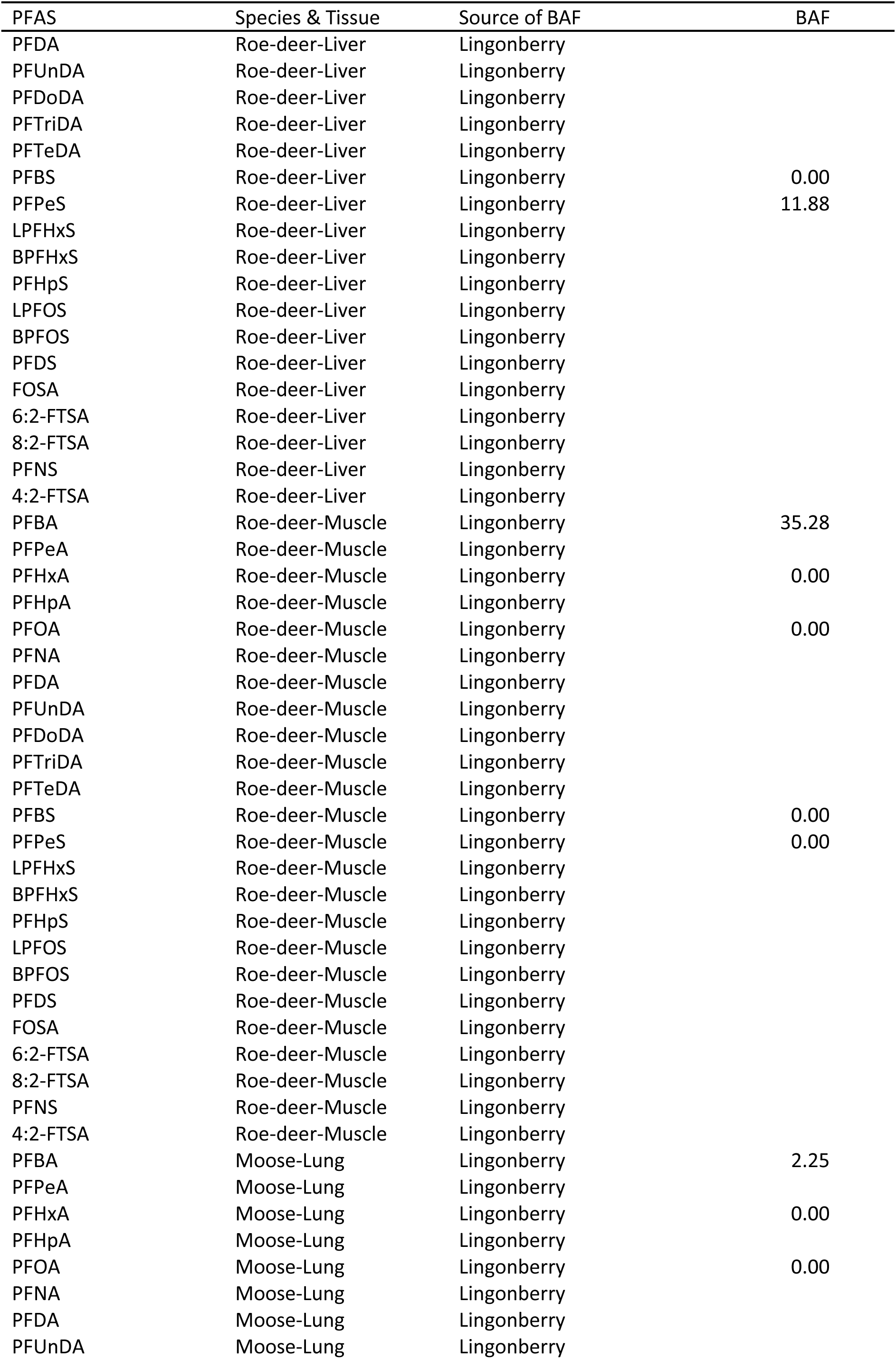
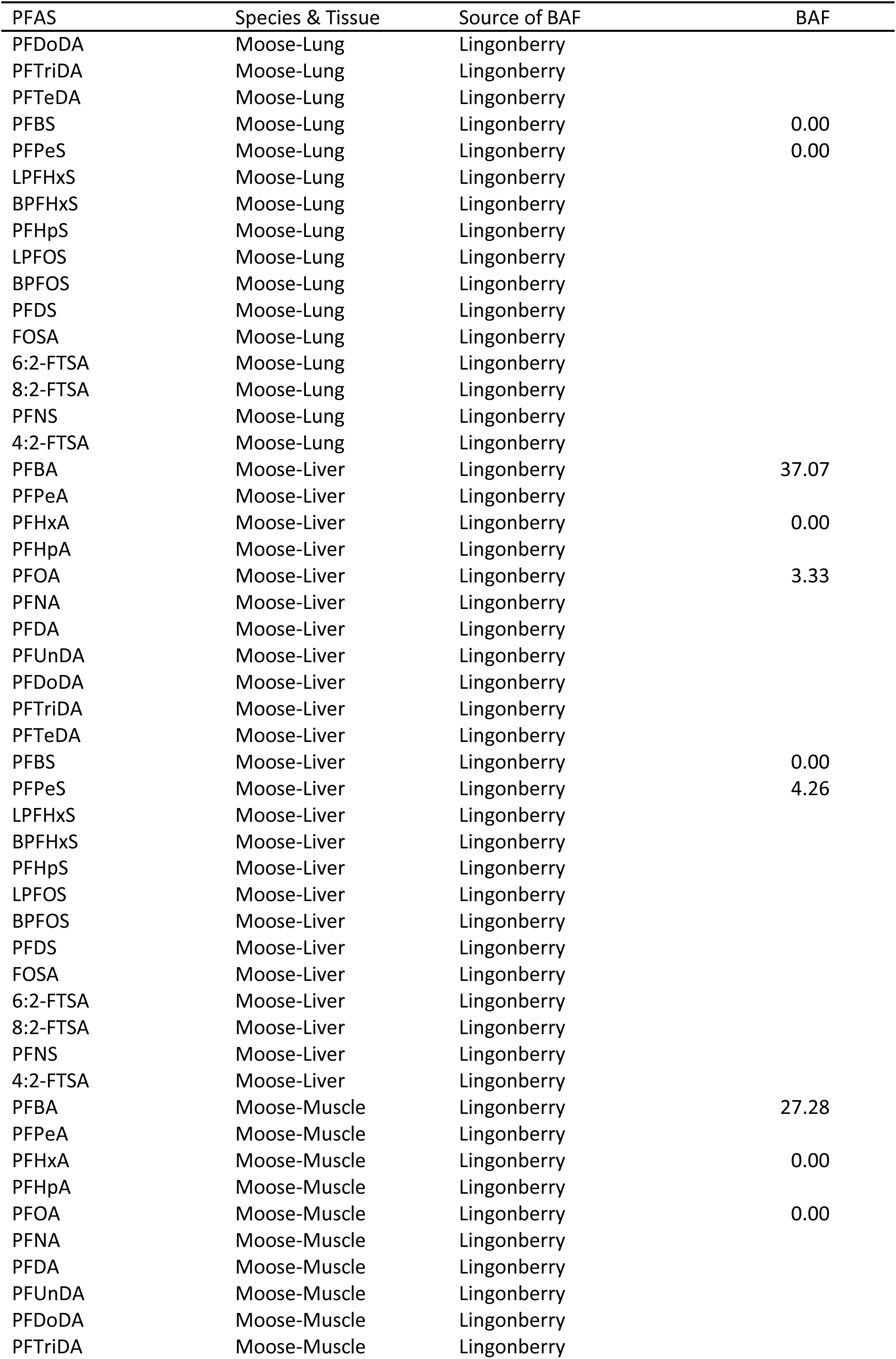
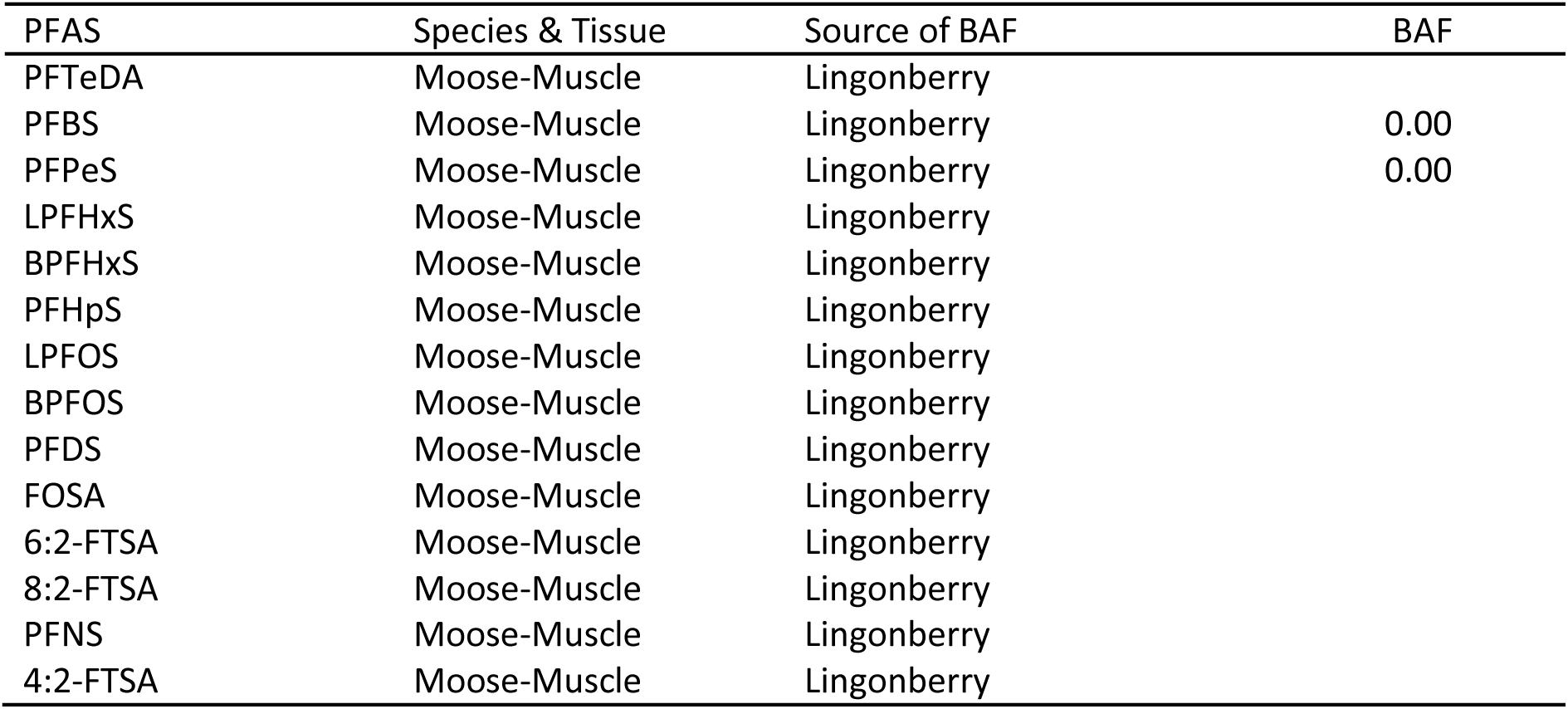
Bioaccumulation factors (BAF) for PFAS in different species and tissues. Ref refers to the reference areas near Umeå.

## Footnotes

1 https://www.efsa.europa.eu/en/news/pfas-food-efsa-assesses-risks-and-sets-tolerable-intake

## References

1. Ahrens L, Bundschuh M. Fate and effects of poly– and perfluoroalkyl substances in the aquatic environment: A review. Environmental Toxicology and Chemistry 2014; 33: 1921–1929.

2. Axelsson M, Bard J. MTU avseende PFAS inom f.d. Jämtlands flygflottilj, F 4 Frösön Linköping: NIRAS Sweden AB, 2015.

3. Beans C. How “forever chemicals” might impair the immune system. Proceedings of the National Academy of Sciences 2021; 118: e2105018118.

4. Bergstedt B. Home ranges and movements of the rodent species *Clethrionomys glareolus* (Schreber), *Apodemus flavicollis* (Melchior) and *Apodemus sylvaticus* (Linné) in southern Sweden. Oikos 1966; 17: 150–157.

5. Burkhard LP, Votava LK. Review of per– and polyfluoroalkyl substances (PFAS) bioaccumulation in earthworms. Environmental Advances 2023; 11: 100335.

6. Bustnes JO, Bangjord G, Ahrens L, Herzke D, Yoccoz NG. Perfluoroalkyl substance concentrations in a terrestrial raptor: Relationships to environmental conditions and individual traits. Environmental Toxicology and Chemistry 2015; 34: 184–191.

7. Cederlund G, Ljungqvist H, Markgren G, Stalfelt F. Foods of moose and roe-deer at Grimsö in central Sweden-results of rumen content analyses. Swedish Wildlife Research 1980; 11: 171–247.

8. Chelcea IC, Ahrens L, Örn S, Mucs D, Andersson PL. Investigating the OECD database of per-and polyfluoroalkyl substances–chemical variation and applicability of current fate models. Environmental Chemistry 2020; 17: 498–508.

9. Costello E, Rock S, Stratakis N, Eckel SP, Walker DI, Valvi D, et al. Exposure to per– and Polyfluoroalkyl Substances and Markers of Liver Injury: A Systematic Review and Meta-Analysis. Environ Health Perspect 2022; 130: 46001.

10. Dickman RA, Aga DS. A review of recent studies on toxicity, sequestration, and degradation of per– and polyfluoroalkyl substances (PFAS). Journal of Hazardous Materials 2022; 436: 129120.

11. Ecke F, Benskin JP, Berglund ÅMM, de Wit CA, Engström E, Plassmann MM, et al. Spatio-temporal variation of metals and organic contaminants in bank voles (Myodes glareolus). Science of The Total Environment 2020; 713: 136353.

12. Ecke F, Berglund ÅMM, Rodushkin I, Engström E, Pallavicini N, Sörlin D, et al. Seasonal shift of diet in bank voles explains trophic fate of anthropogenic osmium? Science of The Total Environment 2018; 624: 1634–1639.

13. Ericson L. The influence of voles and lemmings on the vegetation in a coniferous forest during a 4– year period in northern Sweden. Wahlenbergia 1977; 4: 1-114.

14. European Commission. COMMISSION REGULATION (EU) 2022/2388 of 7 December 2022 amending Regulation (EC) No 1881/2006 as regards maximum levels of perfluoroalkyl substances in certain foodstuffs 2022.

15. Evich MG, Davis MJB, McCord JP, Acrey B, Awkerman JA, Knappe DRU, et al. Per– and polyfluoroalkyl substances in the environment. Science 2022; 375: eabg9065.

16. Falk S, Brunn H, Schröter-Kermani C, Failing K, Georgii S, Tarricone K, et al. Temporal and spatial trends of perfluoroalkyl substances in liver of roe deer (*Capreolus capreolus*). Environmental Pollution 2012; 171: 1–8.

17. Golovko O, Kaczmarek M, Asp H, Bergstrand K-J, Ahrens L, Hultberg M. Uptake of perfluoroalkyl substances, pharmaceuticals, and parabens by oyster mushrooms (*Pleurotus ostreatus*) and exposure risk in human consumption. Chemosphere 2022; 291: 132898.

18. Grønnestad R, Vázquez BP, Arukwe A, Jaspers VLB, Jenssen BM, Karimi M, et al. Levels, Patterns, and Biomagnification Potential of Perfluoroalkyl Substances in a Terrestrial Food Chain in a Nordic Skiing Area. Environmental Science & Technology 2019; 53: 13390–13397.

19. Hansson L. Small Rodent Food, Feeding and Population Dynamics: A Comparison between Granivorous and Herbivorous Species in Scandinavia. Oikos 1971; 22: 183–198.

20. Hansson L. Condition and diet in relation to habitat in bank voles *Clethrionomys glareolus*: Population or community approach? Oikos 1979a; 33: 55–63.

21. Hansson L. Food as a limiting factor for small rodent numbers. Oecologia 1979b; 37: 297–314.

22. Hansson L. *Clethrionomys* food; generic, specific and regional characteristics. Annales Zoologici Fennici 1985a; 22: 315–318.

23. Hansson L. The food of bank voles, wood mice and yellow-necked mice. In: Flowerdew JR, Gurnell J, Gipps JHW, editors. The ecology of woodland rodents: bank voles and wood mice. 55. Symposia of the Zoological Society of London, Oxford, 1985b, pp. 141–168.

24. Hansson L. Grazing impact by small rodents in a steep cyclicity gradient. Oikos 1988; 51: 31–42.

25. Hipkiss T, Hörnfeldt B. High interannual variation in the hatching sex ratio of Tengmalm’s owl broods during a vole cycle. Population Ecology 2004; 46: 263–268.

26. Hipkiss T, Stefansson O, Hornfeldt B. Effect of cyclic and declining food supply on great grey owls in boreal Sweden. Canadian Journal of Zoology 2008; 86: 1426–1431.

27. Hörnfeldt B, Carlsson BG, Löfgren O, Eklund U. Effects of cyclic food supply on breeding performance in Tengmalm’s owl. Canadian Journal of Zoology 1990; 68: 522–530.

28. Jones KE, Bielby J, Cardillo M, Fritz SA, O’Dell J, Orme CDL, et al. PanTHERIA: a species-level database of life history, ecology, and geography of extant and recently extinct mammals. Ecology 2009; 90: 2648–2648.

29. Jongman RHG, ter Braak CJF, van Tongeren OFR. Data analysis in community and landscape ecology. Cambridge: Cambridge University Press, 1995.

30. Khalil H, Ecke F, Evander M, Bucht G, Hörnfeldt B. Population dynamics of bank voles predict human Puumala hantavirus risk. EcoHealth 2019; 16: 545–555.

31. Khalil H, Olsson G, Ecke F, Evander M, Hjertqvist M, Magnusson M, et al. The importance of bank vole density and rainy winters in predicting nephropathia epidemica incidence in Northern Sweden. Plos One 2014; 9: e111663.

32. Koch A, Kärrman A, Yeung LW, Jonsson M, Ahrens L, Wang T. Point source characterization of per– and polyfluoroalkyl substances (PFASs) and extractable organofluorine (EOF) in freshwater and aquatic invertebrates. Environmental Science: Processes & Impacts 2019; 21: 1887–1898.

33. Kouba M, Bartoš L, Tomášek V, Popelková A, Šťastný K, Zárybnická M. Home range size of Tengmalm’s owl during breeding in Central Europe is determined by prey abundance. PLOS ONE 2017; 12: e0177314.

34. Krebs CJ, Myers JH. Population cycles in small mammals. Advances in Ecoligical Research 1974; 8: 267–399.

35. Kurwadkar S, Dane J, Kanel SR, Nadagouda MN, Cawdrey RW, Ambade B, et al. Per-and polyfluoroalkyl substances in water and wastewater: A critical review of their global occurrence and distribution. Science of The Total Environment 2022; 809: 151003.

36. Löfgren O. Spatial organization of cyclic Clethrionomys females: occupancy of all available space at peak densities? Oikos 1995; 72: 29–35.

37. Magnusson M, Samelius G, Hörnfeldt B, Ecke F. Diet shift in bank voles induced by competition from grey-sided voles? Integrative Zoology 2019; 14: 376–382.

38. Mitchell-Jones AJ, Amori G, Bogdanowicz W, Kruŝtufek B, Reijnders PJH, Spitzenberger F, et al. The atlas of European mammals. London: Poyser Natural History, 1999.

39. Modin H. PFAS i Östersunds kommun, 2021. Report Number: 00073–2021 i databas MSN.

40. Nassazzi W, Lai FY, Ahrens L. A novel method for extraction, clean-up and analysis of per-and polyfluoroalkyl substances (PFAS) in different plant matrices using LC-MS/MS. Journal of Chromatography B 2022; 1212: 123514.

41. R Development Core Team. R: a language and environment for statistical computing. Version 4.2.0. Foundation for Statistical Computing, Vienna, Austria, Vienna, Austria, 2021.

42. Roos AM, Gamberg M, Muir D, Kärrman A, Carlsson P, Cuyler C, et al. Perfluoroalkyl substances in circum-ArcticRangifer: caribou and reindeer. Environmental Science and Pollution Research 2022; 29: 23721–23735.

43. Smithwick M, Muir DC, Mabury SA, Solomon KR, Martin JW, Sonne C, et al. Perfluoroalkyl contaminants in liver tissue from East Greenland polar bears (*Ursus maritimus*). Environmental Toxicology and Chemistry: An International Journal 2005; 24: 981–986.

44. Spitzer R, Felton A, Landman M, Singh NJ, Widemo F, Cromsigt JP. Fifty years of European ungulate dietary studies: a synthesis. Oikos 2020; 129: 1668–1680.

45. Vapalahti O, Mustonen J, Lundkvist A, Henttonen H, Plyusnin A, Vaheri A. Hantavirus infections in Europe. Lancet Infectious Diseases 2003; 3: 753–754.

